# Multi-resolution localization of causal variants across the genome

**DOI:** 10.1101/631390

**Authors:** Matteo Sesia, Eugene Katsevich, Stephen Bates, Emmanuel Candès, Chiara Sabatti

## Abstract

We present *KnockoffZoom*, a flexible method for the genetic mapping of complex traits at multiple resolutions. *KnockoffZoom* localizes causal variants by testing the conditional associations of genetic segments of decreasing width while provably controlling the false discovery rate using artificial genotypes as negative controls. Our method is equally valid for quantitative and binary phenotypes, making no assumptions about their genetic architectures. Instead, we rely on well-established genetic models of linkage disequilibrium. We demonstrate that our method can detect more associations than mixed effects models and achieve fine-mapping precision, at comparable computational cost. Lastly, we apply *KnockoffZoom* to data from 350k subjects in the UK Biobank and report many new findings.

## I. INTRODUCTION

Since the sequencing of the human genome, there have been massive efforts to identify genetic variants that affect phenotypes of medical relevance. In particular, single-nucleotide polymorphisms (SNPs) have been genotyped in large cohorts in the context of genome-wide association studies (GWAS), leading to the discovery of thousands of associations with different traits and diseases.^1^ However, it has been challenging to translate these findings into actionable knowledge.^2^ As a first step in this direction, we present a new statistical method for the genetic mapping of complex phenotypes that improves our ability to resolve the location of causal variants.

The analysis of GWAS data initially proceeded SNP-by-SNP, testing *marginal* independence with the trait (not accounting for the rest of the genome) using univariate regression. Today the leading solutions rely on a linear mixed model (LMM) that differs from the previous method insofar as it includes a random term approximating the effects of other variants in distant loci.^3–8^ Nonetheless, one can still think of the LMM as testing marginal independence because it does not account for linkage disequilibrium (LD).^9,10^ Thus, it cannot distinguish causal SNPs from nearby variants that may have no interesting biological function, since neither are independent of the phenotype. This limitation becomes concerning as we increasingly focus on polygenic traits and rely on large samples; in this setting it has been observed that most of the genome is correlated with the phenotype, even though only a fraction of the variants may be important.^11^ Therefore, the null hypotheses of no association should be rejected for most SNPs,^2^ which is a rather uninformative conclusion. The awareness among geneticists that marginal testing is insufficient has led to the heuristic practice of post-hoc aggregation (*clumping*) of associated loci in LD^8,12^ and to the development of fine-mapping.^13^ Fine-mapping methods are designed to refine marginally significant loci and discard associated but non-causal SNPs by accounting for LD, often within a Bayesian perspective.^14–17^ However, this two-step approach is not fully satisfactory because it requires switching models and assumptions in the middle of the analysis, obfuscating the interpretation of the findings and possibly invalidating type-I error guarantees. Moreover, as LD makes more non-causal variants appear marginally associated with the trait in larger samples, the standard fine-mapping tools face an increasingly complicated task refining wider regions.

*KnockoffZoom* simultaneously addresses the current difficulties in locus discovery and fine-mapping by searching for causal variants over the entire genome and reporting those SNPs (or groups thereof) that appear to have a distinct influence on the trait while accounting for the effects of all others. This search is carried out by testing the conditional association of pre-defined groups at multiple resolutions, ranging from that of locus discovery to that of fine-mapping. In this process, we precisely control the false discovery rate (FDR)^18^ and obtain findings that are directly interpretable, without heuristic post-processing. Our inferences rest on the assumption that LD is adequately described by hidden Markov models (HMMs) that have been successfully employed in many areas of genetics.^19–22^ We do not require any model linking genotypes to phenotypes; therefore, our approach seamlessly applies to both quantitative and qualitative traits.

This work is facilitated by recent advances in statistics, notably *knockoffs*,^23^ whose general validity for GWAS has been explored and discussed before.^24–29^ We leverage these results and introduce several key innovations. First, we develop new algorithms to analyze the data at multiple levels of resolution, in such a way as to maximize power in the presence of LD without pruning the variants.^23,24^ Second, we improve the computational efficiency and apply our method to a large dataset, the UK Biobank^30^—a previously computationally unfeasible task.

## II. RESULTS

### A. KnockoffZoom

To localize causal variants as precisely as possible and provide researchers with interpretable results, *KnockoffZoom* tests *conditional hypotheses* (Online Methods A). In the simplest GWAS analysis, a variant is null if the distribution of its alleles is independent of the phenotype. By contrast, our hypotheses are much stricter: a variant is null if it is independent of the trait *conditionally* on all other variants, including its neighbors. Suppose that we believed, consistently with the classical polygenic literature,^31^ that a multivariate linear model realistically describes the genetic makeup of the trait; then, a conditional hypothesis would be non-null if the corresponding variant had a non-zero coefficient. In general, in the absence of unmeasured confounders (Sections II B–II C) or any feedback effects of the phenotype onto the genome, our tests lead to the discovery of causal variants. In particular, we can separate markers that have a distinct effect on the phenotype from those whose associations are merely due to LD.

The presence of LD makes the conditional null hypotheses challenging to reject, especially with small sample sizes. This is why *t*-tests for multivariate linear regression may have little power in the presence of collinearity; in this case, *F* -tests provide a possible solution. Analogously, we group variants that are too similar for their distinct effects to be discerned. More precisely, we test whether the trait is independent of all SNPs in an LD block, conditional on the others. A block is null if it only contains SNPs that are independent of the phenotype, conditional on the other blocks. Concretely, we define contiguous blocks at multiple resolutions by partitioning the genome via adjacency-constrained hierarchical clustering.^32^ We adopt the *r*^2^ coefficients computed by PLINK^12^ as a similarity measure for the SNPs and cut the dendrogram at different heights; see Figure 1 (b).

However, any other choice of groups could be easily integrated into *KnockoffZoom* as long as the groups are determined before looking at the phenotype data. In particular, we note that contiguity is not necessary; we have adopted it because it corresponds well to the idea that researchers are attempting to localize causal variants within a genomic segment, and because it produces results that are easily interpretable when some of the genetic variation is not genotyped (Section II C). Additionally, when assuming an HMM for the distribution of the genotypes, contiguous groups of SNPs tend to lead to higher power, as explained in Supplementary Section S2 A 1.

**FIG. 1.**
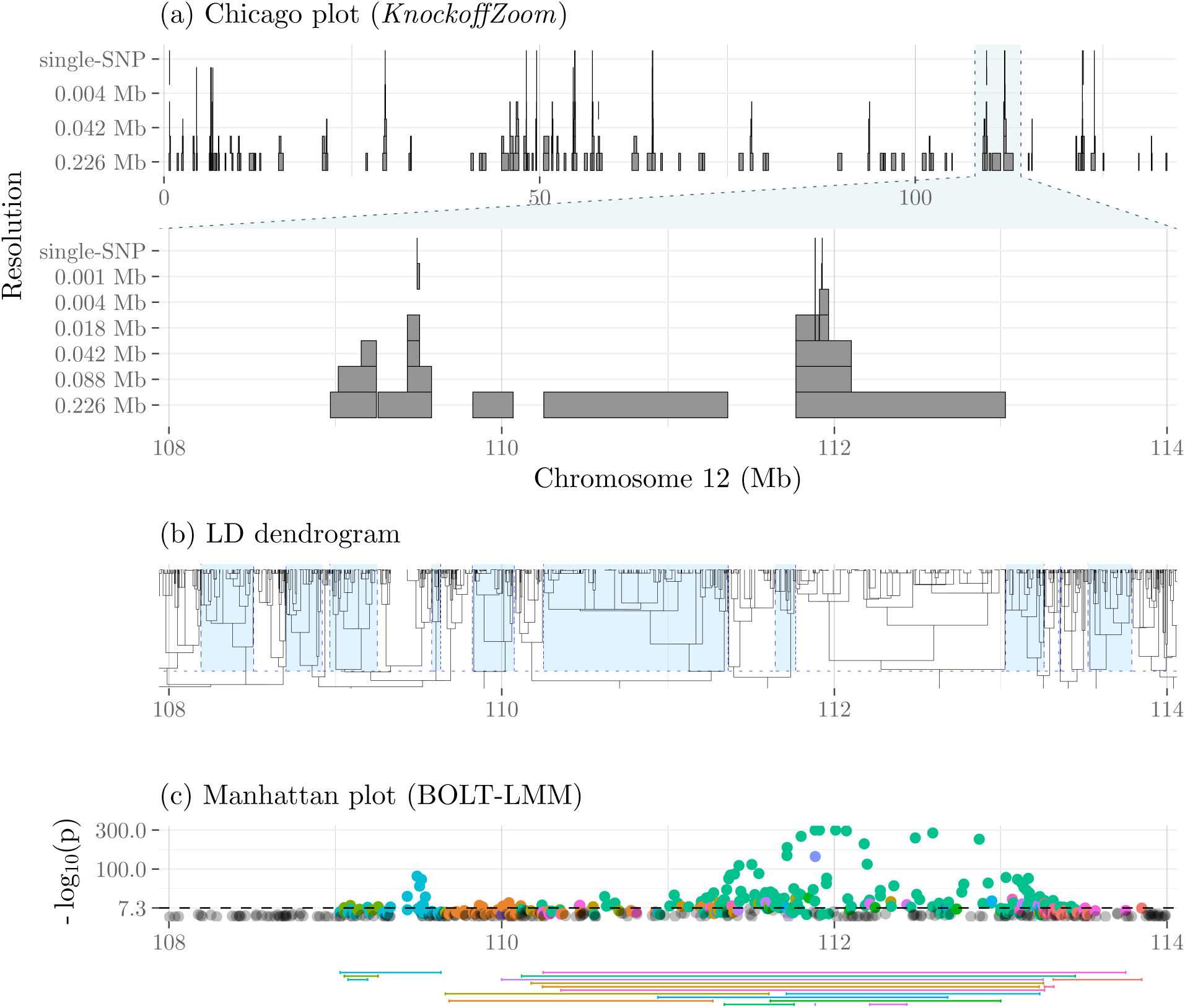
*KnockoffZoom* discoveries (a) on chromosome 12 for the phenotype *platelet* in the UK Biobank, controlling the FDR below 0.1. Each shaded rectangle represents a discovery at the resolution indicated by its vertical position (measured by the average width of the blocks), so that the highest-resolution findings are on top. The lower part of (a) focuses on a smaller genomic region. The hypotheses are pre-specified by cutting the LD dendrogram (b) at different heights. As an example, by alternating blue and white shading in (b), we indicate the lowest-resolution blocks. The Manhattan plot (c) shows the BOLT-LMM p-values from the same data, while the segments below represent the region spanned by the significant discoveries clumped with PLINK at the genome-wide significance level (5 × 10^−8^). For each clump, the colors match those of the corresponding p-values. All plots are vertically aligned, except for the top part of (a).

To balance power and resolution, we consider multiple partitions, starting with a coarse view and successively refining it (Supplementary Table S3). An example of our results for *platelet* is shown in Figure 1 (a). Each rectangle in the *Chicago* plot (named for its stylistic resemblance to the Willis tower) spans a region that includes variants carrying distinct information about the trait compared to the other blocks on the same level. The blocks at higher levels correspond to higher resolutions; these are narrower and more difficult to discover, since only strong associations can be refined precisely.

We can mathematically prove that *KnockoffZoom* controls the FDR below any desired level *q* (at most a fraction *q* of our findings are false positives on average), at each resolution. The FDR is a meaningful error rate for the study of complex traits,^33,34^ but its control is challenging.^35^ We overcome this difficulty with *knockoffs*. These are carefully engineered synthetic variables that can be used as negative controls because they are exchangeable with the genotypes and reproduce the spurious associations that we want to winnow.^23,24,36,37^ The construction of knockoffs requires specifying the distribution of the genotypes, which we approximate as an HMM. In this paper, we implement the HMM of fastPHASE,^20^ (Supplementary Section S2 B), which we show works well for the genotypes of relatively homogeneous individuals, even though it has some limitations. In particular, it is not designed to describe population structure (Section II B), and it tends to be less accurate for rare variants (Supplementary Sections S4 H and S5 G). However, our framework is sufficiently flexible to accommodate other choices of HMM in the future. Meanwhile, we extend an earlier Monte Carlo algorithm for generating HMM knockoffs^24^ to target the multi-resolution hypotheses defined above (Supplementary Section S1 A). Moreover, we reduce the computational complexity of this operation through exact analytical derivations (Supplementary Section 2). With the UK Biobank data, for which the haplotypes have been phased,^22^ we accelerate the algorithm further by avoiding implicit re-phasing (Supplementary Section S2 B 2).

To powerfully separate interesting signals from spurious associations, we fit a multivariate predictive model of the trait and compute feature importance measures for the original and synthetic genotypes. The contrast between the importance of the genotypes and their knockoffs is used to compute a test statistic in each of the LD blocks defined above, for which a significance threshold is calibrated by the *knockoff filter*.^36^ As a predictive model, we adopt an efficient implementation of sparse linear and logistic regression designed for massive genetic datasets,^38^ although *KnockoffZoom* could easily incorporate other methods.^23^ Thus, *KnockoffZoom* exploits the statistical power of variable selection techniques that would otherwise offer no type-I error guarantees.^23,31^ This methodology is described in more detail in the Online Methods B. A full schematic of our workflow is in Supplementary Figure S1, while software and tutorials are available from https://msesia.github.io/knockoffzoom. The computational cost (Supplementary Table S1) compares favorably to that of alternatives, e.g., BOLT-LMM.^7,8^

Revisiting Figure 1, we note that our discoveries are clearly interpretable because they are distinct by construction and each suggests the presence of an interesting SNP. The interpretations of hypotheses at different resolutions are easily reconciled because our partitions are nested (each block is contained in exactly one larger block from the resolution below), while the null hypothesis for a group of variants is true if and only if all of its subgroups are null.^39^ Most of our findings are confirmed by those at lower resolution, even though this is not explicitly enforced and some “floating” blocks are occasionally reported, as also visible in Figure 1. These are either false positives or true discoveries below the adaptive significance threshold for FDR control at lower resolution. A variation of our procedure can explicitly avoid “floating” blocks by coordinating discoveries at multiple resolutions, although with some power loss (Supplementary Section S1 B).

While our final output at each resolution is a set of distinct discoveries that controls the FDR, it is also possible to quantify the statistical significance of individual findings, as long as these are sufficiently numerous, by estimating a local version of the FDR,^40^ as explained in Supplementary Section S4 K.

Our findings lend themselves well to cross-referencing with gene locations and functional annotations. An example is shown in Figure 5. By contrast, the output of BOLT-LMM in Figure 1 (c) is less informative: many of the clumps reported by the standard PLINK algorithm (Section II D 2) are difficult to interpret because they are wide and overlapping. This point is clearer in simulations where we know the causal variants, as previewed in Figure 2. Here, we see two consequences of stronger signals: our method precisely identifies the causal variants, while the LMM reports wider and increasingly contaminated sets of associated SNPs. The details of our numerical experiments are discussed in Section II D.

**FIG. 2.**
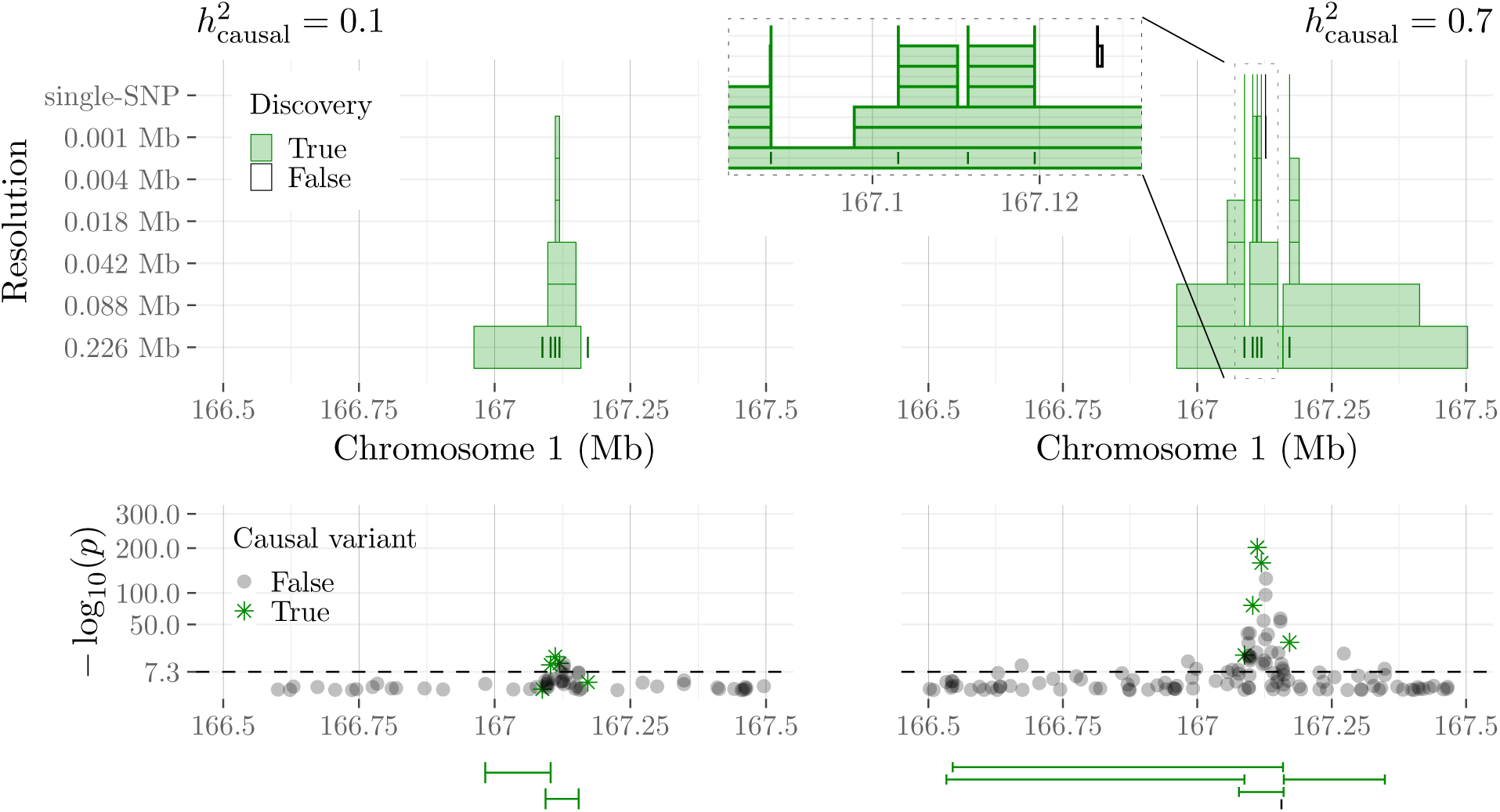
*KnockoffZoom* discoveries for two simulated traits with the same genetic architecture but different heritability 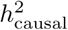, within a genomic window. Other details are as in Figure 1. This region contains 5 causal SNPs whose positions are marked by the short vertical segments at the lowest resolution. The zoomed-in view (right) shows the correct localization of the causal SNPs, as well as a “floating” false positive.

**FIG. 3.**
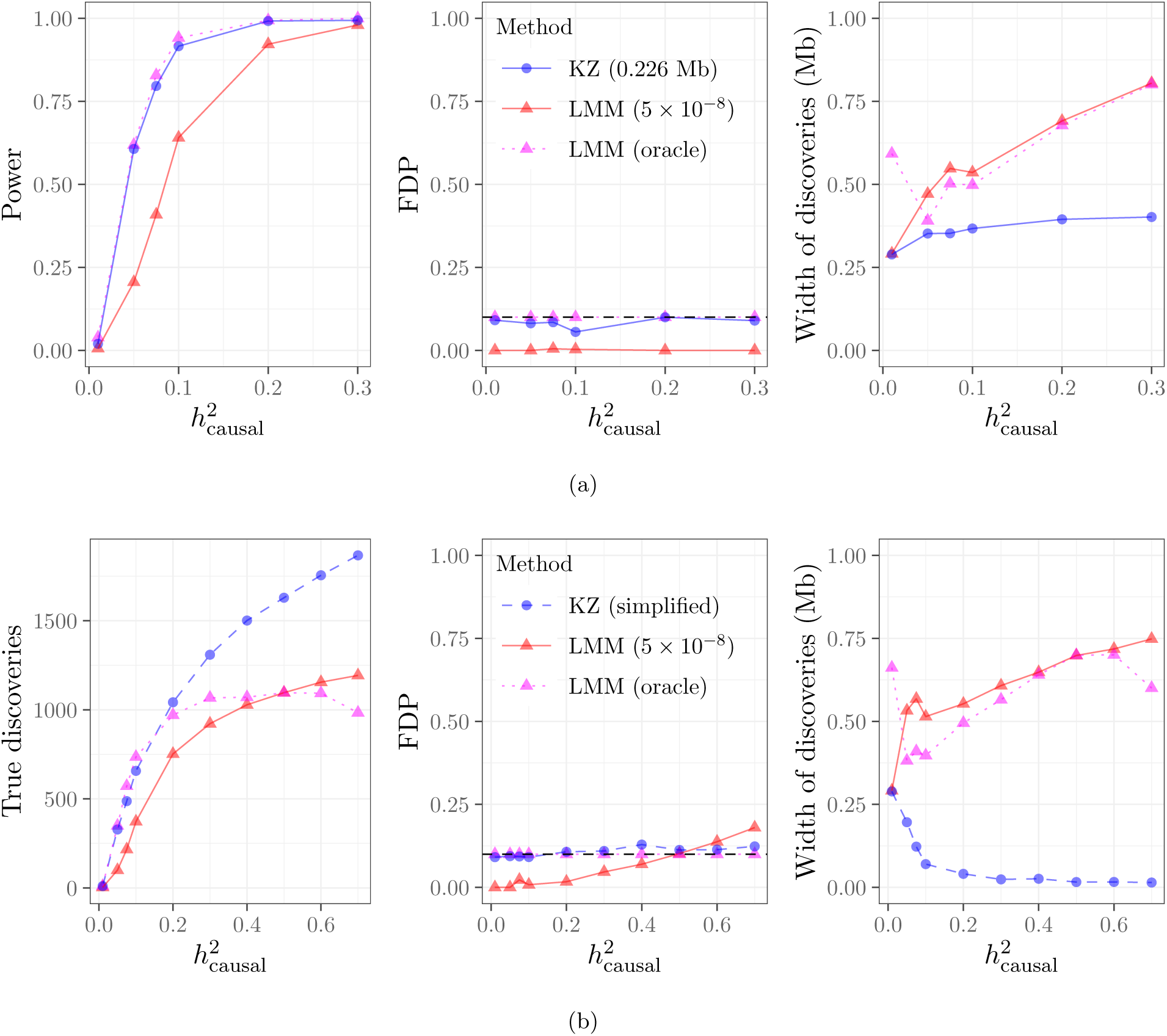
Locus discovery for a simulated trait with *KnockoffZoom* (nominal FDR 0.1) and BOLT-LMM (5 × 10^−8^ and *oracle*). (a): low-resolution *KnockoffZoom* and strongly clumped LMM p-values. (b): multi-resolution *KnockoffZoom* (simplified count) and weakly clumped LMM p-values.

**FIG. 4.**
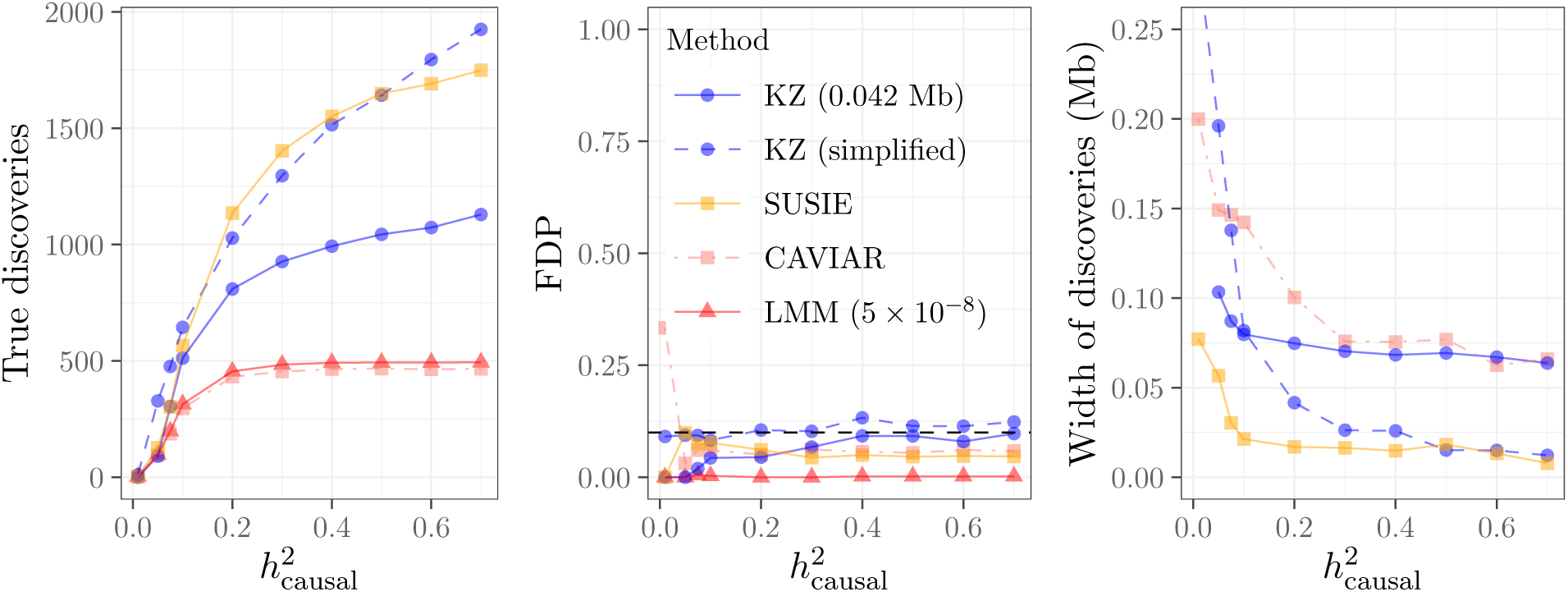
*KnockoffZoom* compared to a two-step fine-mapping procedure consisting of BOLT-LMM followed by CAVIAR or SUSIE, in the same simulations as in Figure 3. Our method, CAVIAR and SUSIE control a similar notion of FDR at the nominal level 0.1.

**FIG. 5.**
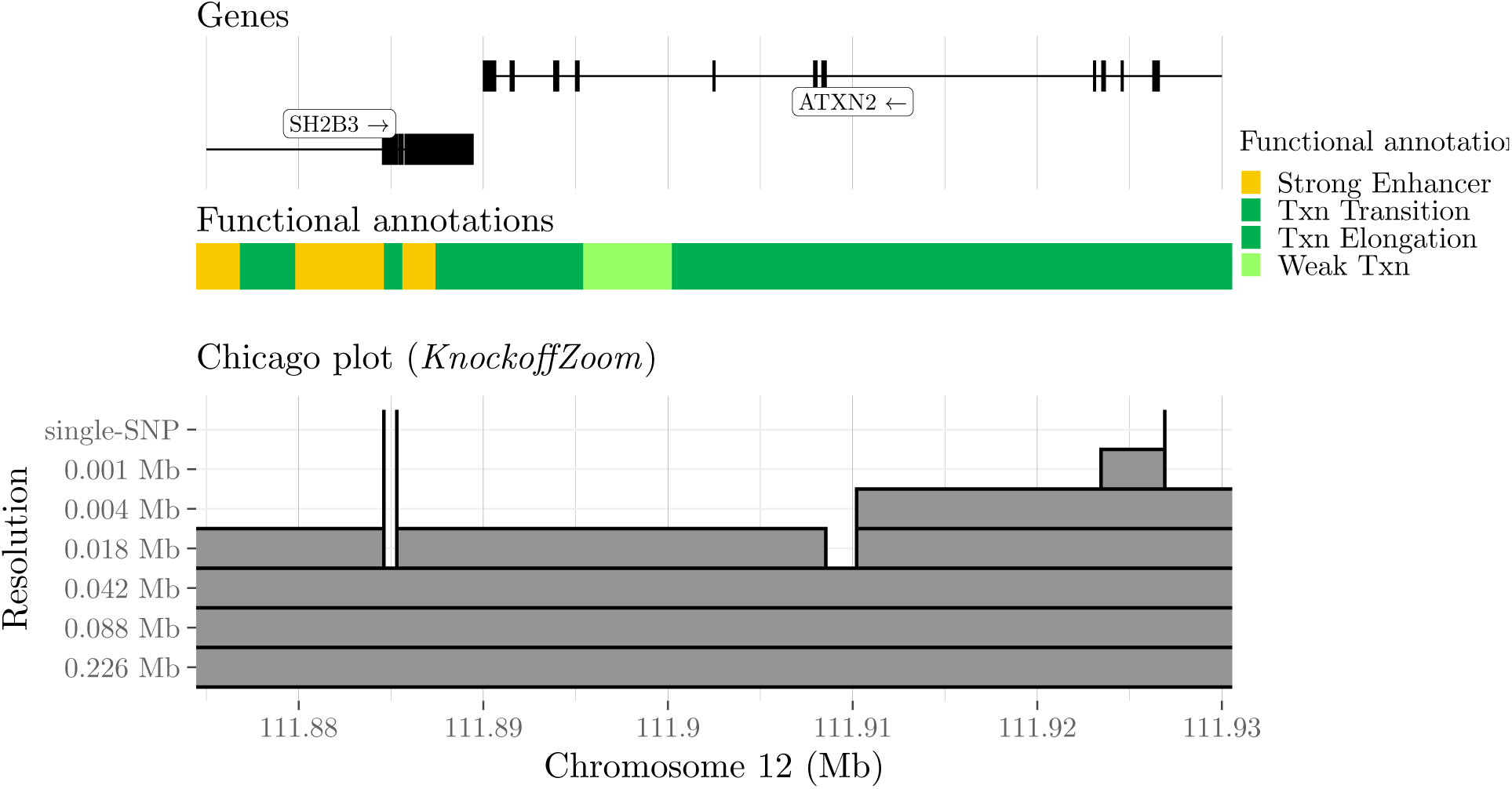
Visualization of some discoveries made with *KnockoffZoom* for *platelet* in the UK Biobank, along with gene positions and functional annotations (in the LocusZoom^45^ style). The three discoveries at single-variant resolution are labeled. Other details as in Figure 1.

### B. Conditional hypotheses and population structure

*KnockoffZoom* discoveries, by accounting for LD, bring us closer to the identification of functional variants; therefore, they are more interesting than simple marginal association. Moreover, conditional hypotheses are less susceptible to the confounding effect of population structure.^29^ As an illustration, consider studying the genetic determinants of blood cholesterol levels, blindly sampling from European and Asian populations, which differ in the distribution of the trait (due to diet or genetic reasons) and in the prevalence of several markers (due to different ancestries). It is well known that marginal associations do not necessarily indicate interesting biological effects, as they may simply reflect a diversity in the allele frequencies across populations (which is correlated with the trait). This is why the traditional analysis must account for population structure via principal component analysis^41^ or linear mixed models.^3–5^ By contrast, multivariate methods, like *KnockoffZoom*, are naturally less susceptible to such confounding because they already account for the information encompassed in all genotypes, which includes the population structure.^4,5,42^

Conditional hypotheses are most robust at the highest resolution, where we condition on all SNPs except one, capturing most of the genetic variation. The susceptibility to confounding may increase as more variants are grouped and removed from the conditioning set, since these may be informative regarding the population structure. However, we always consider fairly small LD blocks, typically containing less than 0.01% of the SNPs, even at the lowest resolution. By comparison, LMMs may not account for an entire chromosome at once, to avoid “proximal contamination”.^7^

Our inferences rely on a model for the distribution of the genotypes, which requires some approximation. The HMM implemented here^20^ is more suitable to describe homogeneous and unrelated individuals because it does not capture long-range dependencies, which is a technical limitation we plan to lift in the future. Meanwhile, we analyze unrelated British individuals, for which we verify in Section II D the robustness of our approximation with simulations involving real genotypes. In the data analysis of Section II E, our results already explicitly account for any available covariates, including the top principal components of the genetic matrix; see Section II E. Moreover, one could account for any known structure (labeled populations) with the current implementation of our procedure by fitting separate HMMs and generating knockoffs independently for each population.

### C. Missing and imputed variants

In this paper, we analyze genotype data from SNP arrays, which only include a fraction of all variants. It is then likely that at least some of the true causal variants are missing. In this sense, our conditional hypotheses are useful proxies for the ultimate goal, as they *localize* important effects precisely, although they may not allow us to *identify* exactly the true causal variants. Only in the simulations below, where we know that the causal variants are not missing, we can verify that *KnockoffZoom* identifies them exactly while controlling the FDR.

In a data analysis, one could impute the missing variants and then analyze them as if they had been measured. However, while meaningful in the case of marginal tests, this would not be useful to study conditional association as defined in the Online Methods A. Imputed variants contain no additional information about any phenotype beyond that already carried by the genotyped SNPs; in fact, they are conditionally independent of any phenotype given the genotyped SNPs. This is because the imputed values are a function of the SNP data and of an LD model estimated from a separate panel of independent individuals. This implies that it is impossible to determine whether the true causal variant is the missing one or among those used to impute it, unless one is willing to make much stricter modeling assumptions whose validity cannot be verified (e.g., homoscedastic Gaussian linear models), as discussed in Supplementary Section S6 A.

### D. Power, resolution and false positive control in simulations

#### 1. Setup

To test and compare *KnockoffZoom* with state-of-the-art methods, we first rely on simulations. These are designed using 591k SNPs from 350k unrelated British individuals in the UK Biobank (Online Methods C). We simulate traits using a linear model with Gaussian errors: 2500 causal variants are placed on chromosomes 1–22, clustered in evenly-spaced 0.1 Mb-wide groups of 5. The effect sizes are heterogeneous, with relative values chosen uniformly at random across clusters so that the ratio between the smallest and the largest is 1/19. The heritability is varied as a control parameter. The causal variables in the model are standardized, so rarer variants have stronger effects. The direction of the effects is randomly fixed within each cluster, using the major allele as reference. This architecture is likely too simple to be realistic,^11^ but it facilitates the comparison with other tools, whose assumptions are reasonable in this setting. By contrast, we are not protecting *KnockoffZoom* against any model misspecification because we use real genotypes and approximate their distribution (our sole assumption is that we can do this accurately) with the same HMM as in the analysis of the UK Biobank phenotypes. Therefore, the only difference between the simulations and the analysis is the conditional distribution of *Y* | *X*, about which our method makes no assumptions. In this sense, the simulations explicitly demonstrate the robustness of *KnockoffZoom* to model misspecification, including the possible presence of some population structure among the unrelated British individuals in the UK Biobank (Section II B), and the lower accuracy of the implemented HMM for rarer variants (Supplementary Sections S4 H and S5 G).

In this setup, the tasks of *locus discovery* and *fine-mapping* are clearly differentiated. The goal of the former is to detect broad genomic regions that contain interesting signals; also, scientists may be interested in counting how many distinct associations have been found. The goal of the latter is to identify the causal variants precisely. Here, there are 500 well-separated regions and 2500 signals to be found. For locus discovery, we compare in Section II D 2 *KnockoffZoom* at low resolution to BOLT-LMM.^8^ For fine-mapping, we apply in Section II D 3 *KnockoffZoom* at 7 levels of resolution (Supplementary Section S4 B), and compare it to CAVIAR^14^ and SUSIE^17^ as discussed below (see Supplementary Section S4 G 1 for more details about these fine-mapping tools). For each method, we report the power, false discovery proportion (FDP) and mean width of the discoveries (the distance in base pairs between the leftmost and rightmost SNPs). Since locus discovery and fine-mapping have different goals, we need to clearly define false positives and count the findings appropriately; our choices are detailed in the next sections. Simulations with explicit coordination of the *KnockoffZoom* discoveries at different resolutions are discussed in Supplementary Section S4 J. Further details regarding the simulation code, relevant third-party software, and tuning parameters can be found in Supplementary Section S4 L.

#### 2. Locus discovery

We apply *KnockoffZoom* at low resolution, with typically 0.226 Mb-wide LD blocks, targeting an FDR of 0.1. For BOLT-LMM,^8^ we use the standard p-value threshold of 5 × 10^−8^, which has been motivated by a Bonferroni correction to control the family-wise error rate (FWER) below 0.05, for the SNP-by-SNP hypotheses of no marginal association.

To assess power, any of the 500 interesting genomic regions is detected if there is at least one finding within 0.1 Mb of a causal SNP. This choice favors BOLT-LMM, which is not designed to precisely localize causal variants and reports wider findings (Figure 3). To evaluate the FDP, we need to count true and false discoveries. This is easy with *KnockoffZoom* because its findings are distinct and count as false positives if causal variants are not included. In comparison, distinct LMM discoveries are more difficult to define because the p-values test marginal hypotheses. A common solution is to group nearby significant loci. Throughout this paper, we clump with the PLINK^12^ greedy algorithm with the same parameters found in earlier work,^8^ as explained in Supplementary Section S4 L. For reference, this is how the clumps in Figure 1 are obtained. Then, we define the FDP as the fraction of clumps whose range does not cover a causal SNP. For locus discovery, we consolidate clumps within 0.1 Mb.

*KnockoffZoom* and BOLT-LMM target different error rates, complicating the comparison. Ideally, we would like to control the FDR of the distinct LMM findings. Unfortunately, this is difficult^35^ (Supplementary Section S4 E). Within simulations, we can address this obstacle by considering a hypothetical *oracle* procedure based on the LMM p-values that is guaranteed to control the FDR. The oracle knows in advance which SNPs are causal and uses this information to identify the most liberal p-value threshold such that the fraction of falsely reported clumps is below 0.1. Clearly, this cannot be implemented in practice. However, the power of this oracle provides an informative upper bound for any future concrete FDR-controlling procedure based on BOLT-LMM p-values.

Figure 3 (a) compares the performances of these methods as a function of the heritability. The results refer to an independent realization of the simulated phenotypes for each heritability value. The average behavior across repeated experiments is discussed in Supplementary Section S4 I. The FDP of *KnockoffZoom* is below the nominal level (the guarantee is that the FDP is controlled on average), while its power is comparable to that of the oracle. Our method consistently reports more precise (narrower) discoveries, while the BOLT-LMM discoveries become wider as the signal strength increases, as anticipated from Figure 2. Moreover, the standard LMM p-value threshold of 5 × 10^−8^ is substantially less powerful.

Before considering alternative fine-mapping methods, we compare *KnockoffZoom* and BOLT-LMM at higher resolutions. Above, the block sizes in *KnockoffZoom* and the aggressive LMM clumping strategy are informed by the known location of the regions containing causal variants, to ensure that each is reported at most once. However, this information is generally unavailable and the scientists must determine which discoveries are important. Therefore, we set out to find as many distinct associations as possible, applying *KnockoffZoom* at multiple resolutions (Supplementary Section S4 B), and interpreting the LMM discoveries assembled by PLINK as distinct, without additional consolidation. An example of the outcomes is visualized in Figure 2. In this context, we follow a stricter definition of type-I errors: a finding is a true positive if and only if it reports a set of SNPs that includes a causal variant. We measure power as the number of true discoveries. Instead of showing the results of *KnockoffZoom* at each resolution (Supplementary Figure S6), we count only the most specific findings whenever the same locus is detected with different levels of granularity, and discard finer discoveries that are unsupported at lower resolutions (simplified count). This operation is unnecessary to interpret our results and is not sustained by theoretical guarantees,^43^ although it is quite natural and informative.

Figure 3 (b) summarizes these results as a function of the heritability. *KnockoffZoom* reports increasingly precise discoveries as the signals grow stronger. By contrast, the ability of BOLT-LMM to resolve distinct signals worsens. As stronger signals make more non-causal variants marginally significant through LD, the interpretation of the marginal p-values becomes more opaque and counting different clumps as distinct discoveries clearly leads to an excess of false positives. For this reason, we see that the oracle procedure that knows the ground truth and always controls the FDP must become more conservative and report fewer discoveries when the signals are strong. Overall, these results caution one against placing too much confidence in the estimated number of distinct findings obtained with BOLT-LMM via clumping.^8^

#### 3. Fine-mapping

In this section, we start with BOLT-LMM (5 × 10^−8^ significance) for locus discovery and then separately fine-map each reported region with either CAVIAR^14^ or SUSIE,^17^ since these methods are not designed to operate genome-wide. We aggressively clump the LMM findings to ensure that they are distinct, as in Figure 3 (a). We also provide unreported nearby SNPs as input to SUSIE, to attenuate the selection bias (Supplementary Figure S7). Within each region, CAVIAR and SUSIE report sets of SNPs that are likely to contain causal variants. We tune their parameters to obtain a genome-wide FDR comparable to our target (Supplementary Section S4 G 1).

The results are shown in Figure 4, defining the power and FDP as in Figure 3 (b). The output of *KnockoffZoom* is presented in two ways: at a fixed resolution (e.g., 0.042 Mb; see Supplementary Table S3) and summarizing the multi-resolution results by counting only the highest-resolution discoveries in each locus, as in Figure 3 (b). Again, this simplified count is useful to summarize the performance of our method within these simulations, even though in theory we can only guarantee that the FDR is controlled at each level of resolution separately. All methods considered here appear to control the FDR and detect the 500 interesting regions as soon as the heritability is sufficiently large, as seen earlier in Figure 3. Moreover, they report precise discoveries, each including only a few SNPs. CAVIAR cannot make more than 500 discoveries here, as it is intrinsically unable to distinguish between multiple causal variants in the same locus.^17^ SUSIE and *KnockoffZoom* successfully identify more distinct signals, with comparable performance in this setting. A more detailed analysis highlighting their differences is available in Supplementary Section S4 G 2.

Finally, note that in the experiments of Figure 4 we have applied *KnockoffZoom* at each resolution to test pre-defined hypotheses defined over contiguous groups of SNPs, while CAVIAR and SUSIE have some additional flexibility in the sense they can report sets of non-contiguous SNPs. Therefore, *KnockoffZoom* may group together nearby SNPs that are not as highly correlated with each other compared to those grouped together by the other methods, especially when the signal is too weak to obtain very high-resolution discoveries, as shown in Supplementary Figure S9. As mentioned earlier in Section II A, contiguity is not required by *KnockoffZoom* in principle, but we find it to be particularly meaningful for genetic mapping in studies where not all variants are genotyped (Section II C), as discussed further in Supplementary Section S4 G 3.

### E. Analysis of different traits in the UK Biobank data

#### 1. Findings

We apply *KnockoffZoom* to 4 continuous traits and 5 diseases in the UK Biobank (Supplementary Table S5), using the same SNPs from 350k unrelated individuals of British ancestry (Online Methods C) and the same knockoffs of Section II D. Our discoveries are compared to the BOLT-LMM findings in Table I. We apply our method accounting for the top 5 principal components^41^ and a few other covariates in the predictive model (Online Methods D). The results do not change much if we ignore the principal components (Supplementary Table S6), confirming the intrinsic robustness to population structure. *KnockoffZoom* misses very few of the BOLT-LMM discoveries (see Online Methods B for more details about the case of *glaucoma*) and reports many additional findings (Supplementary Table S7), even when the latter is applied to a larger sample^8^ (Supplementary Table S8). The interpretation of the LMM findings is unclear because many clumps computed by PLINK are overlapping, as shown in Figure 1. As we consolidate them, they become distinct but less numerous, inconsistently with the results reported by others using the same data.^8^

**TABLE I.**
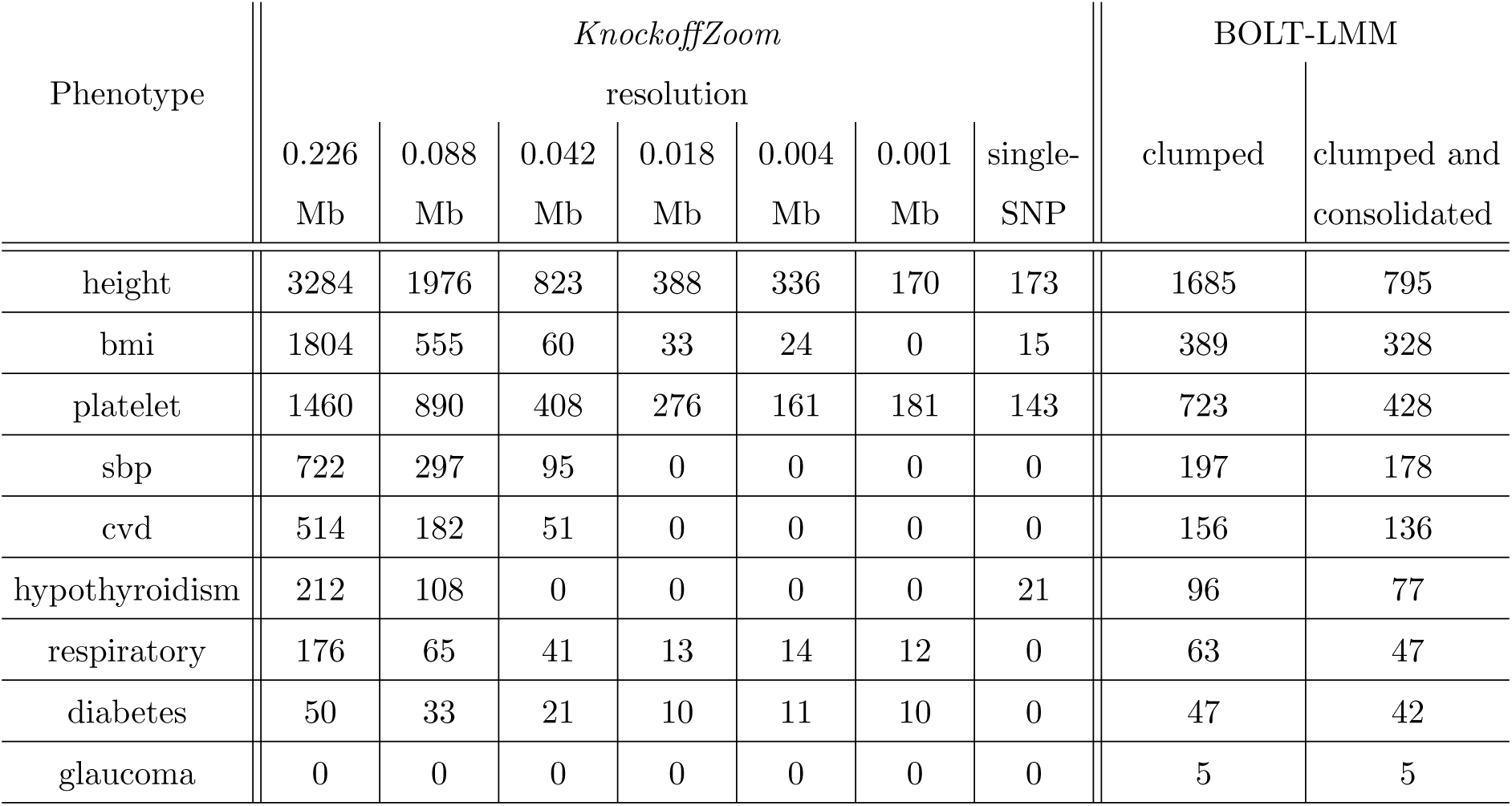
Numbers of distinct findings made by *KnockoffZoom* at different resolutions (FDR 0.1), on some phenotypes in the UK Biobank, compared to BOLT-LMM (p-values ≤ 5 × 10^−8^). Data from 350k unrelated British individuals. The LMM discoveries are counted with two clumping heuristics, as in Figure 3.

| Phenotype | *KnockoffZoom* |  |  |  |  |  |  | BOLT-LMM |  |
| --- | --- | --- | --- | --- | --- | --- | --- | --- | --- |
|  | resolution |  |  |  |  |  |  | clumped | clumped and consolidated |
|  | 0.226 Mb | 0.088 Mb | 0.042 Mb | 0.018 Mb | 0.004 Mb | 0.001 Mb | single-SNP |  |  |
| height | 3284 | 1976 | 823 | 388 | 336 | 170 | 173 | 1685 | 795 |
| bmi | 1804 | 555 | 60 | 33 | 24 | 0 | 15 | 389 | 328 |
| platelet | 1460 | 890 | 408 | 276 | 161 | 181 | 143 | 723 | 428 |
| sbp | 722 | 297 | 95 | 0 | 0 | 0 | 0 | 197 | 178 |
| cvd | 514 | 182 | 51 | 0 | 0 | 0 | 0 | 156 | 136 |
| hypothyroidism | 212 | 108 | 0 | 0 | 0 | 0 | 21 | 96 | 77 |
| respiratory | 176 | 65 | 41 | 13 | 14 | 12 | 0 | 63 | 47 |
| diabetes | 50 | 33 | 21 | 10 | 11 | 10 | 0 | 47 | 42 |
| glaucoma | 0 | 0 | 0 | 0 | 0 | 0 | 0 | 5 | 5 |

As the resolution increases, we typically report fewer findings, which is unsurprising since high-resolution conditional hypotheses are more challenging, although there a few exceptions. Some exceptions can be explained by the presence of multiple distinct signals within the same locus (e.g., as in Figure 2), while others may be due to random variability in the analyses at different resolutions, which involve different knockoffs. The results obtained by explicitly coordinating discoveries at different resolutions using the variation of *KnockoffZoom* mentioned in Section II A are reported in Supplementary Table S9. In general, our findings can be compared to the known gene positions and the available functional annotations (based on ChromHMM,^44^ GM12878 cell line) to shed more light onto the causal variants. We provide an online tool to explore these results interactively (https://msesia.github.io/knockoffzoom/ukbiobank); for examples, see Figure 5 and Supplementary Figure S15.

#### 2. Reproducibility

To investigate the reproducibility of the low-resolution findings obtained with different methods, we study *height* and *platelet* on a subset of 30k unrelated British individuals and verify that the discoveries are consistent with those previously reported for BOLT-LMM applied to all 459k European subjects.^8^ For this purpose, a discovery is replicated if it falls within 0.1 Mb of a SNP with p-value below 5 × 10^−9^ on the larger dataset. We do not consider the other phenotypes because, with this sample size, both methods make fewer than 10 discoveries for each of them. The LMM findings are clumped by PLINK without consolidation, although this makes little difference because extensive overlap occurs only with larger samples. To illustrate the difficulty of controlling the FDR with the LMM (Supplementary Section 4), we try to naïvely apply a pre-clumping Benjamini-Hochberg (BH) correction^18^ to the LMM p-values.

The results are summarized in Table II. The FDP of *KnockoffZoom* is below the target FDR. All genome-wide significant BOLT-LMM discoveries in the smaller dataset (5 × 10^−8^) are replicated, at the cost of lower power. The BH procedure with the LMM p-values does not control the FDR. The relative power of these alternative methods is discussed in Supplementary Section S5 E.

**TABLE II.** Reproducibility of the low-resolution discoveries made with *KnockoffZoom* (0.226 Mb) and BOLT-LMM on *height* and *platelet*, using 30k individuals in the UK Biobank.

| Phenotype | Method | Discoveries |  |  |
| --- | --- | --- | --- | --- |
|  |  | # | Not replicated | Size (Mb) |
| height | *KnockoffZoom* (FDR 0.1) | 121 | 8 (6.6%) | 0.308 |
| | LMM ($5 \times 10^{-8}$) | 54 | 0 (0.0%) | 0.965 |
|  | LMM-BH (FDR 0.1) | 714 | 203 (28.4%) | 0.379 |
| platelet count | *KnockoffZoom* (FDR 0.1) | 81 | 5 (6.2%) | 0.319 |
| | LMM ($5 \times 10^{-8}$) | 47 | 0 (0.0%) | 0.674 |
|  | LMM-BH (FDR 0.1) | 272 | 92 (33.8%) | 0.433 |

As a preliminary verification of the biological validity of our findings, we use GREAT^46^ to conduct a gene ontology enrichment analysis of the regions reported by *KnockoffZoom* for *platelet* at each resolution. The enrichment among 5 relevant terms is highly significant (Supplementary Table S12) and tends to strengthen at higher resolutions, which suggests increasingly precise localization of causal variants.

Finally, we cross-reference with the existing literature the discoveries reported by *KnockoffZoom* at the highest resolution for *platelet*. Note that three of these findings are shown in Figure 5: *rs3184504* (missense, *SH2B3* gene), *rs72650673* (missense, *SH2B3* gene) and *rs1029388* (intron, *ATXN2* gene). Many of the other discoveries are in coding regions and may plausibly localize a direct functional effect on the trait (Supplementary Table S13). Some have been previously observed to be associated with *platelet* (Supplementary Table S14), while others are likely to indicate new findings. In particular, six of our discoveries are missense variants that had not been previously reported to be associated with *platelet* (Supplementary Table S15).

## III. DISCUSSION

The goal of genetic mapping is to localize, as precisely as possible, variants that influence a trait. Geneticists have widely sought this goal with a two-step strategy, partly due to computational limitations, in an attempt to control type-I errors while achieving high power. First, all variants are probed in order to identify promising regions, without accounting for LD. To reduce false positives, a Bonferroni correction is applied to the marginal p-values. Then, the roles of different variants are explored with multivariate models, separately within each associated region. This strategy is suboptimal. Indeed, if the phenotypes are influenced by hundreds or thousands of genetic variants, the notion of FWER in the first step is too stringent and inhibits power unnecessarily. This error rate is a legacy of the earlier studies of Mendelian diseases and it has been retained for the mapping of complex traits mostly due to methodological difficulties, rather than a true need to avoid any false positives. In fact, we have shown that the traditional two-step paradigm of locus discovery followed by fine-mapping already effectively tries to control an error rate comparable to the FDR that we directly target. Besides, the type-I error guarantee in the first step is only valid for the SNP-by-SNP marginal hypotheses of no association. These are of little interest to practitioners, who typically interpret the findings as if they contained causal variants. By contrast, *KnockoffZoom* unifies locus discovery and fine-mapping into a coherent statistical framework, so that the findings are immediately interpretable and equipped with solid guarantees.

We are not the first to propose a multi-marker approach to genetic mapping,^31,35,42,47–49^ nor to consider testing the importance of groups of SNPs at different resolutions,^50^ although knockoffs finally allow us to provide solid type-I error guarantees based on relatively realistic assumptions.

For ease of comparison, we have simulated phenotypes using a linear model with Gaussian errors, which satisfies many of the assumptions in the standard tests. Unfortunately, there is very little information on how real traits depend on the genetic variants; after all, the goal of a GWAS is to discover this. Therefore, relying on these assumptions can be misleading. In contrast, *KnockoffZoom* only relies on knowledge of the genotype distribution, which we can estimate accurately due to the availability of genotypes from millions of individuals. Indeed, geneticists have developed phenomenological HMMs of LD that work well in practice for phasing and imputation.

It may be useful to highlight with an example that our framework is not tied to any model for the dependence of phenotypes on genotypes or to a specific data analysis tool. We simulate an imbalanced case-control study that it is well-known to cause BOLT-LMM to fail,^51^ while our method applies seamlessly. The trait is generated from a liability threshold model (probit), obtaining 525 cases and 349,594 controls. We apply *KnockoffZoom* using sparse logistic regression and report the low-resolution results in Figure 6 as a function of the heritability of the latent Gaussian variable.

**FIG. 6.**
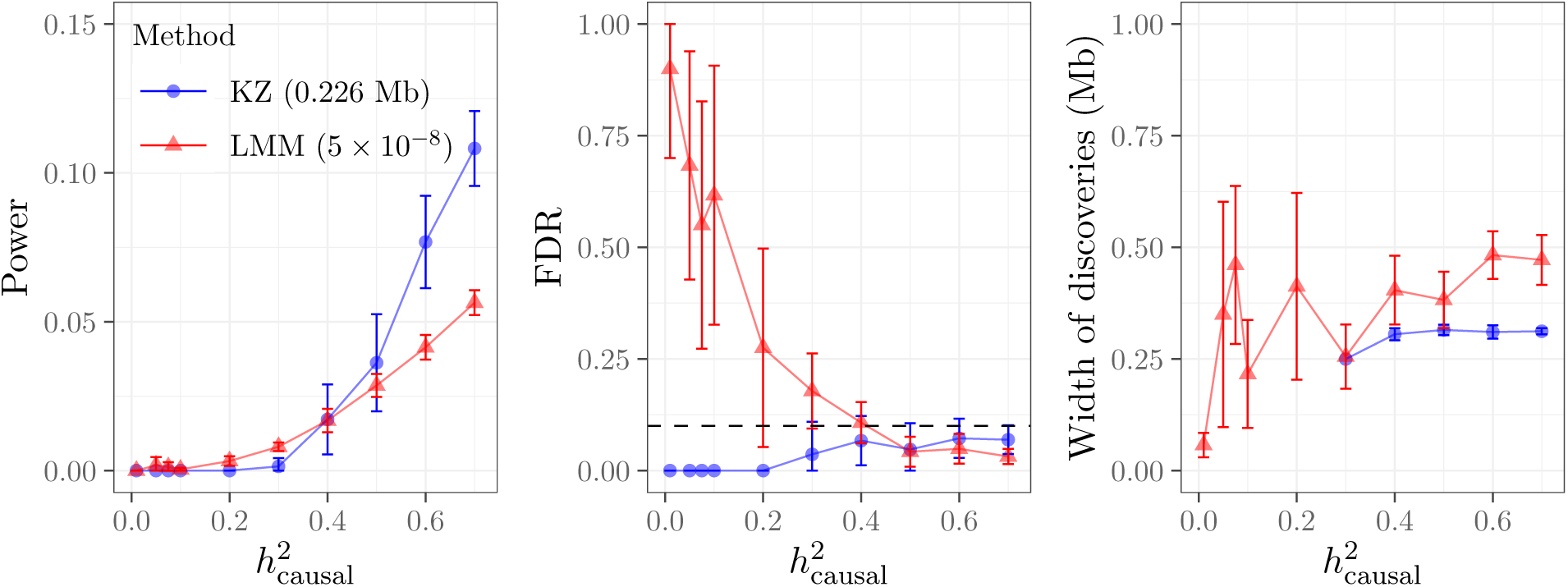
Locus discovery with *KnockoffZoom* and BOLT-LMM for a simulated case-control study. There are 500 evenly spaced causal variants. The error bars indicate 95% confidence intervals for the mean power and FDR, as defined in Figure 3 (a), averaging 10 independent replications of the trait given the same genotypes.

Like many other statistical methods (e.g., data splitting, cross-validation, permutations, Monte Carlo inference), *KnockoffZoom* is randomized, in the sense that its output partially depends on random variables that are not part of the data, in this case the knockoffs. Even though we have not done it in this paper, it is possible to re-sample the knockoffs multiple times and obtain different sets of discoveries.^23,24^ Each set of discoveries is conditionally independent given the data and controls the FDR. However, it is not yet clear how to best combine multiple sets of discoveries while preserving control of the FDR in theory. We believe that this should be the subject of further research but it is technically challenging. Meanwhile, we have applied *KnockoffZoom* for the analysis of the UK Biobank phenotypes using new conditionally independent knockoffs in Supplementary Section S5 J and verified that the discoveries are relatively stable upon resampling, especially when their number is large. It is also worth mentioning that the stabilities of our discoveries can be predicted well from their individual statistical significance, which we can estimate with a local version of the FDR (Supplementary Sections S5 I and S5 J).

Our software (https://msesia.github.io/knockoffzoom) has a modular structure that accommodates many options, reflecting the flexibility of this approach. Users may experiment with different importance measures in addition to sparse linear and logistic regression. For example, one can incorporate prior information, such as summary statistics from other studies. Similarly, there are many ways of defining the LD blocks: leveraging genomic annotations is a promising direction.

We are currently working on more refined knockoff constructions, focusing on new algorithms for heterogeneous populations and rarer variants that rely on an HMM-based approach similar to that of SHAPEIT.^22^ Similarly, we are developing new algorithms for constructing knockoffs for closely related families. In the future, we also plan to leverage these advances to analyze data sets with larger number of variants, possibly from whole-genome sequencing. This will involve additional computational challenges, but it is feasible in principle since our algorithms are parallelizable and their cost scales linearly in the number of variants.

Finally, we highlight that our approach to genetic mapping may naturally lead to the definition of more principled polygenic risk scores, since we already combine multi-marker predictive modeling with interpretable variable selection. Partly with this goal, multi-marker methods for the analysis of GWAS data have been suggested long before our contribution.^31,42,47–49,52,53^ However, knockoffs finally allow us to obtain reproducible findings with provable type-I error guarantees.

## IV. AUTHOR CONTRIBUTIONS

M. S. developed the methodology, designed the experiments, and performed the data analysis. E. K. contributed to some aspects of the analysis (quality control, phenotype extraction and SNP annotations), and the development of the visualization tool. S. B. contributed to the data acquisition. E. C. and C. S. supervised this project. All authors contributed to writing this manuscript.

## V. CODE AVAILABILITY

*KnockoffZoom* is available from https://msesia.github.io/knockoffzoom. The code for these simulations and analysis is available from https://github.com/msesia/ukbiobank_knockoffs.

## VI. DATA AVAILABILITY

Data from the UK Biobank Resource (application 27837); see https://www.ukbiobank.ac.uk/.

## VII. ACKNOWLEDGEMENTS

M. S., E. C. and C. S. are supported by NSF grant DMS 1712800. E. C. and C. S. are also supported by a Math+X grant (Simons Foundation). E. K. is supported by the Hertz Foundation. S. B. is supported by a Ric Weiland fellowship. Finally, we thank Sahitya Mantravadi, Ananthakrishnan Ganesan, Robert Tibshirani and Trevor Hastie.

## ONLINE METHODS

### A. Formally defining the objective of *KnockoffZoom*

We observe a phenotype *Y* ∈ ℝ and genotypes *X* = (*X*_1_, …, *X*_*p*_) ∈ {0, 1, 2}^*p*^ for each individual. We assume that the pairs 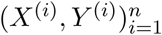 corresponding to *n* subjects are independently sampled from some distribution *P*_*XY*_. The goal is to infer how *P*_*Y*|*X*_ depends on *X*, testing the conditional hypotheses defined below, without assuming anything else about this likelihood, or restricting the sizes of *n* and *p*. Later, we will describe how we can achieve this by leveraging prior knowledge of the genotype distribution *P*_*X*_. Now, we formally define the hypotheses.

Let 𝒢 = (*G*_1_, …, *G*_*L*_) be a partition of {1, …, *p*} into *L* blocks, for some *L* ≤ *p*. For any *g* ≤ *L*, we say that the *g*-th group of variables 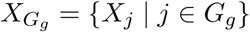 is null if *Y* is independent of 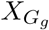 given 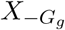 (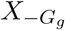 contains all variables except those in *G*_*g*_). We denote by ℋ_0_ ⊆ {1, …, *L*} the subset of null hypotheses that are true. Conversely, groups containing causal variants do not belong to ℋ_0_. For example, if *P*_*Y*|*X*_ is a linear model, ℋ_0_ collects the groups in 𝒢 whose true coefficients are all zero (if there are no perfectly correlated variables in different groups^39^). In general, we want to select a subset *Ŝ* ⊆ {1, …, *L*} as large as possible and such that FDR = 𝔼|*Ŝ* ∩ ℋ_0_|*/* max(1,|*Ŝ*|) ≤ *q*, for a fixed partition 𝒢 of the variants. These conditional hypotheses generalize those defined earlier in the statistical literature,^23,24^ which only considered the variables one-by-one. Our hypotheses are better suited for the analysis of GWAS data because they allow us to deal with LD without pruning the variants (Supplementary Section S1 A). As a comparison, the null statement of the typical hypothesis in a GWAS is that *Y* is marginally independent of *X*_*j*_, for a given SNP *j*.

### B. The knockoffs methodology

*Knockoffs*^23^ solve the problem defined above if *P*_*X*_ is known and tractable, as explained below. The idea is to augment the data with synthetic variables, one for each genetic variant. We know that the knockoffs are null because we create them without looking at *Y*. Moreover we construct them so that they behave similarly to the SNPs in null groups and can serve as negative controls. The original work considered explicitly only the case of a trivial partition 𝒢 into *p* singletons,^23^ but we extend it for our problem by leveraging some previous work along this direction.^37,39^ Formally, we say that 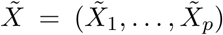 is a group-knockoff of *X* for a partition 𝒢 of {1, …, *p*} if two conditions are satisfied: (1) *Y* is independent of 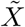 given *X*; (2) the joint distribution of 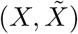 is unchanged when {*X*_*j*_ : *j* ∈ *G*} is swapped with 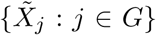, for any group *G* ∈ 𝒢. The second condition is generally difficult to satisfy (unless 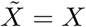, which yields no power), depending on the form of *P*_*X*_. ^23^ In Supplementary Section S2, we develop algorithms to generate powerful group-knockoffs when *P*_*X*_ is an HMM, the parameters of which are fitted on the available data using fastPHASE;^20^ see Supplementary Sections S4 A–S4 C for more details about the model estimation and its goodness-of-fit. Here, we take 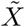 as given and discuss how to test the conditional hypotheses. For the *g*-th group in 𝒢, we compute feature importance measures *T*_*g*_ and 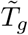 for {*X*_*j*_ : *j* ∈ *G*_*g*_} and 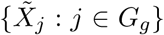, respectively. Concretely, we fit a sparse linear (or logistic) regression model^38^ for *Y* given 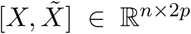, standardizing *X* and 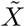; then we define 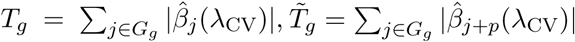. Above, 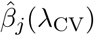 and 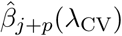 indicate the estimated coefficients for *X*_*j*_ and 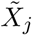, respectively, with regularization parameter *λ*_CV_ tuned by cross-validation. These statistics are designed to detect sparse signals in a generalized linear model—a popular approximation of the distribution of *Y* in a GWAS.^31^ Our power may be affected if this model is misspecified but our inferences remain valid. A variety of other tools could be used to compute more flexible or powerful statistics, perhaps incorporating prior knowledge.^23^ Finally, we combine the importance measures into test statistics *W*_*g*_ through an anti-symmetric function, e.g., 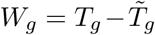, and report groups of SNPs with sufficiently large statistics.^23^ The appropriate threshold for FDR control is calculated by the knockoff filter.^36^ Further details about the test statistics are in Supplementary Section S3.

As currently implemented, our procedure has no power at the nominal FDR level *q* if there are fewer than 1*/q* findings to be made. Usually, this is not a problem for the analysis of complex traits, where many loci are significant. However, this may explain why, at the FDR level *q* = 0.1, we report none of the 5 discoveries obtained by BOLT-LMM for *glaucoma* in Table I. Alternatively, our method may detect these by slightly relaxing the knockoff filter,^36^ at the cost of losing the provable FDR guarantee.

### C. Quality control and data pre-processing for the UK Biobank

We consider 430,287 genotyped and phased subjects with British ancestry. According to the UK Biobank, 147,718 of these have at least one close relative in the dataset; we keep one from each of the 60,169 familial groups, chosen to minimize the missing phenotypes. This yields 350,119 unrelated subjects. We only analyze biallelic SNPs with minor allele frequency above 0.1% and in Hardy-Weinberg equilibrium (10^−6^), among our 350,119 individuals. The final SNPs count is 591,513. A few subjects withdrew consent and we removed their observations from the analysis.

### D. Including additional covariates

We control for the sex, age and squared age of the subjects to increase power (squared age is not used for *height*, as in earlier work^8^). We leverage these covariates by including them in the predictive model for the *KnockoffZoom* test statistics. We fit a sparse regression model on the augmented matrix of explanatory variables 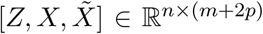, where 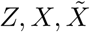 contain the *m* covariates, the genotypes and their knockoff copies, respectively. The coefficients for *Z* are not regularized and we ignore them in the final computation of the test statistics.

## Supplementary Material

### S1. METHODS

#### A. Knockoffs for composite conditional hypotheses

A typical GWAS involves so many variants in LD that the conditional importance of a single SNP given all others may be very difficult to resolve. We simplify this problem by clustering the loci into LD blocks and testing group-wise conditional hypotheses (Online Methods A). In particular, we test whether all variants in any given block are independent of the trait conditional on the rest of the genome.

Consider the following explanatory example. A trait *Y* is described by a linear model (for simplicity) with 4 explanatory variables: 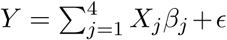 and Gaussian noise *ϵ*. Given independent observations of *X* and *Y*, drawn with *β*_1_ = 1 and *β*_*j*_ = 0 for all *j* ≠ 1, we want to discover which variables influence *Y* (i.e., *X*_1_). To make this example interesting, we imagine an extreme form of LD: *X*_1_ = *X*_2_ and *X*_3_ = *X*_4_, while *X*_1_ and *X*_3_ are independent. This makes it impossible to retrospectively understand whether it is *X*_1_ or *X*_2_ that affects *Y*. However, we can still hope to conclude that either *X*_1_ or *X*_2_ are important. In fact, introductory statistics classes teach us how to do this with an *F* -test, as opposed to a *t*-test, which would have no power in this case.

To solve the above problem within our framework (Online Methods B), we begin by generating powerful knockoffs specifically designed to test group-wise conditional hypotheses. The exact structure of the LD blocks must be taken into account when we create knockoffs. For example, if we naïvely generate knockoffs 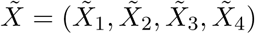 that are pairwise exchangeable with *X* one-by-one, we must set 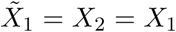 to preserve the equality in distribution between 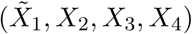 and 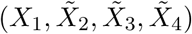 when 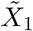 is swapped with *X*_1_. Furthermore, by the same argument we conclude that we need 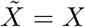. However, these knockoffs are powerless as negative controls (although they are exchangeable). Fortunately, we can generate powerful group-knockoffs (Online Methods B) under milder exchangeability constraints that can still be used to test grouped hypotheses. In the above toy example, we consider the partition 𝒢 = ({1, 2}, {3, 4}), and require that 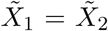 and 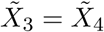, while allowing 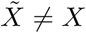.

##### Definition 1

(Group-knockoffs). *Consider random variables Z* = (*Z*_1_, …, *Z*_*p*_), *and a partition* 𝒢 = (*G*_1_, …, *G*_*L*_) *of* {1, …, *p*}. *Then* 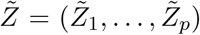 *is said to be a* group-knockoff *for Z with respect to* 𝒢 *if for each group G* ∈ 𝒢, *we have:*

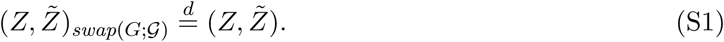

*Above*,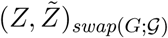 *means that the jth coordinate is swapped with the* (*j*+*p*)*th coordinate* ∀*j* ∈ *G.*

#### B. Coordinating discoveries across resolutions

In this section, we show how to combine the tests statistics computed by our method at different resolutions in order to coordinate the discoveries, so that no “floating” blocks are reported, while rigorously controlling the FDR.

Let *r* = 1, …, *R* index the resolutions we consider (Online Methods A), ordered from the lowest to the highest. At each resolution *r*, the *p* variants are partitioned into *L*^*r*^ groups: 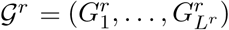. By construction, the partitions are nested, so that for each resolution *r >* 1 and each group *g* ∈ {1, …, *L*^*r*^} there is a unique parent group Pa(*g, r*) ∈ {1, …, *L*^*r*−1^} such that 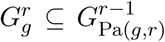. Suppose that we have computed the test statistics 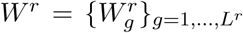 at each resolution (Online Methods B). To avoid “floating” discoveries, we can only select groups whose parent has been discovered at the resolution below. We can enforce this consistency property while preserving an FDR guarantee at each resolution by slightly modifying the final filtering step that computes the significance thresholds (Online Methods B). Instead of applying the knockoff filter separately at each resolution, we proceed with Algorithm 1 sequentially, from lower to higher resolutions.

##### Algorithm 1 Consistent-layers knockoff filter

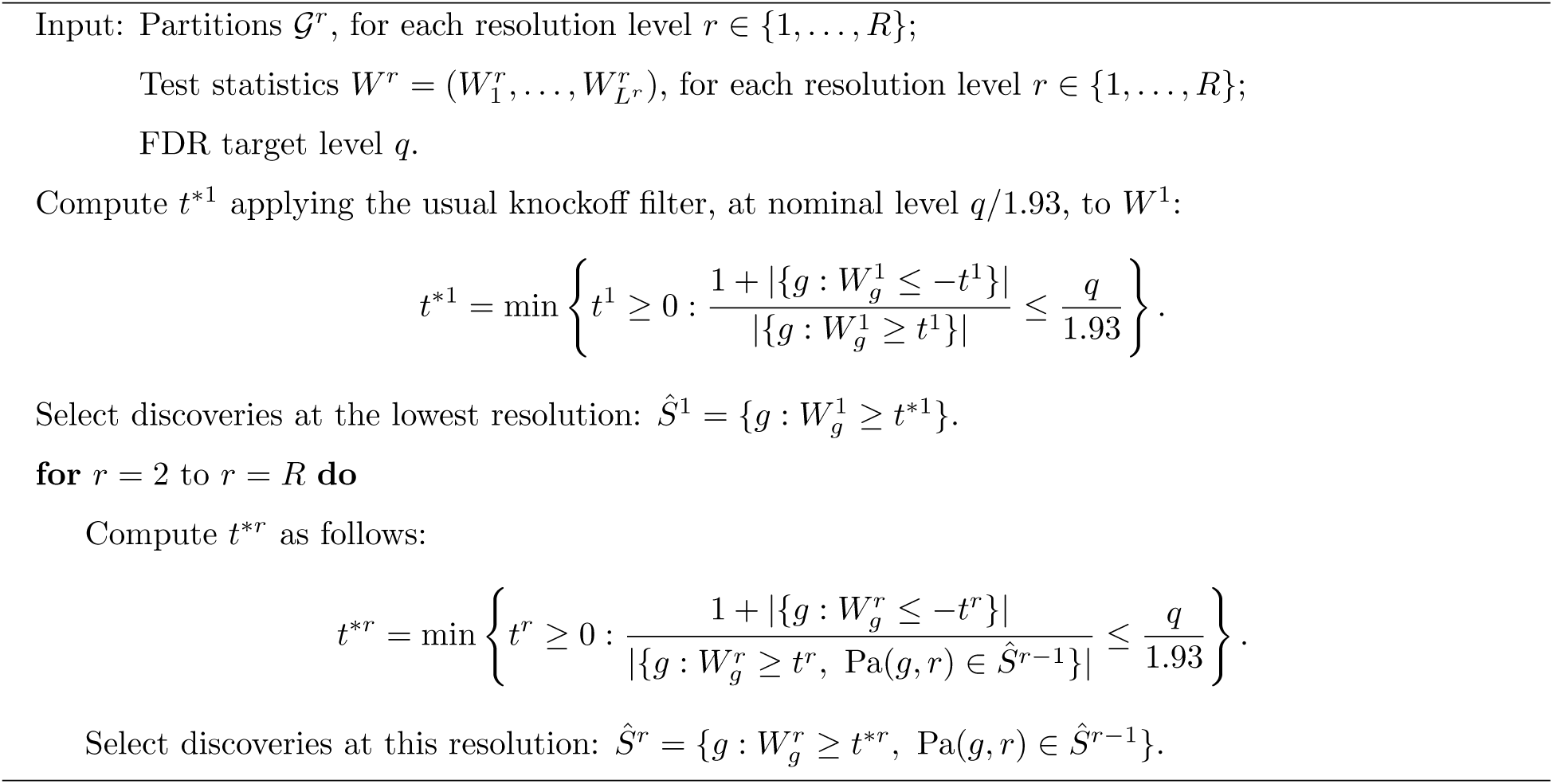

We prove in Proposition 1 that the FDR is controlled at each resolution by Algorithm 1, which is closely inspired by previous work.^1^ The correction of the FDR level by the factor 1.93 is required in the proof for technical reasons, although we have observed in numerical simulations that this may be practically unnecessary; see Section S4 J for empirical evidence. Therefore, to avoid an unjustified power loss while retaining provable guarantees, we have not reported the results obtained with Algorithm 1 in the main paper. However, we have applied it on real data and reported the results in Section S5 D. Based on our experience, should “floating” discoveries be particularly undesirable, we believe that it may be safe to apply Algorithm 1 even without the 1.93 factor.

##### Proposition 1.

*Denote by* FDR^*r*^ *the FDR at resolution r for the discoveries obtained with Algorithm 1. If the data sampling assumptions (Online Methods A) are valid and the knockoff exchangeability holds (Online Methods B), then* FDR^*r*^ ≤ 1.93 · *q*, ∀*r* ∈ {1, …, *R*}.

*Proof.* It was shown in earlier work^1^ (proof of Theorem 1 therein) that

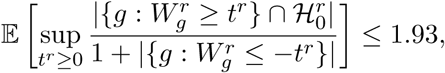

where 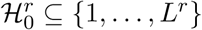 is the set of null groups at resolution *r*. Therefore,

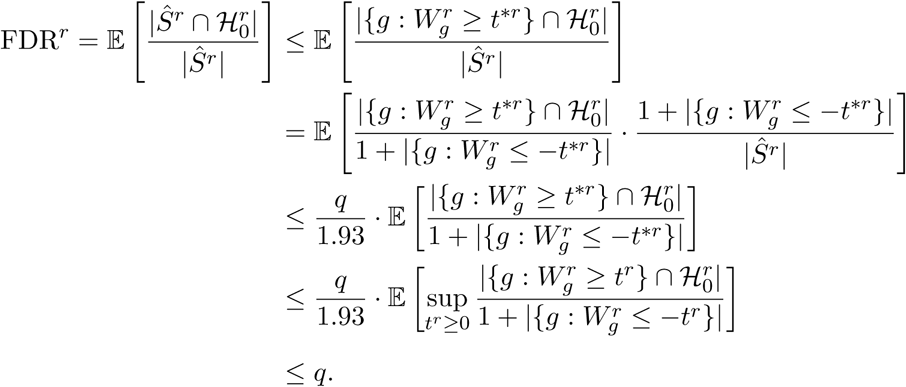

□

### S2. ALGORITHMS FOR KNOCKOFF GENERATION

The construction of knockoffs for groups of genetic variants is the heart of the methodology in this paper and requires a significant extension of the existing algorithms.^2^ Our contribution has two main components: first, we construct group-knockoffs for Markov chains and HMMs; second, we specialize these ideas to obtain fast algorithms for the models of interest in a GWAS. The details of these contributions are in this section, while the associated technical proofs are in Section S7.

#### A. General algorithms for group-knockoffs

##### 1. Markov chains

We begin with group-knockoffs for Markov chains.^2^ We say that *Z* = (*Z*_1_, …, *Z*_*p*_), with each variable taking values in {1, …, *K*} for some *K* ∈ ℕ, is a discrete Markov chain if its joint probability mass function can be written as:

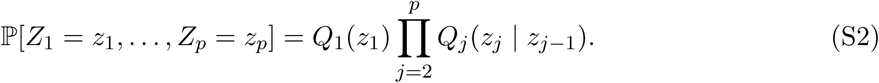

Above, *Q*_1_(*z*_1_) = ℙ[*Z*_1_ = *z*_1_] denotes the initial distribution of the chain, while the transition matrices between consecutive variables are: *Q*_*j*_(*z*_*j*_|*z*_*j*−1_) = ℙ[*Z*_*j*_ = *z*_*j*_|*Z*_*j*−1_ = *z*_*j*−1_].

Since Markov chains have a well-defined sequential structure, the special class of contiguous partitions is of particular interest for generating group-knockoffs.

###### Definition 2

(Contiguous partition). *For any fixed positive integers L* ≤ *p, we call a collection of L sets* 𝒢 = (*G*_1_, …, *G*_*L*_) *a* contiguous partition *of* {1, …, *p*} *if* 𝒢 *is a partition of* {1, …, *p*} *and for any distinct G, G*′ ∈ 𝒢, *either j < l for all j* ∈ *G and l* ∈ *G*′ *or j > l for all j* ∈ *G and l* ∈ *G*′.

For example, the partition 𝒢 = (*G*_1_, …, *G*_4_) of {1, …, 10} shown on the left-hand-side of the example below is contiguous, while 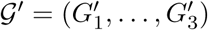, on the right-hand-side, is not.

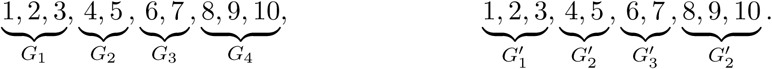

We consider only contiguous partitions when we construct knockoff copies of a Markov chain; if a given partition is not contiguous, we first refine it by splitting all non-contiguous groups.

To simplify the notation in the upcoming result, for any *g* ∈ {1, …, *L*} and associated group *G*_*g*_ ∈ 𝒢, we indicate the variables in *G*_*g*_ as: 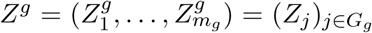. Similarly, we denote the sequence of transition matrices corresponding to the *m*_*g*_ variables contained in the *g*-th group by: 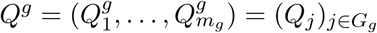. We set 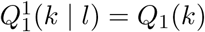 and 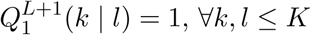.

###### Proposition 2.

*Let* 𝒢 = (*G*_1_, …, *G*_*L*_) *be a contiguous partition of* {1, …, *p*}, *such that the g-th group G*_*g*_ *has m*_*g*_ *elements. Suppose that Z is distributed as the Markov chain in* (S2), *with known parameters Q. Then, a group-knockoff copy* 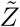, *with respect to* 𝒢, *can be obtained by sequentially sampling, for g* = 1, …, *L, the g-th group-knockoff copy* 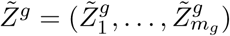 *from:*

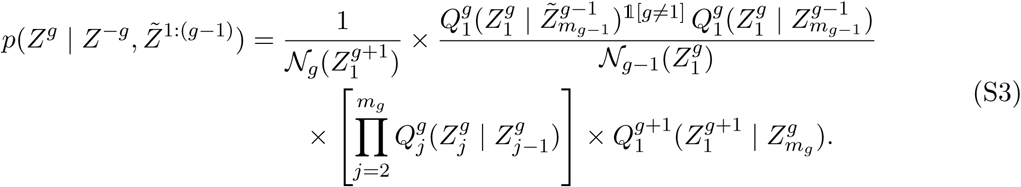

*The functions* 𝒩 *are defined recursively as:*

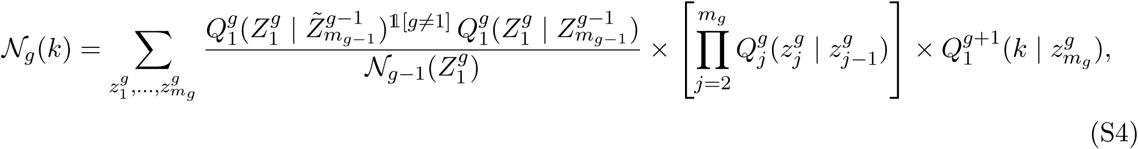

*with the convention that* 𝒩_0_(*z*) = 1 *for all z. Therefore, Algorithm 2 is an exact procedure for sampling group-knockoff copies of a Markov chain.*

###### Algorithm 2 Group-knockoffs for a Markov chain

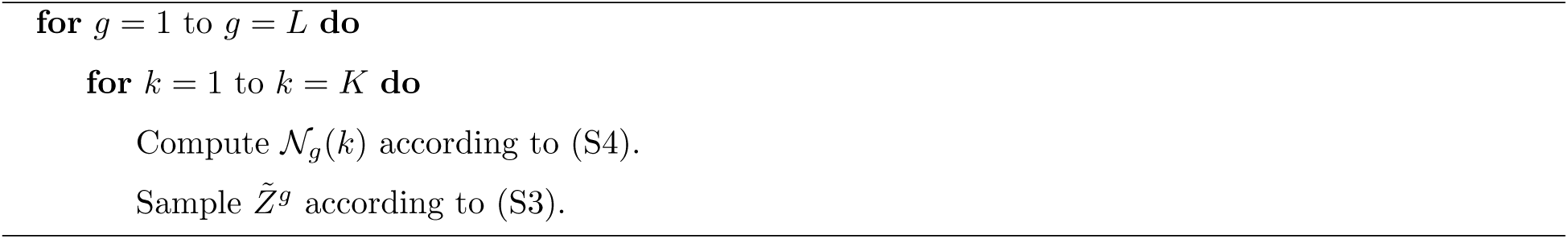

Algorithm 2 reduces to the known result for ungrouped knockoffs^2^ if *L* = *p*. In the general case, we can sample from the distribution in (S3) as follows. From (S3), we can write:

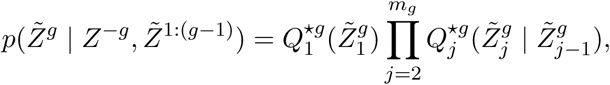

for some initial distribution 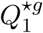 and suitable transition matrices 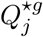. Therefore, 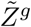 is conditionally a Markov chain. Assuming for simplicity that *g >* 1, we see from (S3) that 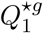 is given by:

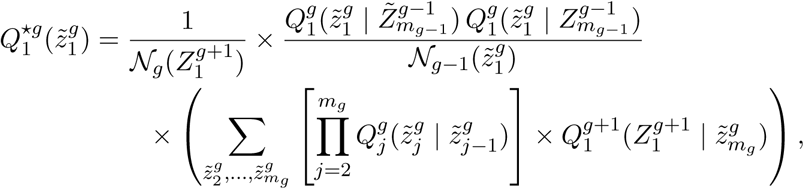

while, for *j* ∈ {2, …, *m*_*g*_},

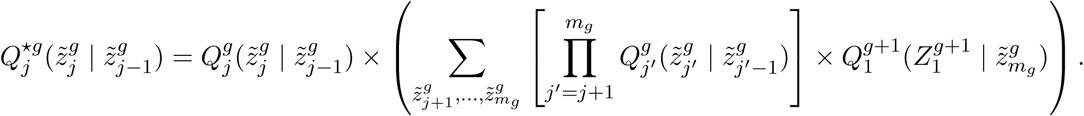

It is easy to verify that the above quantities can be computed through *m*_*g*_ − 1 multiplications of *K* × *K* matrices. Similarly, the functions in (S4) can also be computed efficiently. Therefore, the cost of sampling the *g*-th group-knockoff is 𝒪(*m*_*g*_*K*^3^), if *m*_*g*_ *>* 1, and 𝒪(*m*_*g*_*K*^2^), otherwise. The worst-case complexity of Algorithm 2 is 𝒪(*pK*^3^) because 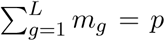. Later, we will derive a more efficient implementation in the special case of genetic variables.

##### 2. Hidden Markov models

We leverage the result in Proposition 2 to derive a construction of group-knockoffs for HMMs, similarly to previous work.^2^ We say that *X* = (*X*_1_, …, *X*_*p*_), with each variable taking values in a finite state space 𝒳, is distributed as an HMM with *K* hidden states if there exists a vector of latent random variables *Z* = (*Z*_1_, …, *Z*_*p*_), with *Z*_*j*_ ∈ {1, …, *K*}, such that:

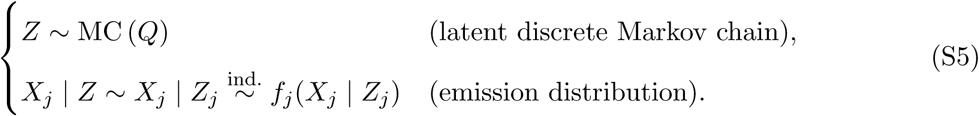

Above, MC (*Q*) indicates the law of a discrete Markov chain, as in (S2).

###### Proposition 3.

*Suppose X* = (*X*_1_, …, *X*_*p*_) *is distributed as the HMM in* (S5), *with an associated latent Markov chain Z* = (*Z*_1_, …, *Z*_*p*_). *Let* 𝒢 = (*G*_1_, …, *G*_*L*_) *be a contiguous partition of* {1, …, *p*}. *Then, Algorithm 3 generates* 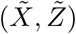 *such that:*

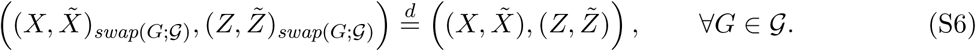

*In particular, this implies that* 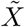 *is a group-knockoff copy of X.*

###### Algorithm 3 Group-knockoffs for an HMM

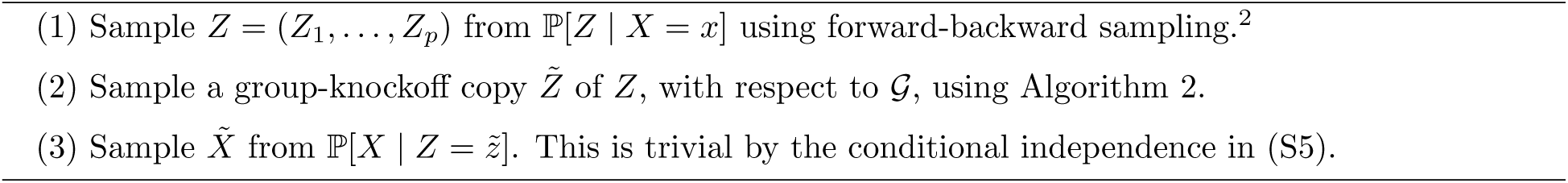

The computational complexity of the first step of Algorithm 3 is known to be 𝒪(*pK*^2^),^2^ while the worst-case cost of the second step is 𝒪(*pK*^3^). The complexity of the third step is 𝒪(*p*|𝒳|). Therefore, the worst-case total complexity of Algorithm 3 is 𝒪(*p*(*K*^3^ +|𝒳|)). Later, we will reduce this in the special case of the fastPHASE HMM by simplifying the first two steps analytically.

#### B. The fastPHASE model

##### 1. Model parameterization

We specialize Algorithm 3 in the case of the fastPHASE model.^3^ This HMM describes the distribution of genotypes as a patchwork of latent ancestral motifs. A quick preview of the results: in this Section we introduce the model; in Section S2 C we optimize the second step of Algorithm 3 to have complexity 𝒪(*pK*) for phased haplotypes, or 𝒪(*pK*^2^) for unphased genotypes; and in Section S2 D we optimize the first step of Algorithm 3 to have complexity 𝒪(*pK*) for phased haplotypes, or 𝒪(*pK*^2^) for unphased genotypes. By combining these results, we decrease the complexity of Algorithm 3 to 𝒪(*pK*) for phased haplotypes, or 𝒪(*pK*^2^) for unphased genotypes. This is an important contribution because it makes *KnockoffZoom* applicable to large datasets.

The fastPHASE model for phased haplotype sequences describes *X* = (*X*_1_, …, *X*_*p*_), with *X*_*j*_ ∈ {0, 1}, as an imperfect mosaic of *K* ancestral motifs, *D*_*i*_ = (*D*_*i*,1_, …, *D*_*i,p*_}, for *i* ∈ {1, …, *K*} and *D*_*i,j*_ ∈ {0, 1}. This can be formalized as an HMM with *K* hidden states, in the form of (S5).^3^ The transitions of the latent Markov chain are simple:

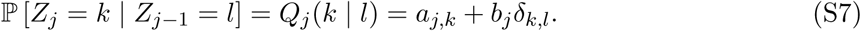

Above, *δ*_*k,l*_ indicates the Kronecker delta: *δ*_*k,l*_ is equal to 1 if *k* = *l*, and 0 otherwise. Conditional on *Z*, each *X*_*j*_ is drawn independently from:

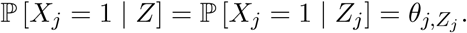

The parameters *θ* = (*θ*_*j,k*_)_*k*∈[*K*],*j*∈[*p*]_ describe the haplotype motifs in *D* and the mutation rates. We can write *a* and *b* consistent with the notation of fastPHASE^3^ as:

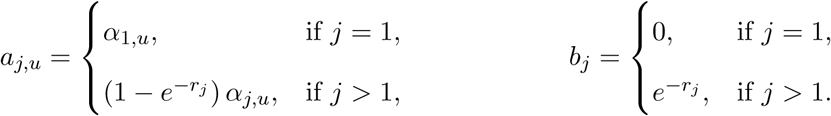

The parameters *α* = (*α*_*j,k*_)_*k*∈[*K*],*j*∈[*p*]_ describe the prevalence of each motif in the population. The likelihood of a transition in the Markov chain depends on the values of *r* = (*r*_1_, …, *r*_*p*_), which capture the genetic recombination rates along the genome. This phenomenological model of LD has inspired several successful applications for phasing and imputation.^4^

An unphased genotype sequence *X* = (*X*_1_, …, *X*_*p*_), with *X*_*j*_ ∈ {0, 1, 2}, can be described as the element-wise sum of two independent and identically distributed haplotype sequences, *H*^*a*^ and *H*^*b*^, that follow the model defined above. Consequently, *X* is also distributed as an HMM in the form of (S5) with *K*_eff_ = *K*(*K* + 1)*/*2 hidden states, where *K* is the number of haplotype motifs. This quadratic dependence on *K* follows from the fact that each latent Markov state of *X* corresponds to an unordered pair of states, 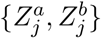, corresponding to the unobserved haplotypes. The transition probabilities for the effective Markov chain have the following structure:

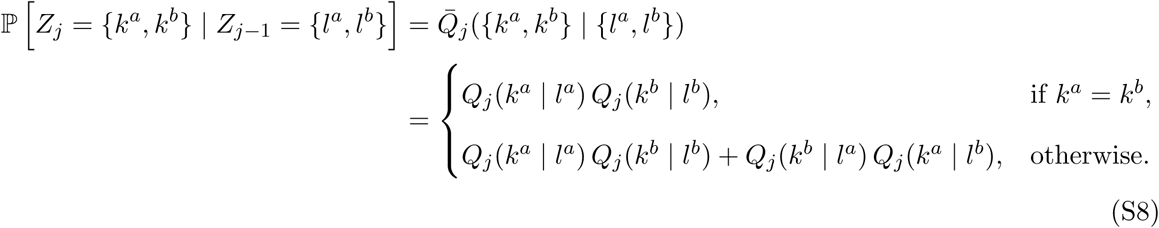

Above, *Q*_*j*_(*k* | *l*) is given by (S7) while the conditional emission distributions are:

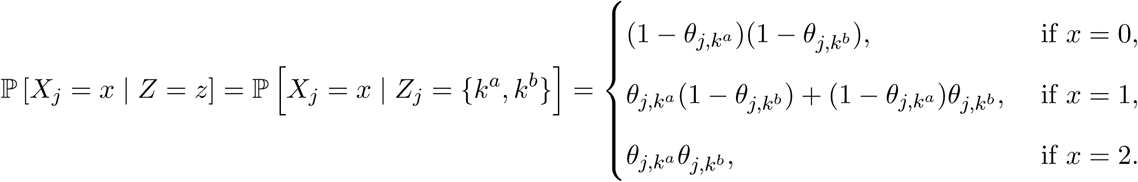

Generating group-knockoffs for genotypes using the algorithms in Section S2 would cost 𝒪(*pK*^6^), while knockoffs for haplotypes would cost 𝒪(*pK*^3^). This is prohibitive for large datasets, since the operation must be repeated separately for each subject. Before proceeding to simplify the algorithm analytically, we state the following useful lemma.

###### Lemma 1.

*The Markov chain transition matrices for the unphased genotypes in the fastPHASE HMM can be written explicitly as:*

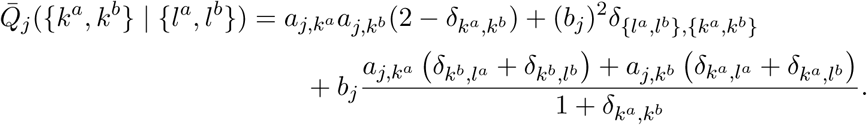

##### 2. Utilizing phased haplotypes

We leverage the phased haplotypes in the UK Biobank data^5^ to accelerate the generation of the group-knockoffs for genotypes. Denote the haplotypes of one subject as *H*^*a*^, *H*^*b*^ ∈ {0, 1}^*p*^, so that the genotypes are *X* = *H*^*a*^ + *H*^*b*^ ∈ {0, 1, 2}^*p*^. Only *X* is measures in a GWAS, whereas *H*^*a*^ and *H*^*b*^ are probabilistic reconstructions based on an HMM^4^ similar to that in Section S2 B 1. Holding this thought, note that the following algorithm generates exact group-knockoffs for the genotypes: first, sample {*H*^*a*^, *H*^*b*^} from their posterior distribution given *X*; next, create group-knockoffs 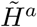 for *H*^*a*^ and 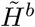 for *H*^*b*^, independently; and lastly, set 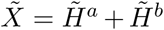. The proof is equivalent to that of Proposition 3. The first step above corresponds to phasing (although sometimes phasing is carried out by reconstructing the most likely haplotypes *H*^*a*^ and *H*^*b*^, as opposed to posterior sampling). Given the reconstructed haplotypes, the second stage of the above algorithm only involves an HMM with *K* latent states, instead of 𝒪(*K*^2^). The third stage of the algorithm is trivial.

#### C. Efficient sampling of Markov chain knockoffs

##### 1. Phased haplotypes

We begin to specialize Algorithm 2 for the HMM of phased haplotypes in Section S2 B, starting from its second step. Recall that the 𝒩 functions in (S4) are defined recursively as:

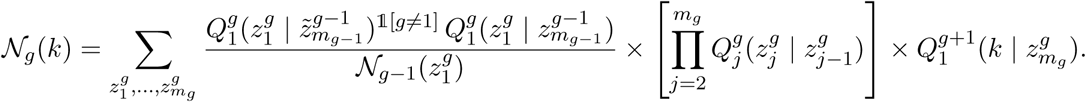

To simplify the notations, define, for each group 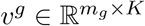 and 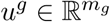 as follows:

- The last row of *v*^*g*^ and the last element of *u*^*g*^ are:

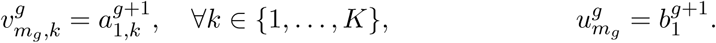
- For *j* ∈ {1, …, *m*_*g*_ − 1}, the *j*-th row of *v*^*g*^ and the *j*-th element of *u*^*g*^ are defined recursively:

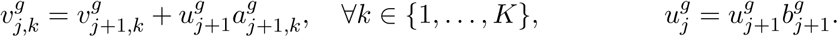

###### Proposition 4.

*For the special case of the fastPHASE model of unphased genotypes, the* 𝒩 *function in Algorithm 2 for the g-th group* (S4) *can be computed recursively as:*

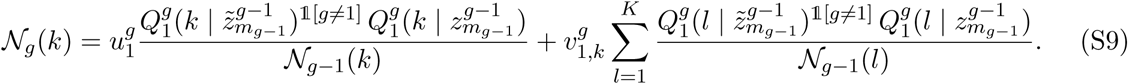

*Above, it is understood that* 𝒩_0_(*k*) = 1, ∀*k.*

Each 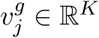 can be computed in 𝒪(*K*) time, while the additional cost for 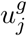 is 𝒪(1). There-fore, we compute 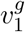 and 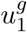 in 𝒪(*m*_*g*_*K*) time and evaluate 𝒩_*g*_(*k*) in 𝒪(*m*_*g*_*K*) time (Proposition 4).

Given 𝒩_*g*_, we must sample the vector 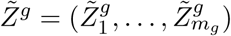 from:

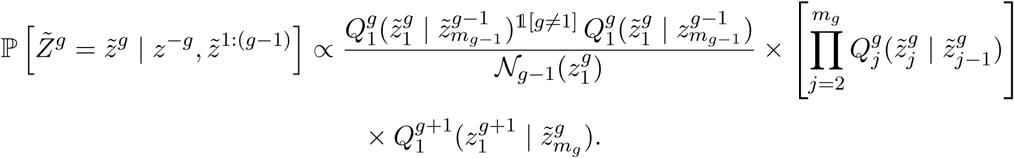

This is a multivariate distribution from which we can sample efficiently, as stated next.

###### Proposition 5.

*In the special case of Algorithm 2 for the fastPHASE model of phased haplotypes, the knockoff copy for the first element of the g-th group can be sampled from:*

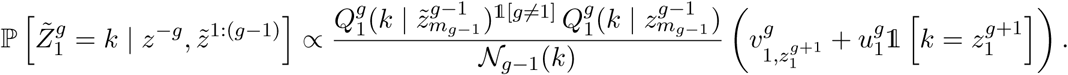

*For j* ∈ {2, …, *m*_*g*_}, *the knockoff copy for the j-th element of the g-th group can be sampled from:*

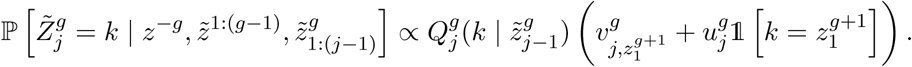

Each coordinate is sampled at an additional cost 𝒪(*K*), reusing 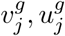 from the computation of the 𝒩 functions. The cost for the sequence is 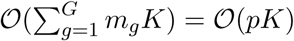, regardless of the grouping.

##### 2. Unphased genotypes

We perform analogous calculations in the case of the HMM for unphased genotypes. Here the notation is more involved. The functions in (S4) are defined recursively as:

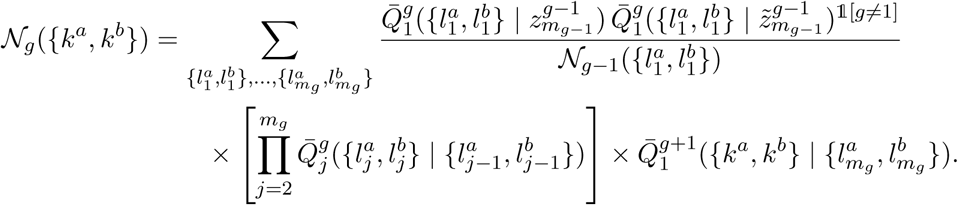

We start by defining some new variables, as in Section S2 C 1. For *k, k*^*a*^, *k*^*b*^ ∈ {1, …, *K*}, let:

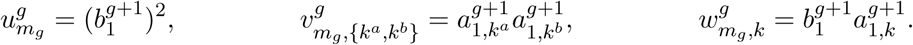

For *j* ∈ {1, …, *m*_*g*_ − 1} and *k*^*a*^, *k*^*b*^ ∈ {1, …, *K*}, we define recursively:

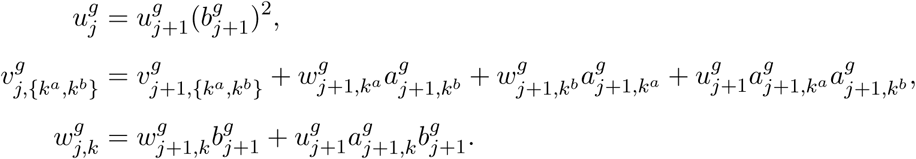

Moreover, we also define:

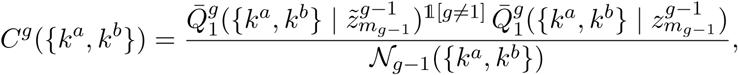

and

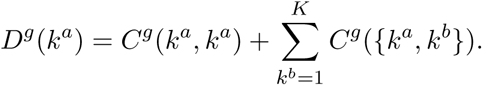

The following result shows that the cost of computing the *g*-th 𝒩 function is 𝒪(*m*_*g*_*K*^2^).

###### Proposition 6.

*For the special case of the fastPHASE model of unphased genotypes, the* 𝒩 *function in Algorithm 2 for the g-th group is given by:*

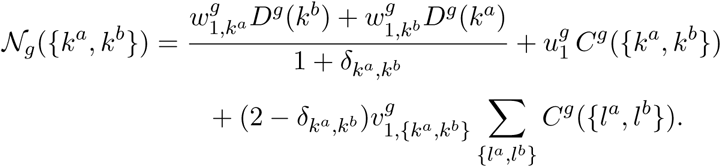

The sampling part of Algorithm 2 is similar to that in Section S2 C 1.

###### Proposition 7.

*For the special case of the fastPHASE model of unphased genotypes, the knockoff copy for the first element of the g-th group can be sampled in Algorithm 2 from:*

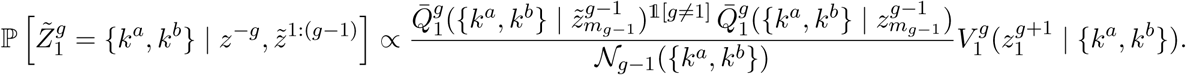

*The knockoff for the j-th element of the g-th group, for j* ∈ {2, …, *m*_*g*_}, *can be sampled from:*

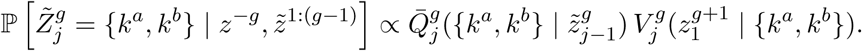

*Above, the variables* 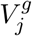 *are defined as:*

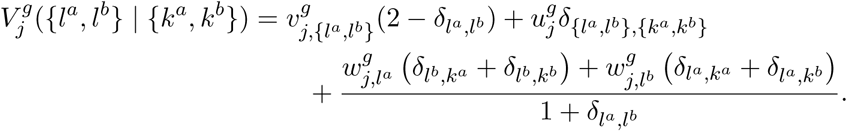

Sampling group-knockoffs costs 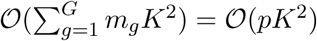 per individual, with any grouping.

#### D. Efficient forward-backward sampling

##### 1. Forward-backward sampling for HMMs

The first step in Algorithm 3 consists of sampling *Z* from the posterior distribution of the latent Markov chain in (S5) given *X*. In general, this requires a variation of Viterbi’s algorithm,^2^ as recalled in Algorithms 4 and 5. Here, *K* indicates the number of states in the Markov chain.

###### Algorithm 4 Forward-backward sampling (forward pass)

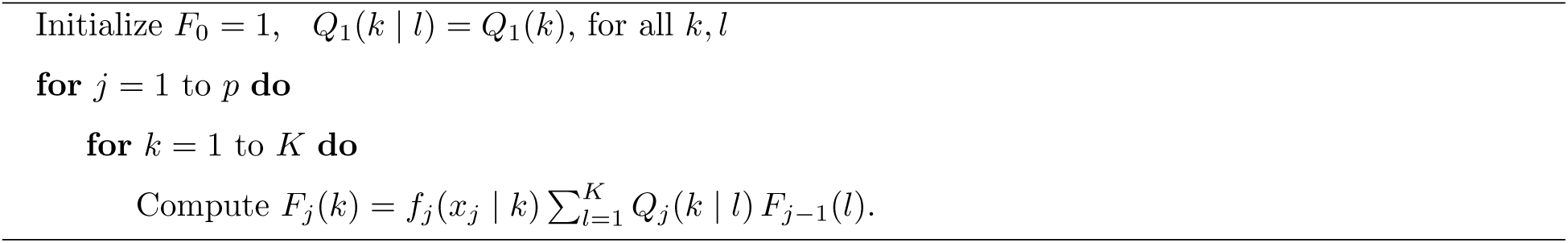

###### Algorithm 5 Forward-backward sampling (backward pass)

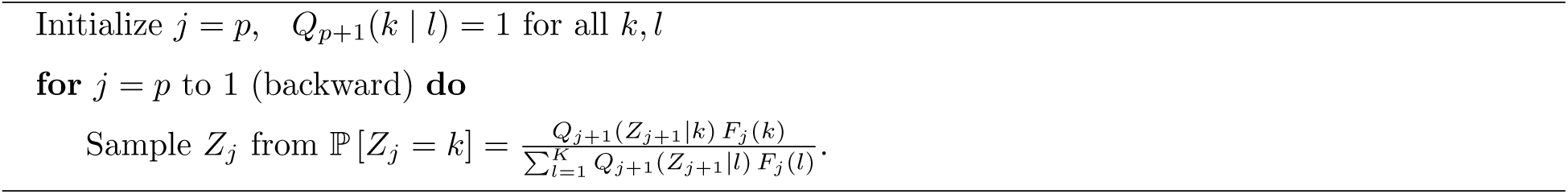

##### 2. Phased haplotypes

The forward probabilities in Algorithm 4 are defined recursively, for *j* ∈ {2, …, *p*}, as:

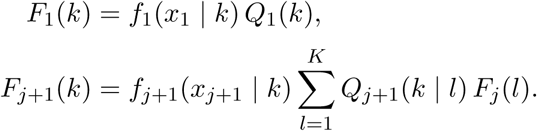

The cost of computing all forward probabilities is generally 𝒪(*pK*^2^) but it can be reduced to 𝒪(*pK*) using the symmetries in (S7). Earlier instances of the same idea can be found in the statistical genetics literature,^6^ although we prefer to present the full results here for completeness, especially since they are easy to present using the notation developed in Section S2 C.

###### Proposition 8.

*In the special case of the fastPHASE model of phased haplotypes, the forward probabilities in Algorithm 4 can be computed recursively as follows:*

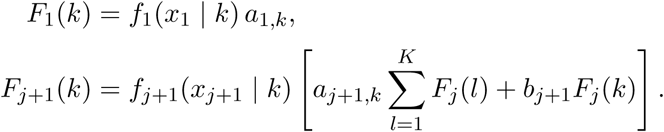

Above, the sum only needs be computed once because it does not depend on the value *k*. Consequently, we can implement the first part of Algorithm S2 A 2 at cost 𝒪(*pK*).

##### 3. Unphased genotypes

We can also implement Algorithm 4 efficiently for unphased genotypes. For *j* ∈ {2, …, *p*}, the forward probabilities are defined as:

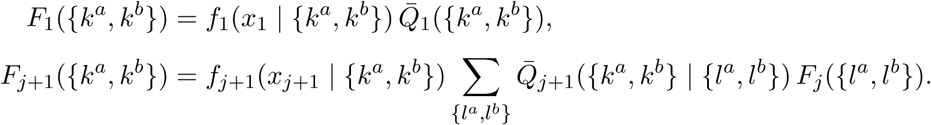

This computation generally costs 𝒪(*pK*^4^) because the number of discrete states in the latent Markov chain is quadratic in the number of haplotype motifs.

###### Proposition 9.

*In the special case of the fastPHASE model of unphased genotypes, the forward probabilities in Algorithm 4 are given by the following recursive formula:*

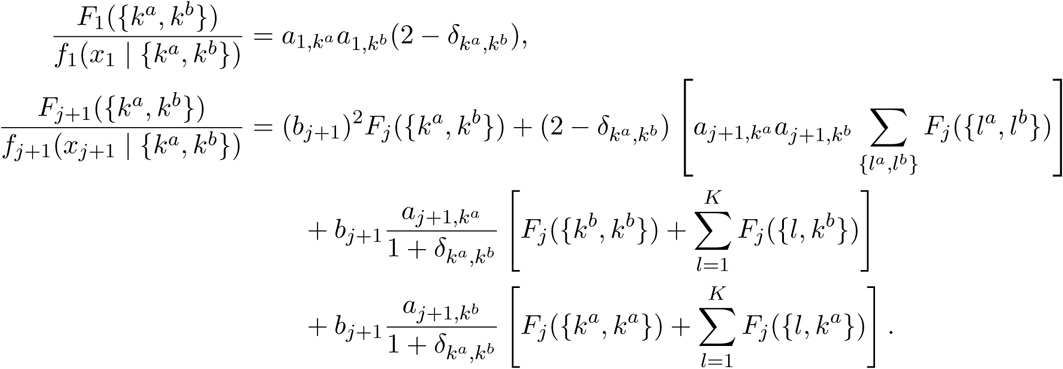

This gives us a 𝒪(*pK*^2^) algorithm because the above sums can be efficiently pre-computed.

### S3. TECHNICAL DETAILS

#### A. Schematic

**FIG. S1.**
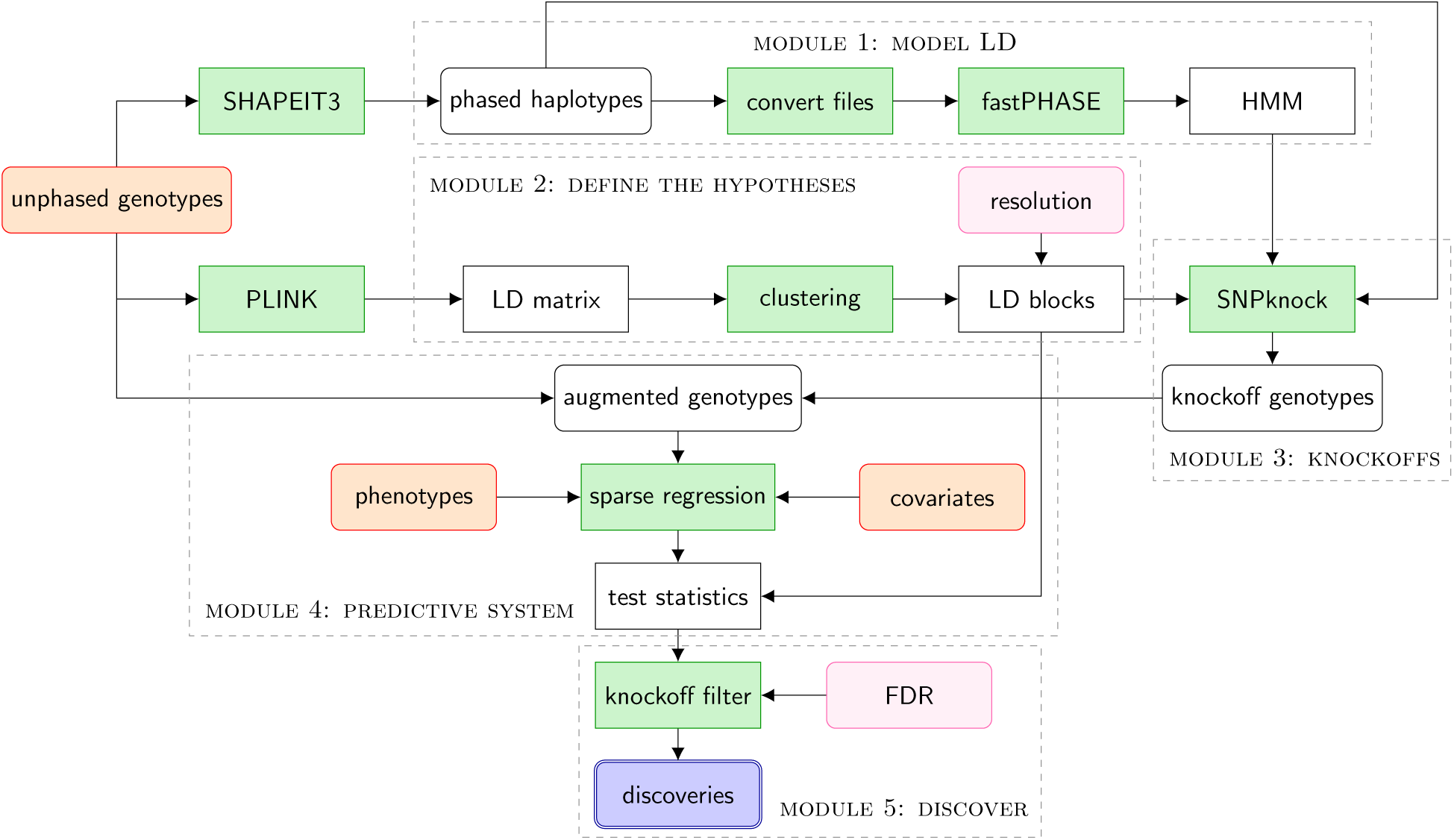
Schematic of the *KnockoffZoom* method for the analysis of GWAS data. The inputs are the unphased genotypes (or the phased haplotypes, if available), the phenotypes, and any relevant covariates (e.g., age, sex, other demographics, principal components). The user chooses the resolution at which our method performs genome-wide fine-mapping and the nominal FDR level. In the first module, an HMM is fit using the phased haplotypes using fastPHASE. Meanwhile, the typed loci are assigned to blocks that represent our units of inference (second module), based on the observed LD, the physical locus positions and the resolution. Then, the estimated HMM and the partition of the variants are used by our software (SNPknock) to generate the knockoffs (third module). In the fourth module, a test statistic for each block is computed by a machine-learning system that predicts the phenotype and estimates the feature importance of genotypes and knockoffs, without knowing which one is which. The knockoffs serve as negative controls, allowing the filter to calibrate the statistics for FDR control (fifth module).

#### B. Computational and memory resources

The computation time required by *KnockoffZoom* for the data analysis in this paper is reported in Table S1, for each operation described in the flowchart of Figure S1. The entire procedure, starting from phased haplotypes, takes about 12 days—this is less than the time needed for phasing.^5^ In principle, our procedure can be applied to unphased genotypes, although this is not recommended if the dataset is very large, for the knockoff generation would be slower. If we want to analyze different phenotypes, only the fourth and fifth modules need to be repeated (the latter requires negligible resources). The analysis of a new trait for the same individuals in the UK Biobank takes us less than 12 hours with an efficient implementation of the fourth module. As a comparison, BOLT-LMM takes between a few days and a week, depending on the phenotype. These are rough upper bounds because the exact time depends on the computing hardware and other factors.

**TABLE S1.** Computation time and resources for each module of *KnockoffZoom*, for the analysis of the phenotype *height* in the UK Biobank (591k SNPs and 350k individuals).

| Module | Operation | Time (h) | Memory (GB) | Machines | Cores |
| --- | --- | --- | --- | --- | --- |
| 1 | convert haplotypes | 30 | 50 | 22 | 1 |
| 1 | fit HMM | 144 | 50 | 22 | 1 |
| 2 | compute LD matrix | 2 | 50 | 22 | 1 |
| 2 | hierarchical clustering | 8 | 200 | 22 | 1 |
| 3 | generate knockoffs | 72 | 30 | 22 | 1 |
| 3 | combine augmented data | 8 | 60 | 1 | 1 |
| 4 | memory mapping | 12 | 60 | 1 | 1 |
| 4 | sparse regression | 12 | 20 | 1 | 10 |

The resource requirements of our method are also summarized in Table S1. The computations in modules 1–3 are divided between 22 machines, one for each autosome (our estimates refer to the longest one). The memory footprint is low, except for clustering, which requires about 200GB. If this becomes limiting, one can define the LD blocks differently, e.g., locally and concatenating the results. By comparison, BOLT-LMM requires approximately 100 GB with the same data.

We can easily analyzed the theoretical complexity of the first and third modules. In order to efficiently fit an HMM for the genotypes, the haplotypes are phased using SHAPEIT3;^5^ then fastPHASE^3^ is applied to the inferred haplotypes. The computational complexity of SHAPEIT3 and fastPHASE are 𝒪(*pn* log *n*) and 𝒪(*pnK*), respectively, where *p* is the number of variants, *n* is the number of individuals and *K* is the number of haplotype motifs in the HMM. Our analytical calculations for the fastPHASE HMM have reduced the cost of the third module to 𝒪(*pnK*). If the phased haplotypes are not available, a modified version of our algorithm has cost 𝒪(*pnK*^2^). The second and fourth modules are more difficult to analyze theoretically, as different choices of tools are available to cluster the variants and compute the test statistics. For the latter, we rely on very fast and memory-efficient implementations of sparse linear and logistic regression.^7^ The fifth and final module of *KnockoffZoom* is computationally negligible.

#### C. Computing the test statistics

The feature importance measures that we have adopted to define the test statistics of *KnockoffZoom* are based on sparse multivariate linear and logistic regression, since Bayesian and penalized regression approaches are currently the state-of-the-art for predicting complex traits.^8,9^ However, our methodology could easily incorporate other scalable machine learning tools, like SVMs, random forests, and deep learning.^10–12^ In principle, the test statistics of *KnockoffZoom* could also be computed using cheaper single-marker methods, e.g., by contrasting the univariate regression p-values for the original genotypes and the knockoffs; however, we do not recommend this for two reasons. First, our method is more powerful if used in combination with a multi-marker predictive model that explains a larger fraction of the variance in the phenotypes. Second, multivariate methods are less susceptible to the confounding effect of population structure.^13^ Therefore, they improve our robustness against possible violations of the modeling assumptions, i.e., the ability of the HMM to correctly describe the real distribution of the genotypes.

In this paper, we use the R packages `bigstatsr` and `bigsnpr`^7^ to fit a sparse generalized linear model the trait given the augmented matrix of explanatory variables 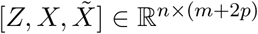. Here, *Z* is a matrix containing the covariates for all individuals, while *X* and 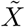 indicate the observed genotypes and their knockoff copies, respectively. The regularization penalty is tuned using a modified form of 10-fold cross-validation. The regression coefficients for each variable are averaged over the 10 folds, finally obtaining an estimate of the quantity 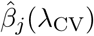 from the Online Methods B. Since the knockoff-augmented genotype data is stored on an hard-drive as a memory-mapped file, the algorithm is memory efficient.^14^ In addition, if the knockoffs in a group *g* are known in advance to have very little power as negative controls, e.g., 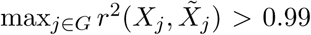, the corresponding hypothesis is automatically discarded by setting *W*_*g*_ = 0. This operation is independent of the phenotypes and preserves the symmetry required to control the FDR,^15^ but it may improve power, especially at high resolution when the pairwise *r*^2^ can be high (Section S4 B).

It is important to remark that even though the order in which the variables are provided to the black box that computes the feature importance measures (e.g., the lasso) does not matter in theory, it can make a difference in practice. For instance, numerical instabilities may induce the black box to give an unfair advantage to variables found earlier in the input, which would result in a symmetry break if the input is 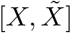, possibly leading to a loss of FDR control. For this reason, our implementation randomly swaps each pair of genotypes and knockoffs before computing the variable importance measures. The original identity of the knockoffs is only revealed later to determine the sign of the test statistics. Therefore, the FDR guarantee of *KnockoffZoom* is completely immune to such numerical instabilities.

#### D. Implementation of the LMM oracle

The FDR-controlling oracle used throughout our simulations is implemented as follows. Significant loci are clumped by applying the PLINK algorithm to the LMM p-values, over a discrete grid of possible values for the primary significance threshold. We use a wide logarithmic scale: 5 × 10^−9^, 5 × 10^−8^, 5 × 10^−7^, 5 × 10^−6^, 10^−5^, 2 × 10^−5^, 5 × 10^−5^, 10^−4^, 2 × 10^−4^, 5 × 10^−4^, 10^−3^. The number of discoveries and the true FDP are computed for each threshold and experiment. To mitigate the discretization, we interpolate the results between the grid points. Then, we count the true discoveries corresponding to the most liberal threshold that controls the FDP (or the FDR, if the experiments are replicated multiple times with the same parameters).

### S4. NUMERICAL SIMULATIONS

#### A. Goodness-of-fit of the HMM

We fit the HMM of Section S2 B to the phased haplotypes for the 350k individuals in the UK Biobank retained in our analysis. The parameters of 22 separate models are estimated applying fastPHASE separately within each autosome. The flexibility of the HMM is controlled by the number *K* of haplotype motifs, which is important for the performance of *KnockoffZoom*. If *K* is too small, the knockoffs may not have the right LD structure to serve as negative controls, resulting in an excess of false discoveries; if *K* is too large, over-fitting may render our procedure overly conservative and reduce power. In order to choose a good value of *K*, we consider a wide range of alternatives and evaluate the goodness-of-fit of each model in terms of its ability to correctly predict missing values in a hold-out sample. For this purpose we divide the individuals into a training set of size 349,119 and a validation set of size 10,000; then, we mask 50% of the haplotypes from the second set and use the fitted HMMs to reconstruct their value given the observed data. The goodness-of-fit is thus measured in terms of imputation error: lower values indicate a more accurate model. The results corresponding to chromosome 22 are shown in Table S2. Even though *K* = 100 seems optimal according to this metric, little improvement is observed above 50. Therefore, we choose *K* = 50 to generate knockoffs for the rest of the analysis, in the interest of computational speed. Finally, we verify that the goodness-of-fit of this HMM with *K* = 50 is significantly better than that of a multivariate Gaussian approximation of the genotype distribution,^15^ with parameters estimated on all 359k samples, which has imputation error equal to 5.52%.

**TABLE S2.** Out-of-sample imputation error of our HMM for haplotypes missing at random, as a function of the number *K* of motifs in the model.

| K | 1 | 2 | 5 | 10 | 15 | 20 | 30 | 50 | 75 | 100 |
| --- | --- | --- | --- | --- | --- | --- | --- | --- | --- | --- |
| Imputation error (%) | 14.59 | 10.23 | 6.74 | 5.57 | 5.04 | 4.77 | 4.4 | 4.01 | 3.75 | 3.71 |

#### B. Resolution and locus partitions

We apply *KnockoffZoom* at 7 levels of resolution. Each level corresponds to a specific partition of the 591k loci into contiguous LD blocks, as summarized in Table S3. At high resolution, the intrinsic difficulty of fine-mapping is reflected by the greater similarity between the original genotypes and their knockoff copies. In this setting, we need many samples or large effect sizes to distinguish a statistically significant contrast between the predictive power of genotypes and knockoffs. Conversely, knockoffs and genotypes have a smaller *r*^2^ at lower resolutions. The full distribution of these *r*^2^ coefficients at different resolutions is shown in the boxplots of Figure S2.

**TABLE S3.**
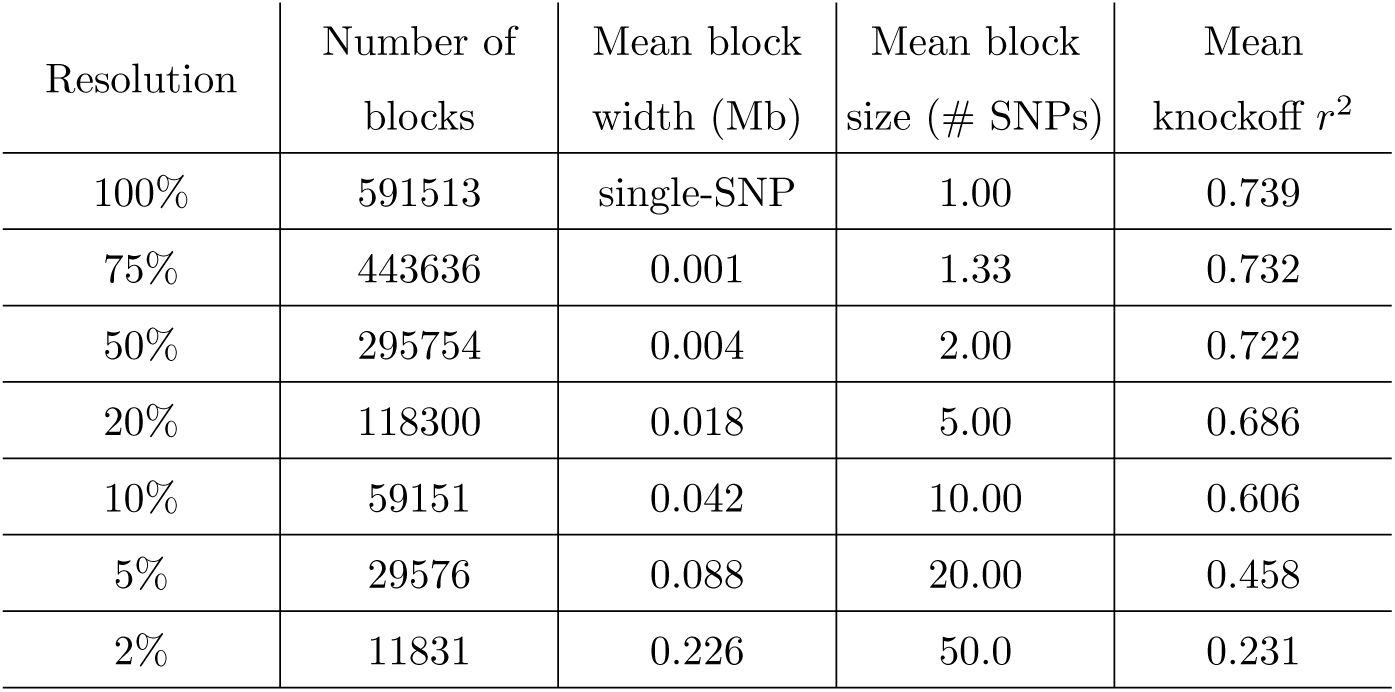
Summary of the genome partitions at different resolutions. The last column indicates the *r*^2^ between genotypes and knockoffs at each resolution.

| Resolution | Number of blocks | Mean block width (Mb) | Mean block size (# SNPs) | Mean knockoff $r^2$ |
| --- | --- | --- | --- | --- |
| 100% | 591513 | single-SNP | 1.00 | 0.739 |
| 75% | 443636 | 0.001 | 1.33 | 0.732 |
| 50% | 295754 | 0.004 | 2.00 | 0.722 |
| 20% | 118300 | 0.018 | 5.00 | 0.686 |
| 10% | 59151 | 0.042 | 10.00 | 0.606 |
| 5% | 29576 | 0.088 | 20.00 | 0.458 |
| 2% | 11831 | 0.226 | 50.0 | 0.231 |

**FIG. S2.**
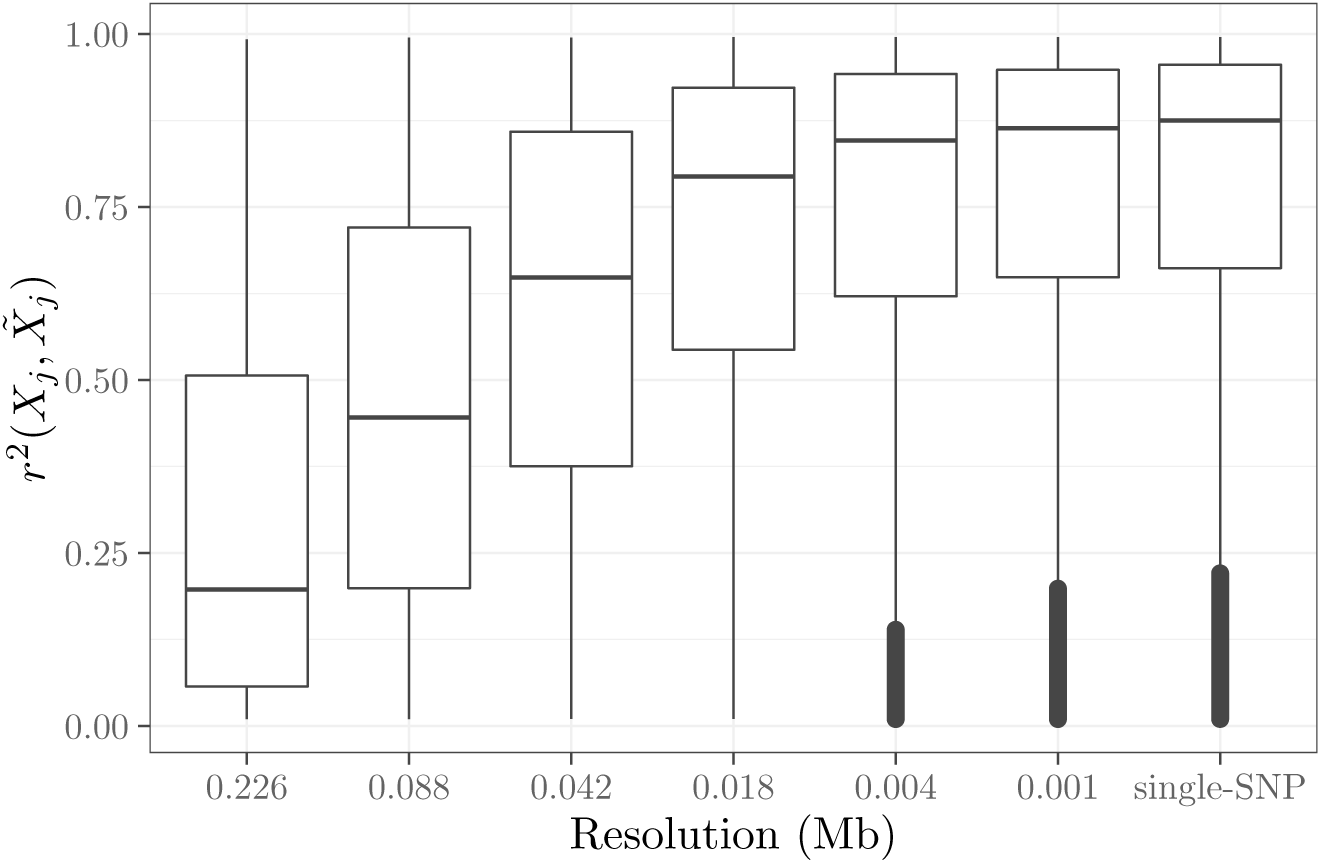
Distribution of *r*^2^ between genotypes and knockoffs at each resolution.

#### C. Exchangeability diagnostics and long-range correlations

Having generated the knockoffs with the estimated HMM discussed above, it is interesting to verify their exchangeability with the real data. In theory, if the distribution of genotypes followed this HMM exactly, the joint distribution of 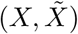 would be unchanged when {*X*_*j*_ : *j* ∈ *G*} is swapped with 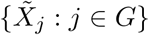, for any group *G* ∈ 𝒢, by construction of 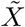. However, since our HMM can only approximate the true distribution of genotypes in the UK Biobank, this exchangeability is not perfect in practice. A simple way to quantify and visualize this exchangeability is to compute the covariance of the augmented matrix of explanatory variables 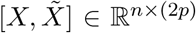. For instance, we compare in the left-hand-side of Figure S3 *r*^2^(*X*_*j*_, *X*_*k*_) with 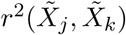, for SNPs *j, k* in different groups on chromosome 22. In the right-hand-side of Figure S3, we also compare *r*^2^(*X*_*j*_, *X*_*k*_) with 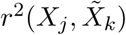, for the same pairs of SNPs. In both scatter plots, the points would concentrate around the 45° line if the HMM were exact.

We observe that the desired exchangeability holds approximately but some deviations occur, especially for lower-frequency variants (see Figure S4), and when the knockoffs are constructed using low-resolution partitions. This should not be very surprising. First, empirical correlations involving lower-frequency variants are naturally noisier, and it is natural that the HMM fits better the distribution of more common variants. Second, the accuracy of knockoffs at low resolution depends on our ability to capture long-range correlations, which are generally weaker compared to short-range LD, but also slightly underestimated by the current implementation of our HMM. In fact, long-range correlations may be partially due to some underlying population structure that we are not explicitly trying to model.

The methods presented in this paper can naturally accommodate a more flexible implementation of the HMM that describes long-range correlations and population structure accurately; this extension will be presented soon in a separate work. For the time being, we have verified that the knockoffs described here lead to FDR control in practice, across a variety of numerical simulations involving the same real genotypes used in the analysis of the unrelated British individuals in the UK Biobank. Finally, we have verified that *KnockoffZoom* tends to select more common variants for all real phenotypes analyzed in this paper (Section S5 G); thus it does not seem to be practically affected by the lower accuracy of knockoffs for rarer variants. However, it may be of particular interest to focus on studying rarer variants in future applications. Therefore, we are also working to improve the current algorithm used to estimate the HMM parameters in order to model their distribution more closely.

**FIG. S3.**
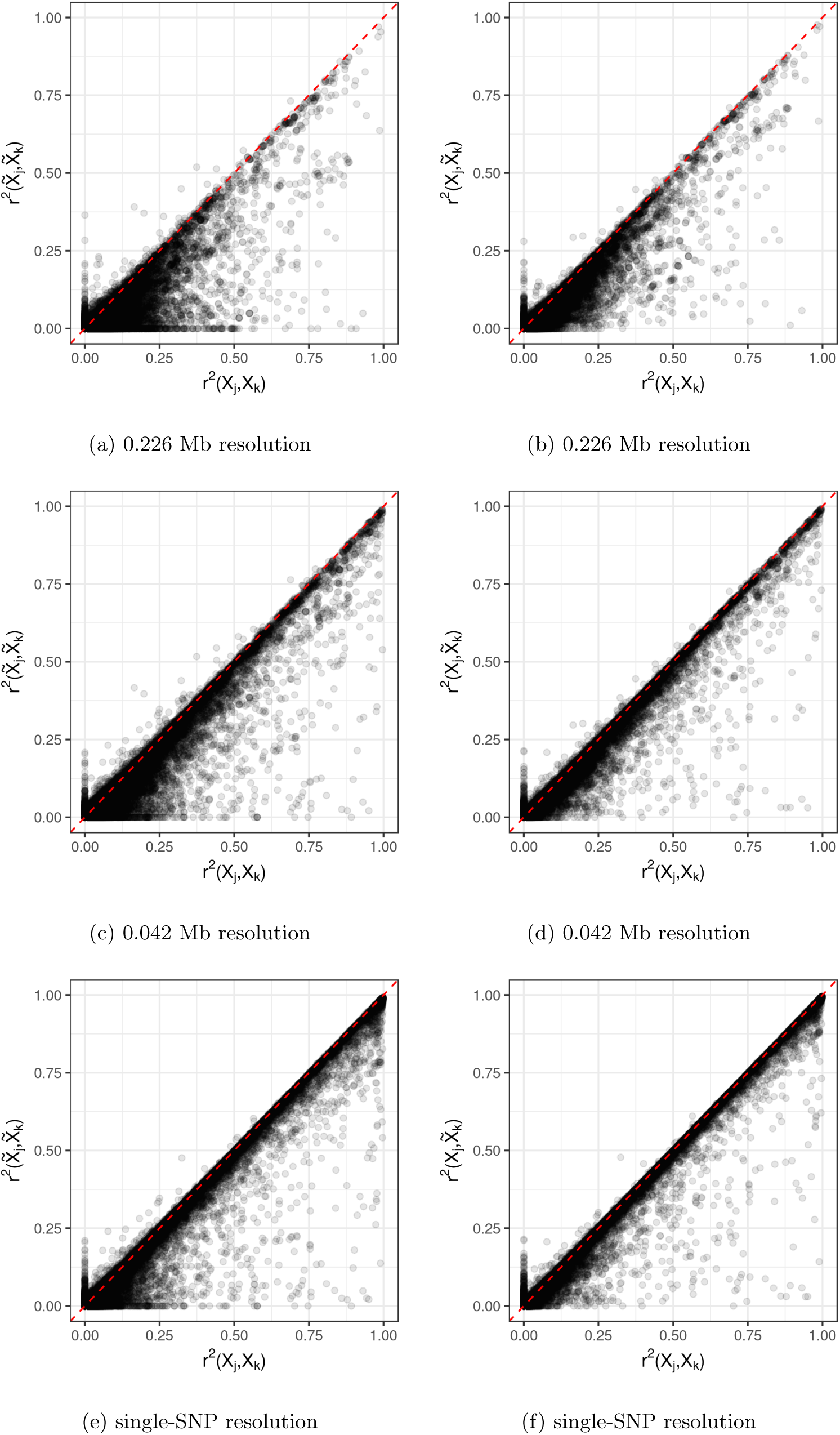
Correlation exchangeability diagnostics for knockoffs at different resolutions.

**FIG. S4.**
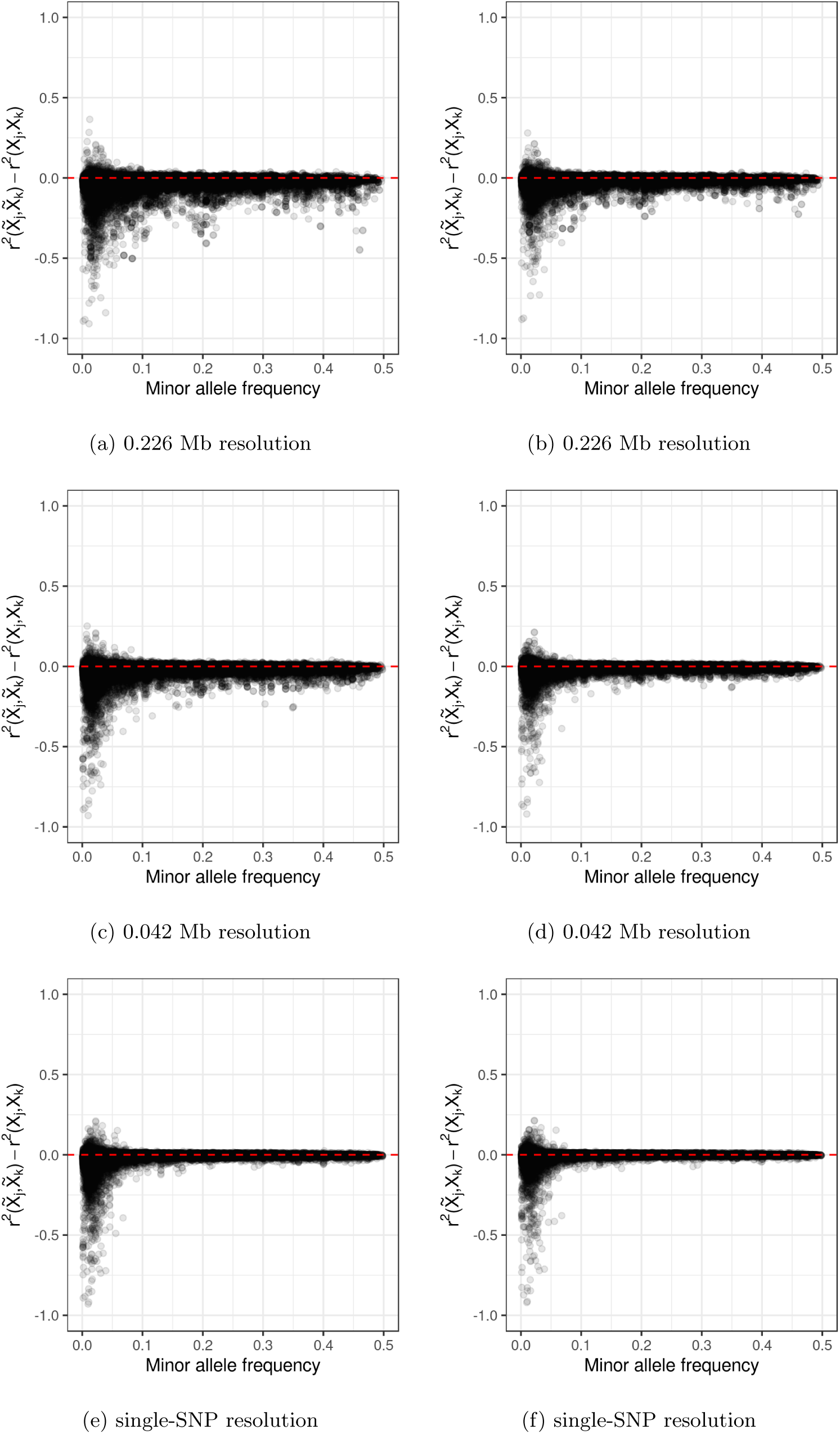
Deviations from their ideal values of the knockoff exchangeability diagnostics in Figure S3, as a function of the smallest minor allele frequency for each pair of variants.

#### D. Genetic architecture of the synthetic phenotypes

We generate synthetic phenotypes with a variety of genetic architectures in order to test the performance of our method in controlled environments, as summarized in Table S4.

**TABLE S4.** Genetic architectures used in the numerical simulations. The first architecture is used in the main numerical experiments in the paper, while the simulated case-control study is based on the second one. The other architectures are considered in the additional experiments presented in this supplement.

| Name | Number of causal regions | Causal variants per region | Total number of causal variants | Width of causal regions (Mb) | Figures |
| --- | --- | --- | --- | --- | --- |
| 1 | 500 | 5 | 2500 | 0.1 | 2–4, S5, S7–S11, S14 |
| 2 | 500 | 1 | 500 | 0.1 | 6 |
| 3 | 100 | 1 | 100 | 0.1 | S12 |
| 4 | 100 | 5 | 500 | 0.1 | S12 |
| 5 | 1000 | 1 | 1000 | 0.1 | S6, S8, S12 |
| 6 | 1000 | 2 | 2000 | 0.1 | S12 |
| 7 | 1000 | 5 | 5000 | 0.1 | S12 |
| 8 | 1000 | 10 | 10000 | 0.1 | S12 |

#### E. The difficulty of controlling the FDR

Targeting the FDR in a GWAS is difficult; in particular, it is challenging to control it over distinct (conditional) discoveries using LMM (marginal) p-values. We illustrate the challenges by considering two variations of the Benjamini-Hochberg (BH) procedure.^16^ First, we apply the BH correction to the BOLT-LMM p-values and report discoveries with the usual PLINK clumping. Second, we use BOLT-LMM to test pre-determined hypotheses defined over the *KnockoffZoom* LD blocks. For this purpose, we summarize the p-values corresponding to SNPs in the same block into a single number, a Simes p-value,^17^ that we provide to the BH procedure without further clumping. For example, we consider two resolutions: 0.226 Mb or 0.018 Mb-wide blocks, on average.

The observed outcome is an excess of false positives, regardless of how we count distinct discoveries; see Figure S5. With the first approach, the FDR is inflated because the BH procedure counts the discoveries *pre-clumping*, while the FDR is evaluated on findings that are defined differently, *post-clumping*.^18^ Moreover, these p-values are only designed for marginal hypotheses, while we are interested in distinct (conditional) findings.^19^ With the second approach, the FDR would be controlled if the p-values for loci in different groups were correct and independent. However, this is not the case, even when the blocks are large, because some inter-block LD remains. These examples are not exhaustive, but they illustrate the fundamental problem that the LMM p-values are not calibrated for conditional hypotheses, as already shown in Section II D 2.

**FIG. S5.**
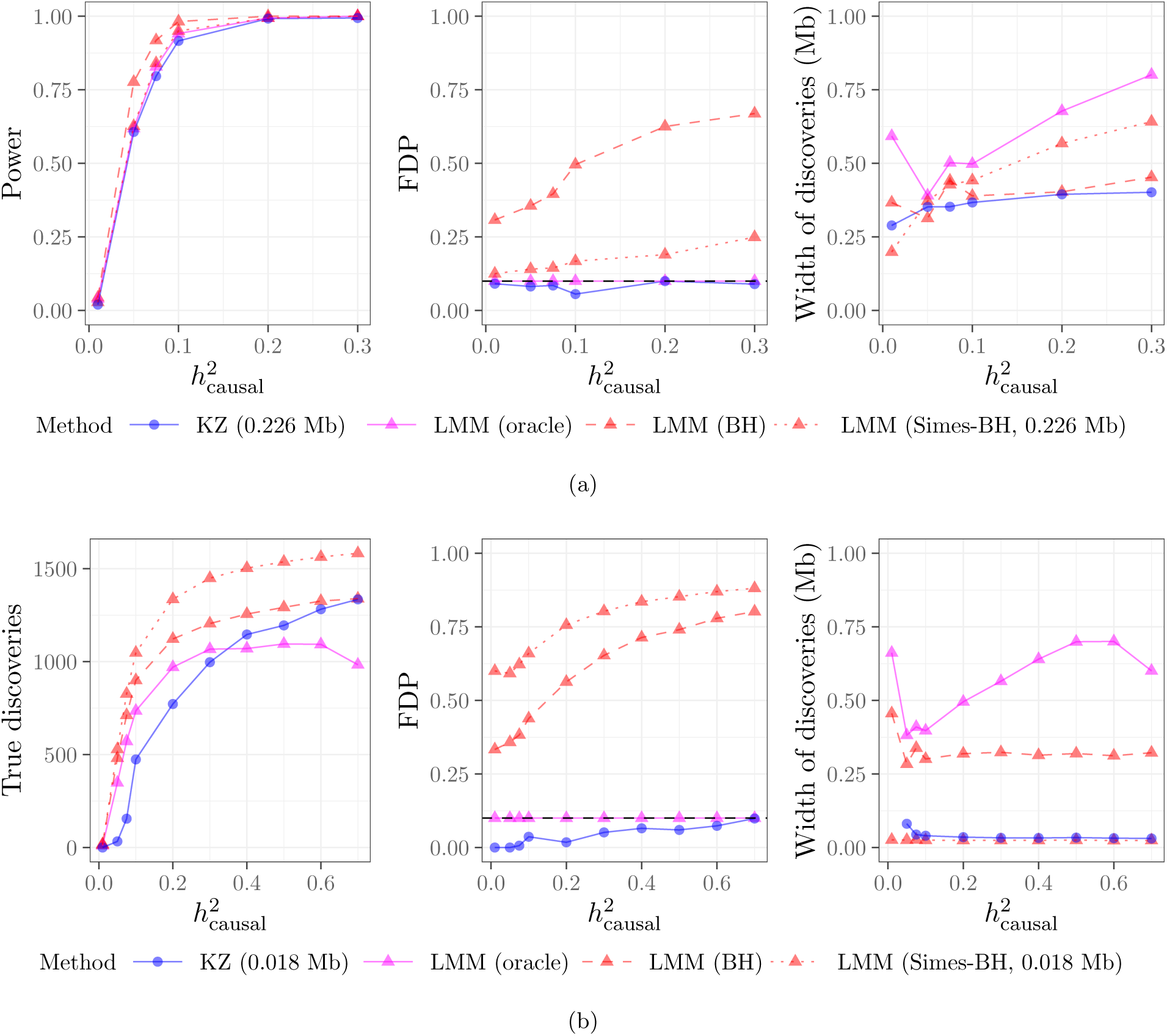
Performance of *KnockoffZoom* and two alternative LMM-based heuristics for FDR control, for the same experiments as in Figure 3. In (b) we do not simplify the *KnockoffZoom* findings at multiple resolutions; instead, we only report those at a fixed resolution, for comparison with the Simes-BH method.

A variation of the BH procedure has been recently proposed in combination with clumping,^18^ but it does not address conditional hypotheses. An alternative solution is based on the distributions of scan statistics,^19^ although we are not aware of whether this has been tested. A different approach based on penalized regression^20^ has shown promise, but it relies on stringent modeling assumptions.

#### F. Locus discovery

We complement in Figure S6 the simulations in Figure 3 by considering the fifth architecture from Table S4, in which all causal SNPs are well-separated. We observe in Figure S6 (b) that, if the signals are weak, it is difficult to reject the high-resolution conditional hypotheses. If the signals are sufficiently strong, we separate nearby casual variants and make additional distinct discoveries.

**FIG. S6.**
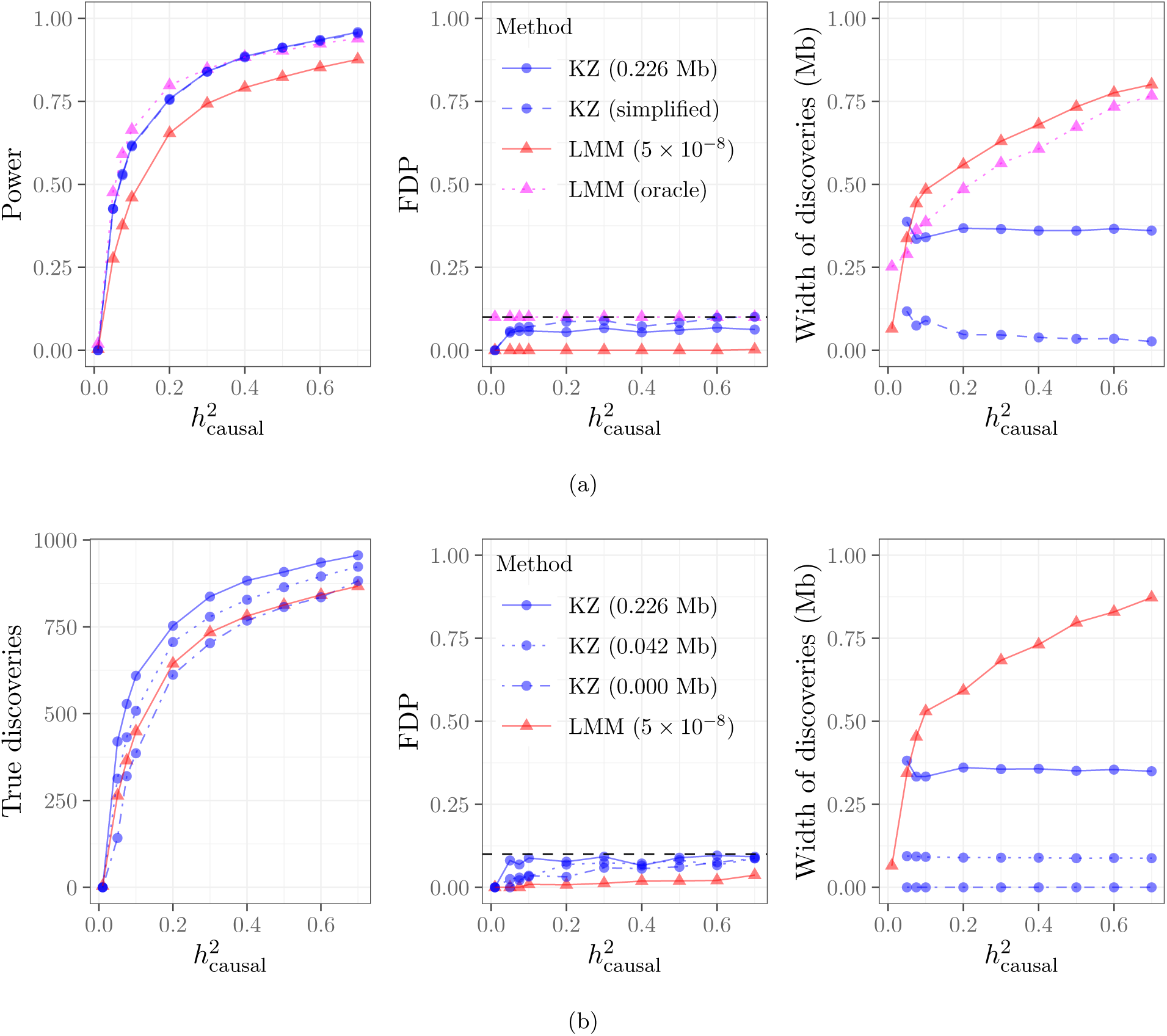
Performance of *KnockoffZoom* and BOLT-LMM for a simulated trait with genetic architecture 5 from Table S4. (b): *KnockoffZoom* at different levels of resolutions. Other details as in Figure 3.

#### G. Fine-mapping

##### 1. Additional details on fine-mapping

CAVIAR represents a broader class of popular tools, including FINEMAP,^21^ PAINTOR,^22^ and many others^23^ that we do not consider explicitly in this paper. It has recently been pointed out that these procedures cannot distinguish between multiple causal variants that may be present within the genomic region under analysis,^24^ and SUSIE was proposed to overcome this limitation. SUSIE is based on Bayesian step-wise regression and uses the original data instead of relying only on summary statistics from locus discovery.^25^ However, neither CAVIAR nor SUSIE are designed to be applied genome-wide for the analysis of complex traits due to computational and statistical reasons, since they rely on relatively simple models with a small number of distinct causal effects. Therefore, we apply them locus-by-locus in a two-step procedure starting with BOLT-LMM.

The two-step procedure is calibrated to target a nominal error rate that is comparable to the FDR of *KnockoffZoom*. Regarding SUSIE, we simply apply it with nominal coverage parameter equal to 90%, so that at most 10% of its reported findings are expected to be false positives. In the case of CAVIAR, the solution is similar but it requires a more careful explanation. Under certain modeling assumptions, CAVIAR is designed to control the probability that any causal variants are erroneously discarded from the input candidate set^25^ (but it does not tell apart distinct causal effects). Therefore, assuming that the input always contains at least one causal SNP (as we ensure in our simulations by applying BOLT-LMM with aggressive clumping), CAVIAR indirectly targets, within the two-step procedure, a notion of FDR similar to that of SUSIE and *KnockoffZoom*. In particular, setting the control parameter of CAVIAR equal to 90%, we can expect that at most 10% of the reported findings do not contain at least one causal variant.

##### 2. Additional simulations under different settings

A downside of the two-step paradigm of fine-mapping is that it may sometimes introduce selection bias, as shown in Figure S7. The biased results refer to SUSIE applied exactly on the clumps reported by BOLT-LMM, without including nearby unselected SNPs.

We complement the results in Figure 4 by considering the fifth architecture from Table S4, as in Figure S6. Here, *KnockoffZoom* is more powerful than SUSIE. Note that the output of SUSIE needs to be simplified because it occasionally reports overlapping subsets of SNPs; therefore, we consolidate those that are not disjoint and retain the smallest if one is included in the other.

**FIG. S7.**
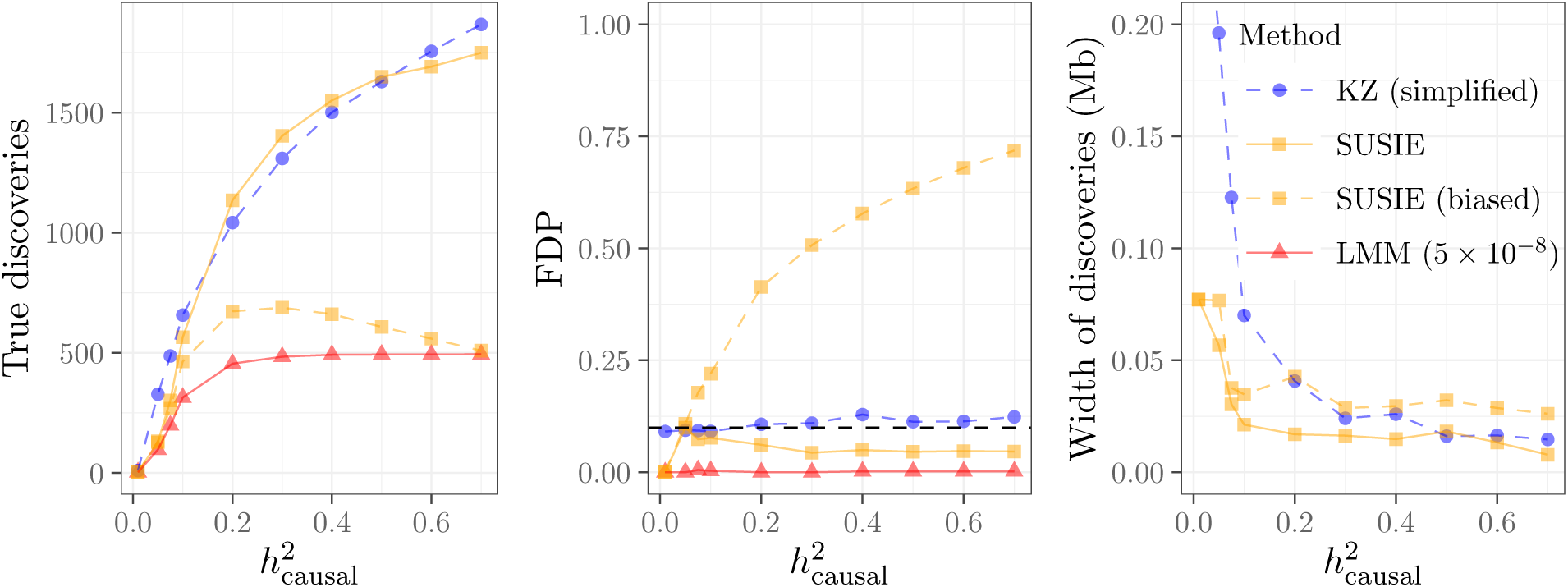
Fine-mapping with BOLT-LMM followed by SUSIE, with and without mitigating the selection bias. Other details as in Figure 4.

**FIG. S8.**
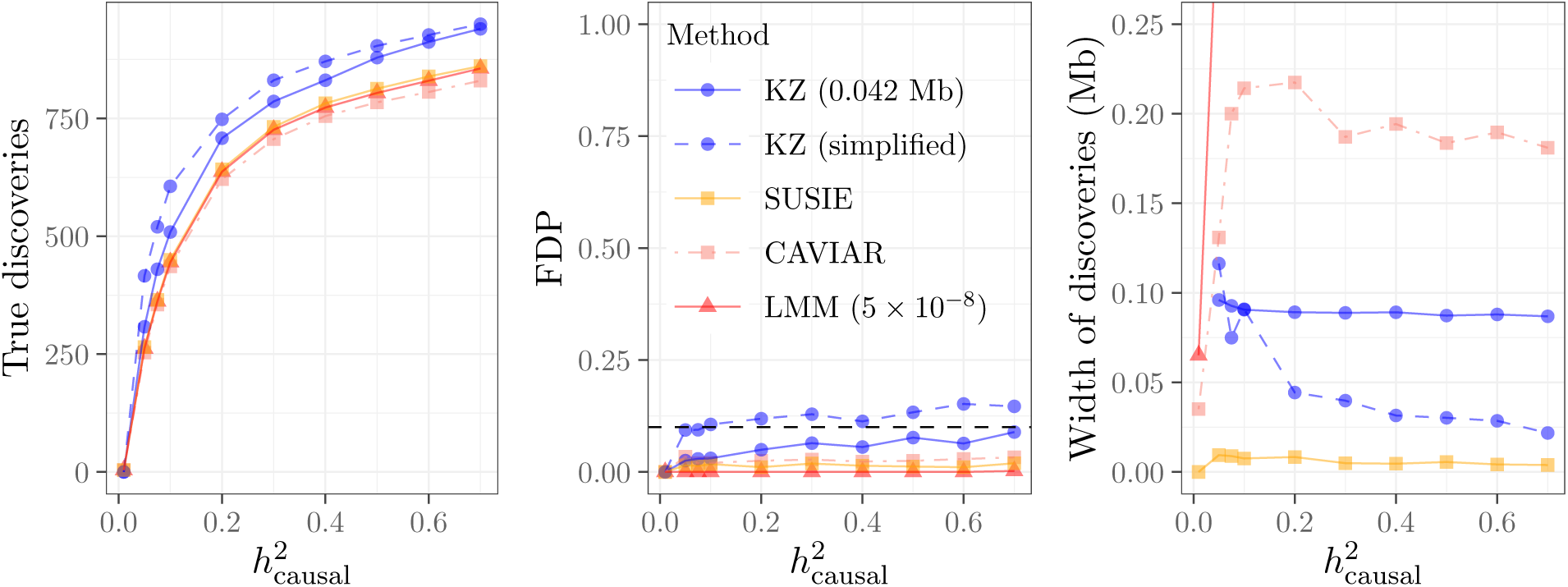
Fine-mapping performance of *KnockoffZoom* and BOLT-LMM followed by CAVIAR or SUSIE. Simulated trait as in Figure S6. Other details as in Figure 4.

##### 3. Contiguous groups and resolution

Until now, we have compared the fine-mapping resolution of different methods according to the average width spanned by each reported discovery, measured in base pairs. We believe that this is a particularly meaningful measure of localization in SNP-array data sets, where the true “causal” variants may not have been genotyped (Section II C), and discoveries should be interpreted as indicating a genetic region of interest (as precisely as possible) rather than exactly identifying the causal SNPs. However, it is also informative to look at different summary statistics of the discoveries to gain a broader perspective. For instance, we compare in Figure S9 the average size of the reported discoveries, measured in number of SNPs, or their homogeneity, measured in terms of the average *r*^2^ within each group (i.e., the average of the squared within-group correlation matrix over all entries), for the different methods in the same settings as in Figures 4 and S8.

These results show that *KnockoffZoom* typically reports discoveries containing more SNPs compared to its alternatives, which is not very surprising given that it tests pre-defined (and in this case contiguous) hypotheses, and therefore lacks some of the flexibility of other fine-mapping tools to explicitly discard nearby variants that are not significant. However, these numbers become more similar as the signal strength increases and *KnockoffZoom* increasingly localizes causal effects at the highest resolution. Regarding the homogeneity of the discoveries, the SNPs reported within the same group by SUSIE are the most similar to each other, with an average *r*^2^ approximately equal to 0.9. The homogeneity of the *KnockoffZoom* discoveries is typically lower compared to those of SUSIE but higher compared to those of CAVIAR. In any case, as the signal strength increases, *KnockoffZoom* increasingly reports single-SNP discoveries.

#### H. Impact of allele frequency

Figure S10 shows the high-resolution performance of *KnockoffZoom* stratified by minor allele frequency. These results show that our method performs well both for higher and lower-frequency variants, even though the knockoffs for lower-frequency variants are less accurate (Section S4 C). Furthermore, *KnockoffZoom* appears to be even more powerful for lower-frequency variants in these simulations, which is not very surprising given that there are stronger signals on rarer variants (i.e., the effect sizes are scaled by the inverse standard deviation of the allele count; see Section II D 1). Although the power will generally depend on the true effect sizes and allele frequencies, it is reassuring to observe that *KnockoffZoom* is not intrinsically limited by lower-frequency variants.

**FIG. S9.**
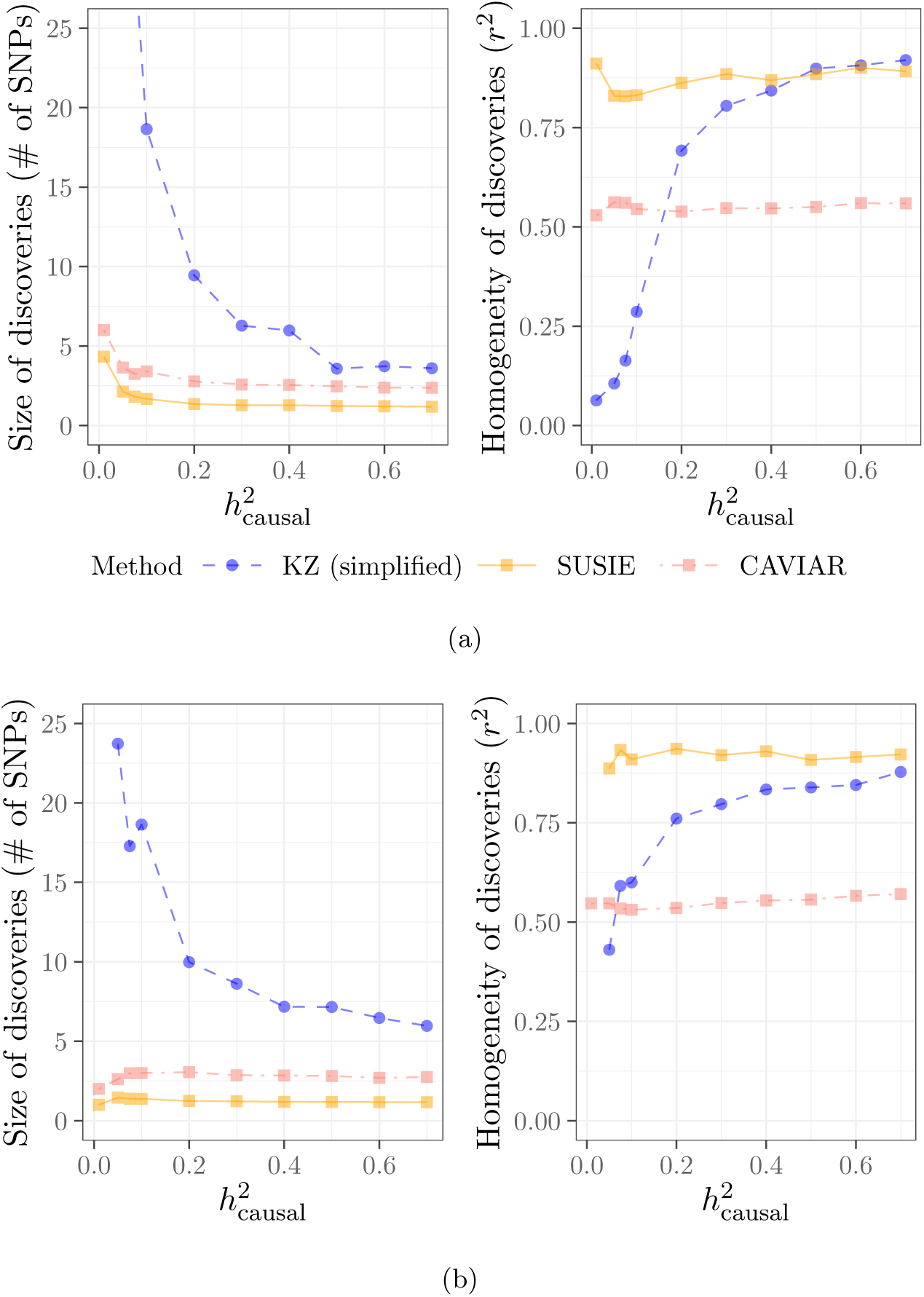
Resolution of fine-mapping discoveries in terms of average number of reported SNPs according to different measures, as a function of the heritability of the simulated trait. Left: average width measured in Mb (lower is better); center: average size measured in number of SNPs (lower is better); right: average homogeneity measured in mean pairwise *r*^2^ (higher is better). (a): other details as in Figure 4; (b): other details as in Figure S8.

**FIG. S10.**
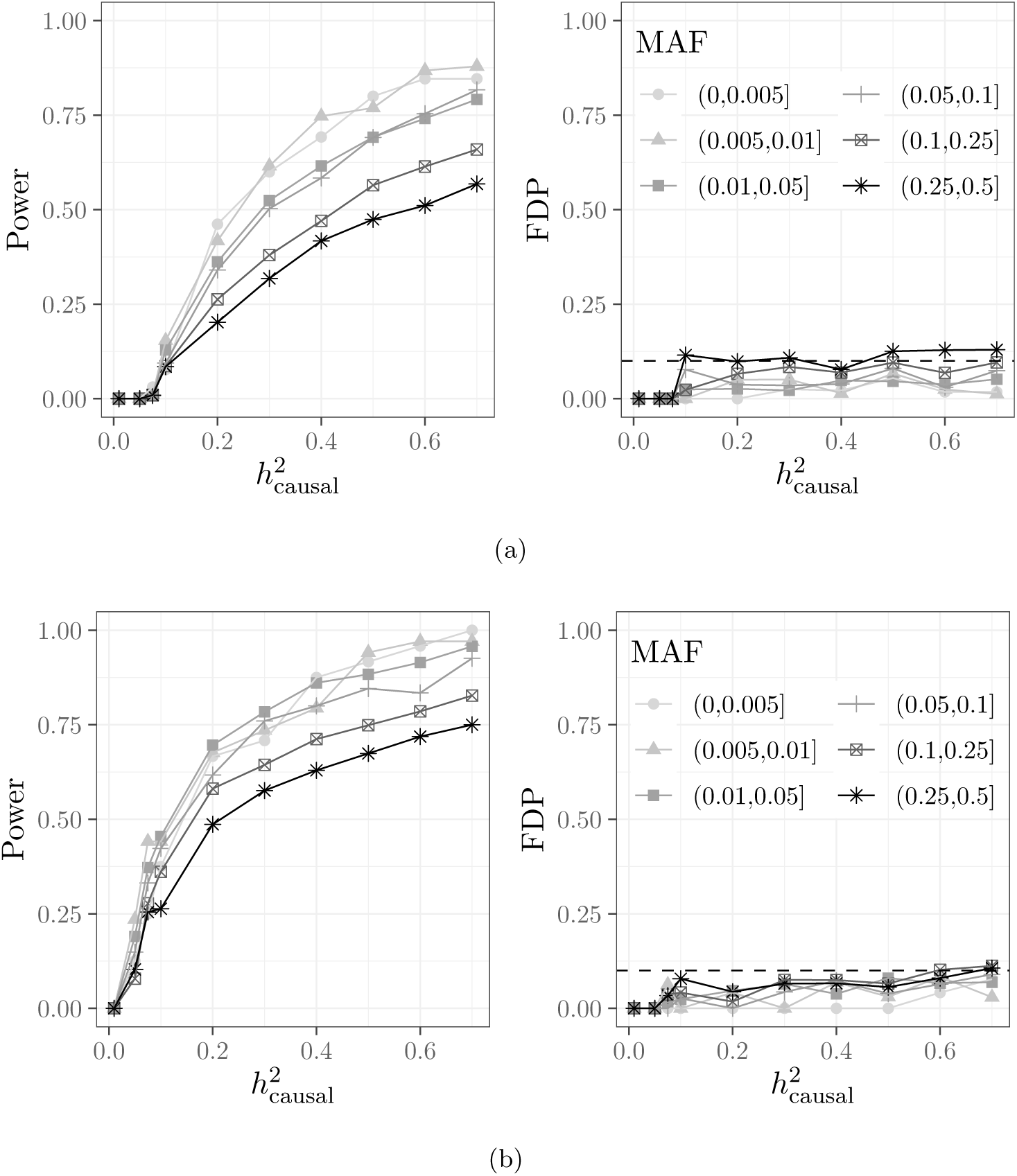
Fine-mapping performance of *KnockoffZoom* at the single-SNP resolution, stratified by the minor allele frequency of the causal variants. (a): other details as in Figure 4; (b): other details as in Figure S8.

#### I. Repeated simulations with smaller sample size

We investigate the average behaviour of *KnockoffZoom* and its alternatives on multiple independent realizations of the genotype data and the simulated phenotypes. We divide the available genotype observations into 10 disjoint subsets, each containing 30k individuals, and analyze them separately after generating artificial phenotypes for each. The FDR is estimated by averaging the FDP over the 10 repetitions. The results are in Figures S11 and S12.

**FIG. S11.**
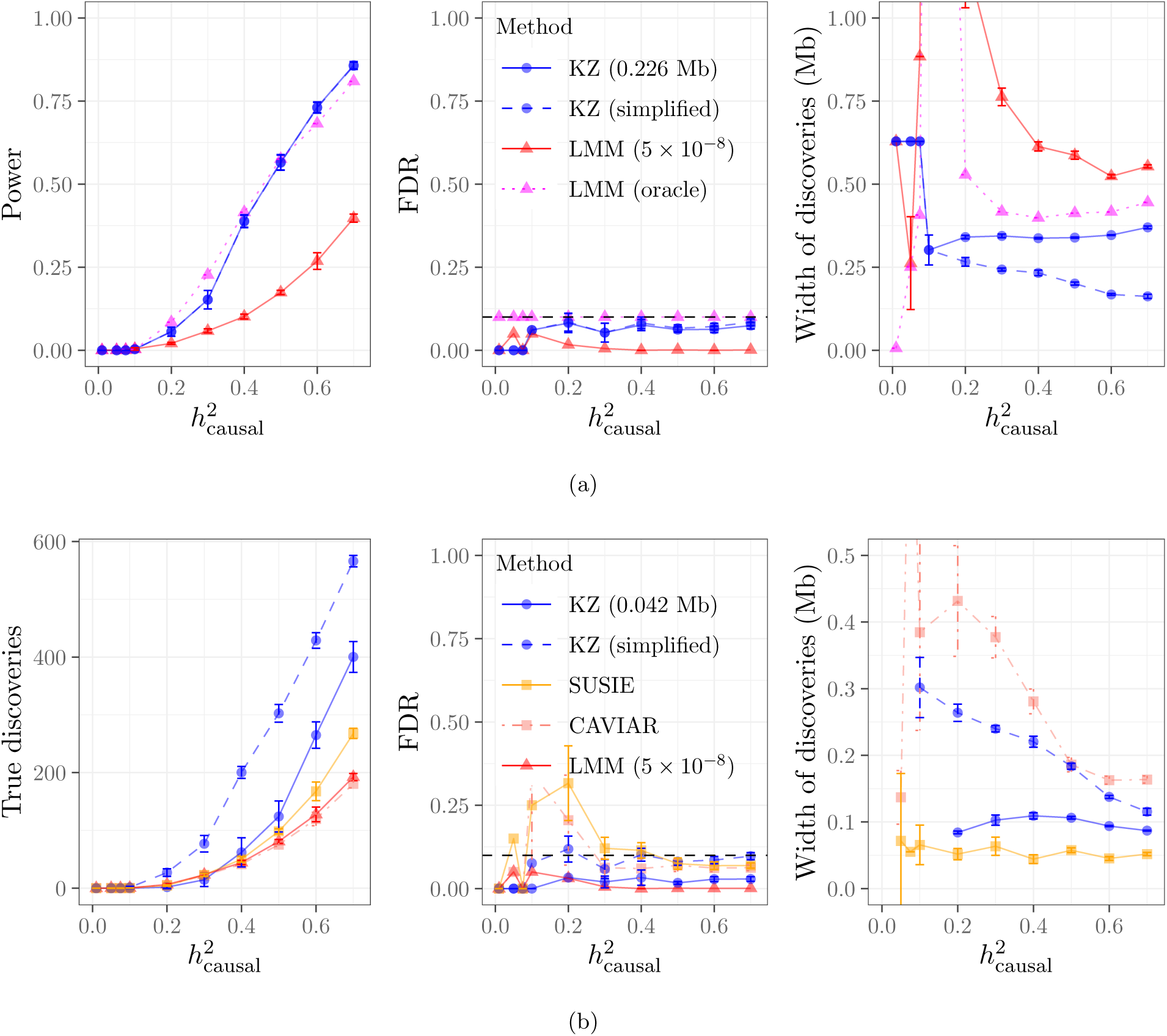
Performance of *KnockoffZoom* and BOLT-LMM for simulated phenotypes, repeating the experiments 10 times on disjoint subsets of the data, each including 30k individuals. The error bars indicate 95% confidence intervals for the mean quantities, estimated from 10 independent replications of the trait given the same genotypes. The other details in (a) and (b) are as in Figures 3 and 4, respectively.

**FIG. S12.**
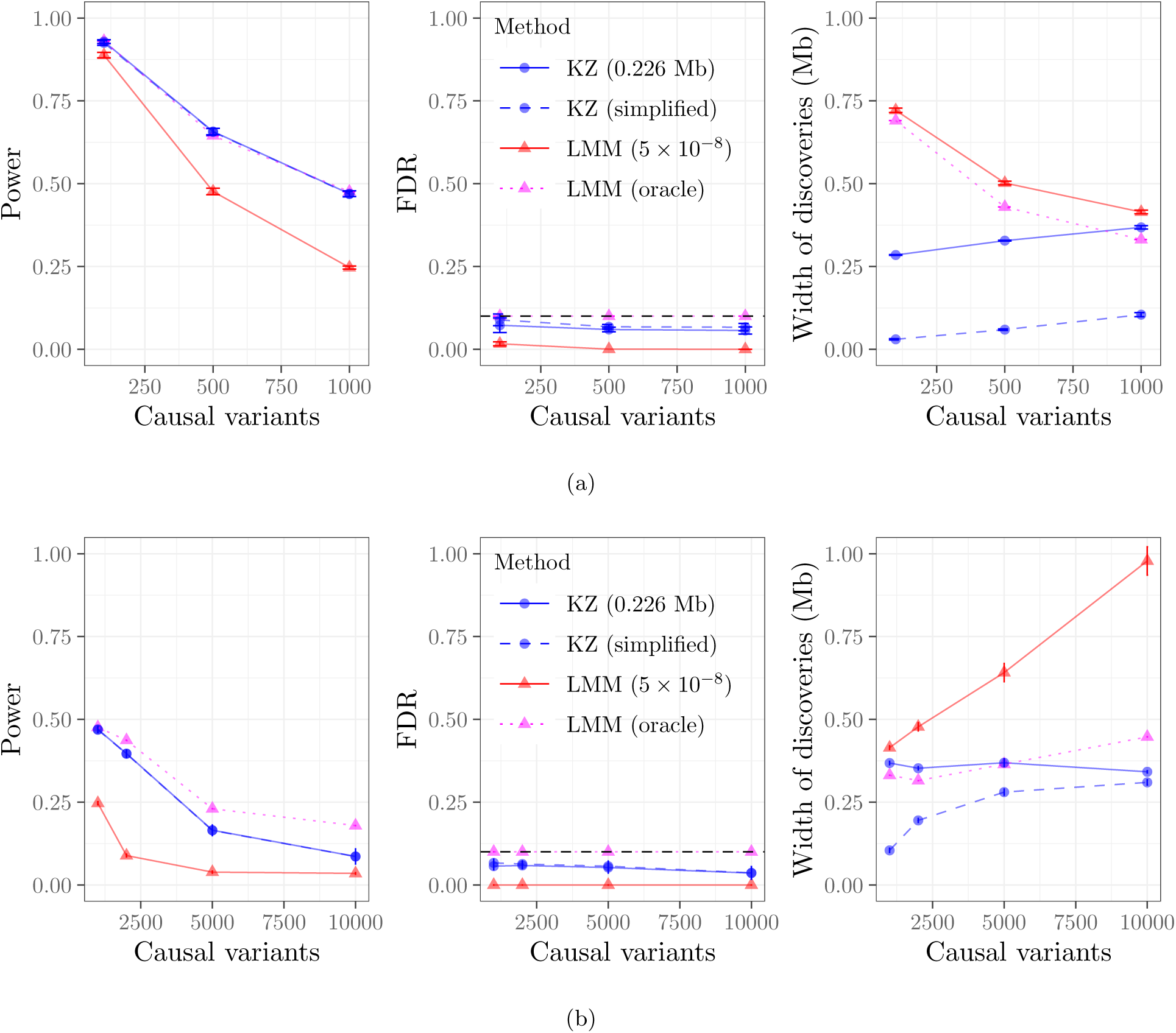
Performance of *KnockoffZoom* and BOLT-LMM for a simulated trait with different genetic architectures from Table S4, as a function of the number of causal variants. The heritability of the trait is 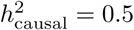. (a): architectures 3,4,5; (b): architectures 5,6,7,8. Other details as in Figure S11 (a).

#### J. Coordinating discoveries across resolutions

We analyze empirically the performance of *KnockoffZoom* with the consistent-layers knockoff filter (Algorithm 1). The results in Figure S13 indicate that the consistent-layers filter without the 1.93 correction factor performs similarly to the usual filter applied separately at each resolution, although it loses some power at high resolution. By contrast, the 1.93 factor is overly conservative, especially if there is a large number of causal variants with weak signals (Figure S13b).

**FIG. S13.**
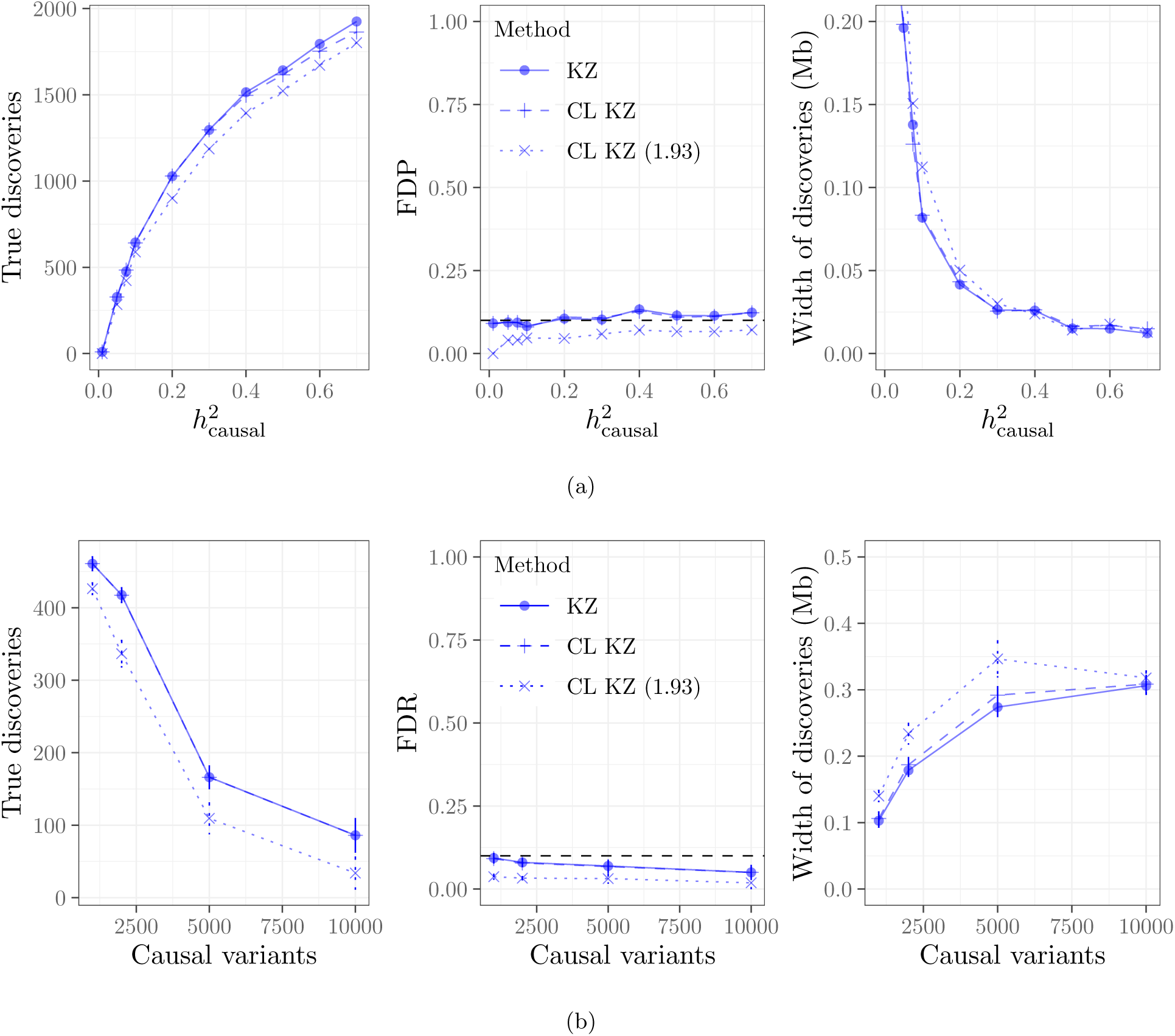
Performance of *KnockoffZoom* (KZ) with and without the consistent-layers knockoff filter. When enforcing layer consistency, we either include or not include the 1.93 factor discussed in Section S1 B; i.e., CL KZ (1.93), CL KZ. The curves corresponding to KZ and CL KZ are almost overlapping. The numbers of findings are simplified in all cases by counting only the finest discoveries in each locus, as in Figure 4. (a): Other details as in Figure 4. (b): Other details as in Figure S12.

#### K. Assessing the individual significance of each discovery

*KnockoffZoom* controls the FDR so that at most a pre-specified fraction *q* of the discoveries are false positives, on average. In addition to this global guarantee, we can quantify, to some extent, the statistical significance of the individual findings. We have not discussed this in the paper in the interest of space and because the underlying theory is not fully developed. Nonetheless, the main idea is intuitive. Keep in mind that the basic ingredient of the knockoff filter^26^ is a suitable estimate of the false discovery proportion: FDP(*t*), i.e., the fraction of false discoveries if we reject all hypotheses in {*g* : *W*_*g*_ ≥ *t*}. Note that the test statistics *W*_*g*_ are defined in Online Methods B. Additional information can be extracted from the test statistics by estimating a local version of the FDR^27^, e.g., the fraction of false discoveries in {*g* : *t* − Δ*t*_1_ ≤ *W*_*g*_ ≤ *t* + Δ*t*_2_}, for some choice of Δ*t*_1_ and Δ*t*_2_. For this purpose, we propose the following estimator:

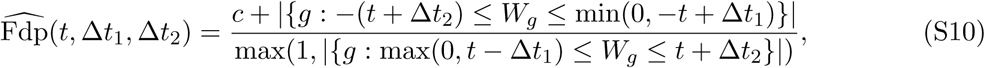

with either *c* = 0 (more liberal) or *c* = 1 (more conservative). The usual estimate of the FDP computed by the knockoff filter^26^ is recovered if Δ*t*_1_ = 0 and Δ*t*_2_ → ∞:

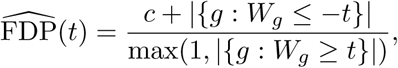

We expect that the findings whose test statistics fall in regions of low 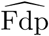 are less likely to be false positives. The choices of Δ*t*_1_ and Δ*t*_2_ determine the accuracy of our estimates: if the span is smaller, the local FDR encodes information at higher resolution; however, our estimate is noisier because it must rely on fewer statistics. We compute the estimate in (S10) in simulations (Figure S14) and on real data (Figure S16), setting Δ*t*_1_ = 100 = Δ*t*_2_ and *c* = 0 (by contrast, we use *c* = 1 in the knockoff filter). This approach is reasonable if *KnockoffZoom* reports sufficiently many discoveries; otherwise, the estimated local FDP may have very high variance.

The simulation in Figure S14 is carried out on an artificial phenotype with the first genetic architecture in Table S4 and heritability 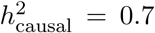. We plot the FDP as a function of the number of selected variants, based on the ordering defined by the *KnockoffZoom* test statistics.

We can draw two interesting observations from these results. First, the estimated cumulative FDP tracks the curve corresponding to the true FDP closely, which is consistent with the fact that the FDP of *KnockoffZoom* is almost always below the nominal FDR level throughout our simulations (Section II D), even though the theoretical guarantee refers to the average behavior. The low variance in the estimated FDP may be explained by the size of the dataset and the large number of discoveries. Second, the estimated local FDP also approximates the corresponding true quantity quite precisely, especially for the statistics above the rejection threshold. This may be very valuable in practice, because it provides us with a good educated guess of which discoveries are likely to be false positives. Whether we can rigorously state more precise results is an interesting question that we plan to explore deeper in the future.

**FIG. S14.**
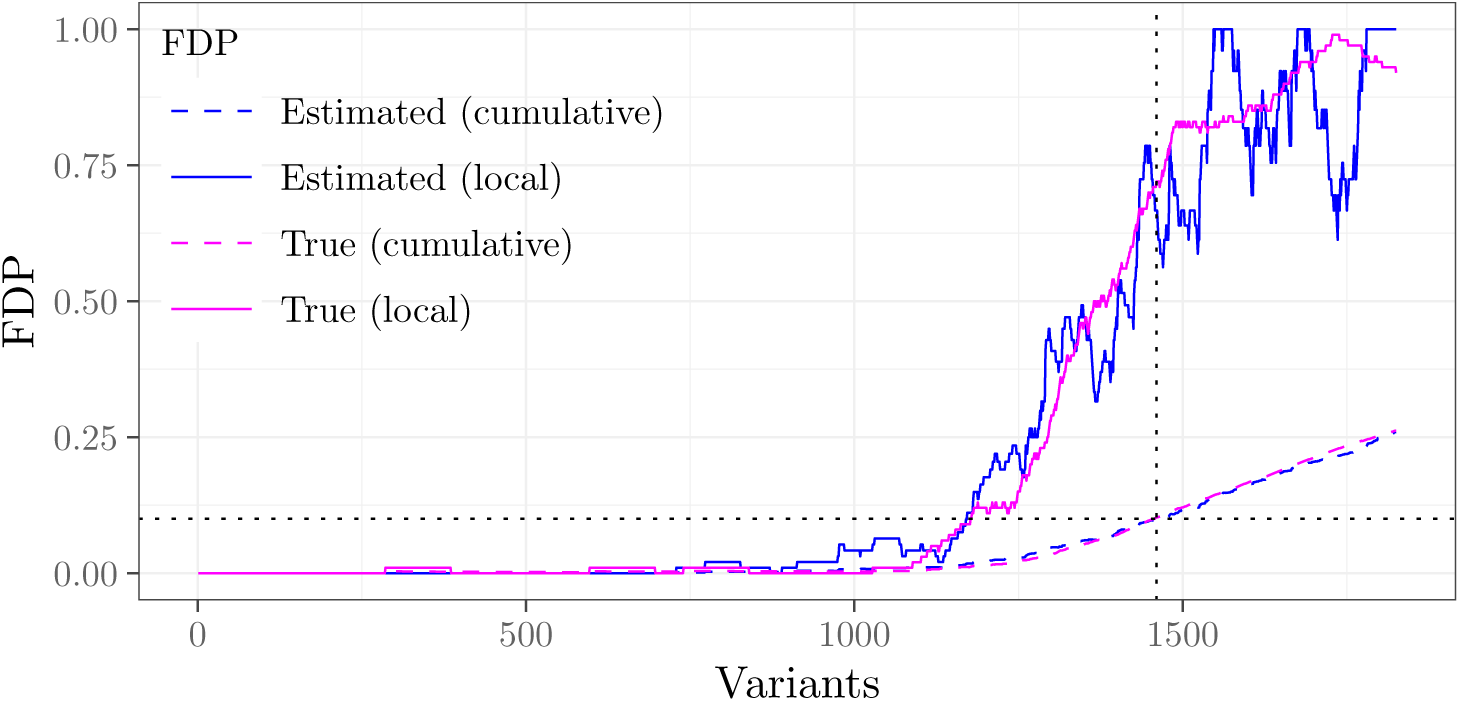
Estimated and true proportion of false discoveries obtained with *KnockoffZoom* at medium resolution (0.018 Mb), for a simulated trait with 2500 causal variants. The dotted vertical line indicates the adaptive significance threshold computed by the knockoff filter at the nominal FDR significance of 0.1. This corresponds exactly to the last crossing of the 0.1 level (dotted horizontal line) by the estimated cumulative FDP curve. The local (or estimated) FDP is computed by looking at 100 statistics within a rolling window.

#### L. Additional information about the implementation of alternative methods

In order to help others reproduce our simulations and build upon our work to design new numerical experiments in the future, we have shared the helper code used to run the comparisons discussed in this paper (https://github.com/msesia/ukbiobank_knockoffs), in addition to the code that implements our method. The version numbers and the input parameters of the relevant third-party software are also summarized below.

##### 1. BOLT-LMM

We apply BOLT-LMM (v. 2.3.2) using the default settings recommended in the official user manual (https://data.broadinstitute.org/alkesgroup/BOLT-LMM/). In the numerical experiments, we fit the model parameters on a subset of SNPs to reduce the computation cost (as suggested in the user manual for the analysis of large data sets), while we use all SNPs to fit the model parameters in the real data analysis.

##### 2. PLINK

We clump nearby significant variants identified by BOLT-LMM by applying PLINK (v. 1.90) using a 5 Mb window and an *r*^2^ threshold equal to 0.01 (to include sub-threshold SNPs in LD with an existing clump), while the secondary significance level is 10^−2^, as in earlier work.^28^

##### 3. CAVIAR

We perform fine-mapping using CAVIAR (v. 2.2) with confidence level equal to 0.9, setting the maximum number of causal SNPs equal to 2 to keep the computational cost manageable in large loci. The other parameters are equal to their default values.

##### 4. SUSIE

We perform fine-mapping using the SUSIE R package (v. 0.7.1) with nominal coverage 0.9, setting the scaled prior variance parameter equal to 0.1 (we observed that this performed better in our examples compared to the default value of 0.2) and allowing the estimation of the residual variance. The other parameters are equal to their default values.

### S5. DATA ANALYSIS

#### A. Phenotype definition in the UK Biobank

We have analyzed the phenotypes in the UK Biobank data defined in Table S5.

**TABLE S5.**
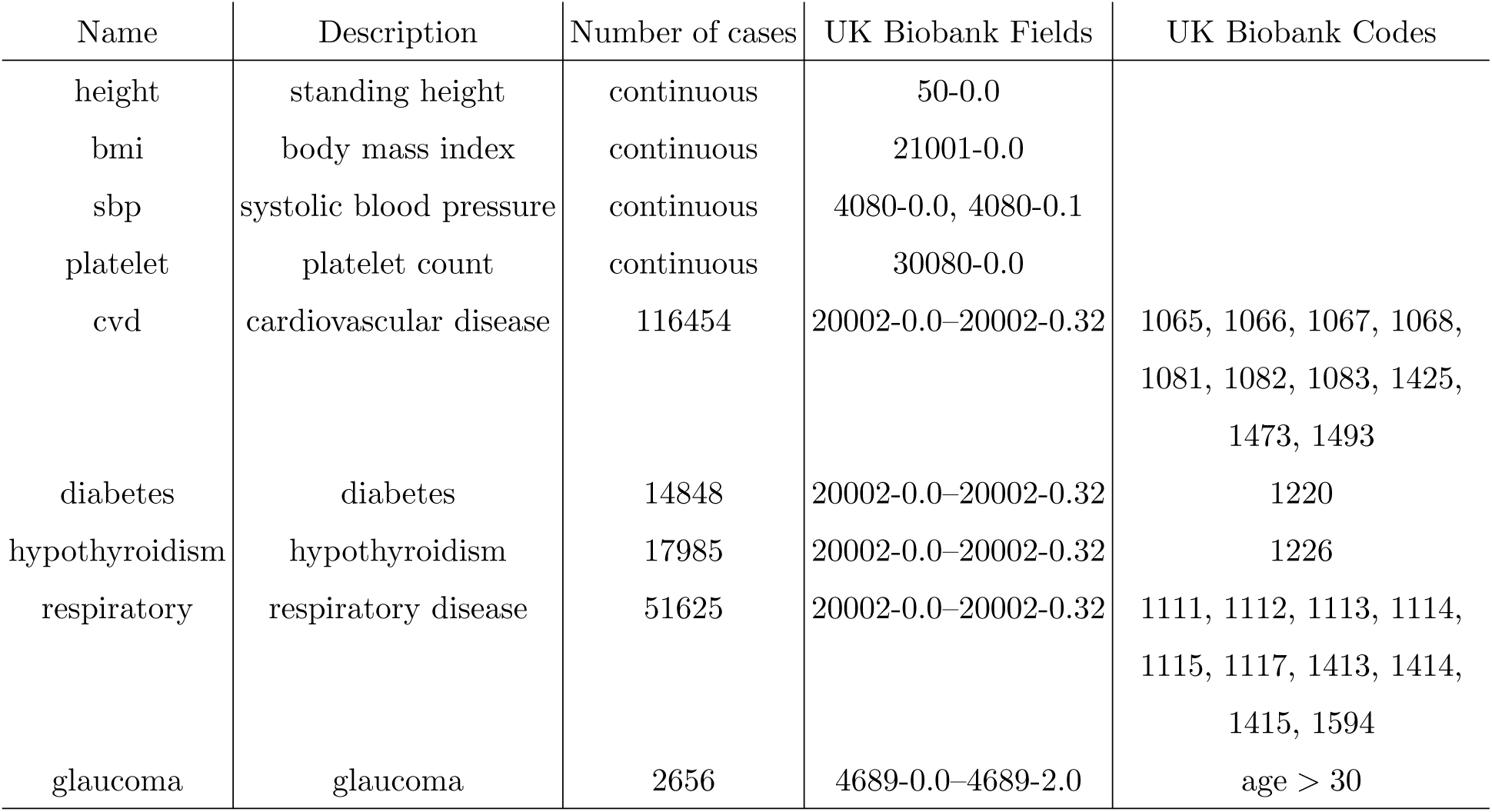
Phenotype definition in the UK Biobank. The numbers of disease cases refer to the subset of unrelated British individuals that passed our quality control.

| Name | Description | Number of cases | UK Biobank Fields | UK Biobank Codes |
| --- | --- | --- | --- | --- |
| height | standing height | continuous | 50-0.0 |  |
| bmi | body mass index | continuous | 21001-0.0 |  |
| sbp | systolic blood pressure | continuous | 4080-0.0, 4080-0.1 |  |
| platelet | platelet count | continuous | 30080-0.0 |  |
| cvd | cardiovascular disease | 116454 | 20002-0.0–20002-0.32 | 1065, 1066, 1067, 1068,<br>1081, 1082, 1083, 1425,<br>1473, 1493 |
| diabetes | diabetes | 14848 | 20002-0.0–20002-0.32 | 1220 |
| hypothyroidism | hypothyroidism | 17985 | 20002-0.0–20002-0.32 | 1226 |
| respiratory | respiratory disease | 51625 | 20002-0.0–20002-0.32 | 1111, 1112, 1113, 1114,<br>1115, 1117, 1413, 1414,<br>1415, 1594 |
| glaucoma | glaucoma | 2656 | 4689-0.0–4689-2.0 | age > 30 |

#### B. Number of discoveries

Table S6 shows that the inclusion of the principal components has little effect on our analysis, which confirms that we are not significantly affected by confounding due to population structure, at least within this cohort of unrelated British individuals.

**TABLE S6.** Number of distinct *KnockoffZoom* findings at different resolutions, with and without including principal components. The findings obtained with principal components correspond to those in Table I.

| Phenotype | Resolution |  |  |  |  |  |  |  |  |  |  |  |  |  |
| --- | --- | --- | --- | --- | --- | --- | --- | --- | --- | --- | --- | --- | --- | --- |
|  | 0.226 Mb |  | 0.088 Mb |  | 0.042 Mb |  | 0.018 Mb |  | 0.004 Mb |  | 0.001 Mb |  | single-SNP |  |
|  | PC | no<br>PC | PC | no<br>PC | PC | no<br>PC | PC | no<br>PC | PC | no<br>PC | PC | no<br>PC | PC | no<br>PC |
| height | 3284 | 3252 | 1976 | 2007 | 823 | 785 | 388 | 391 | 336 | 335 | 170 | 214 | 173 | 167 |
| bmi | 1804 | 1808 | 555 | 471 | 60 | 46 | 33 | 33 | 24 | 24 | 0 | 21 | 15 | 17 |
| platelet | 1460 | 1479 | 890 | 880 | 408 | 413 | 276 | 236 | 161 | 200 | 181 | 156 | 143 | 146 |
| sbp | 722 | 745 | 297 | 322 | 95 | 129 | 0 | 0 | 0 | 0 | 0 | 0 | 0 | 0 |
| cvd | 514 | 475 | 182 | 164 | 51 | 51 | 0 | 0 | 0 | 0 | 0 | 0 | 0 | 0 |
| hypothyroidism | 212 | 163 | 108 | 103 | 0 | 0 | 0 | 0 | 0 | 0 | 0 | 0 | 21 | 21 |
| respiratory | 176 | 183 | 65 | 60 | 41 | 12 | 13 | 13 | 14 | 14 | 12 | 12 | 0 | 0 |
| diabetes | 50 | 48 | 33 | 33 | 21 | 18 | 10 | 10 | 11 | 12 | 10 | 10 | 0 | 0 |
| glaucoma | 0 | 0 | 0 | 0 | 0 | 0 | 0 | 0 | 0 | 0 | 0 | 0 | 0 | 0 |

Table S7 quantifies the overlap between our low-resolution discoveries (including the principal components) and those obtained with BOLT-LMM applied to the same data. We say that findings made by different methods are overlapping if they are within 0.1 Mb of each other. These results confirm that our method makes many new findings, in addition to refining those reported by BOLT-LMM.

**TABLE S7.** Overlap of the discoveries made with *KnockoffZoom* at low resolution (0.226 Mb) and BOLT-LMM (clumped without consolidation), using the same data. *KnockoffZoom* is applied as in Table I. For example, our method reports 3284 findings for *height*, 2246 of which overlap at least one of the 1685 clumps found by the LMM, while 1038 are distinct from those found by the LMM. Conversely, only 6 out of 1685 discoveries made by BOLT-LMM are not detected by *KnockoffZoom*.

| Phenotype | Method | Discoveries | Overlapping | Distinct |
| --- | --- | --- | --- | --- |
| height | *KnockoffZoom* | 3284 | 2246 | 1038 |
|  | BOLT-LMM | 1685 | 1679 | 6 |
| bmi | *KnockoffZoom* | 1804 | 627 | 1177 |
|  | BOLT-LMM | 389 | 378 | 11 |
| platelet | *KnockoffZoom* | 1460 | 848 | 612 |
|  | BOLT-LMM | 723 | 709 | 14 |
| sbp | *KnockoffZoom* | 722 | 243 | 479 |
|  | BOLT-LMM | 197 | 188 | 9 |
| cvd | *KnockoffZoom* | 514 | 188 | 326 |
|  | BOLT-LMM | 156 | 144 | 12 |
| hypothyroidism | *KnockoffZoom* | 212 | 96 | 116 |
|  | BOLT-LMM | 96 | 91 | 5 |
| respiratory | *KnockoffZoom* | 176 | 59 | 117 |
|  | BOLT-LMM | 63 | 59 | 4 |
| diabetes | *KnockoffZoom* | 50 | 40 | 10 |
|  | BOLT-LMM | 47 | 43 | 4 |
| glaucoma | *KnockoffZoom* | 0 | 0 | 0 |
|  | BOLT-LMM | 5 | 0 | 5 |

#### C. Comparison with BOLT-LMM on a larger sample

Our discoveries using data from 350k unrelated British individuals are compared in Table S8 to those obtained in earlier work by applying BOLT-LMM to all 459,327 European subjects in the UK Biobank.^28^ We say that findings made by different methods are overlapping if they are within 0.1 Mb of each other. The LMM makes fewer findings despite the larger sample size. This indicates that our method is more powerful, in addition to providing more interpretable results.

**TABLE S8.**
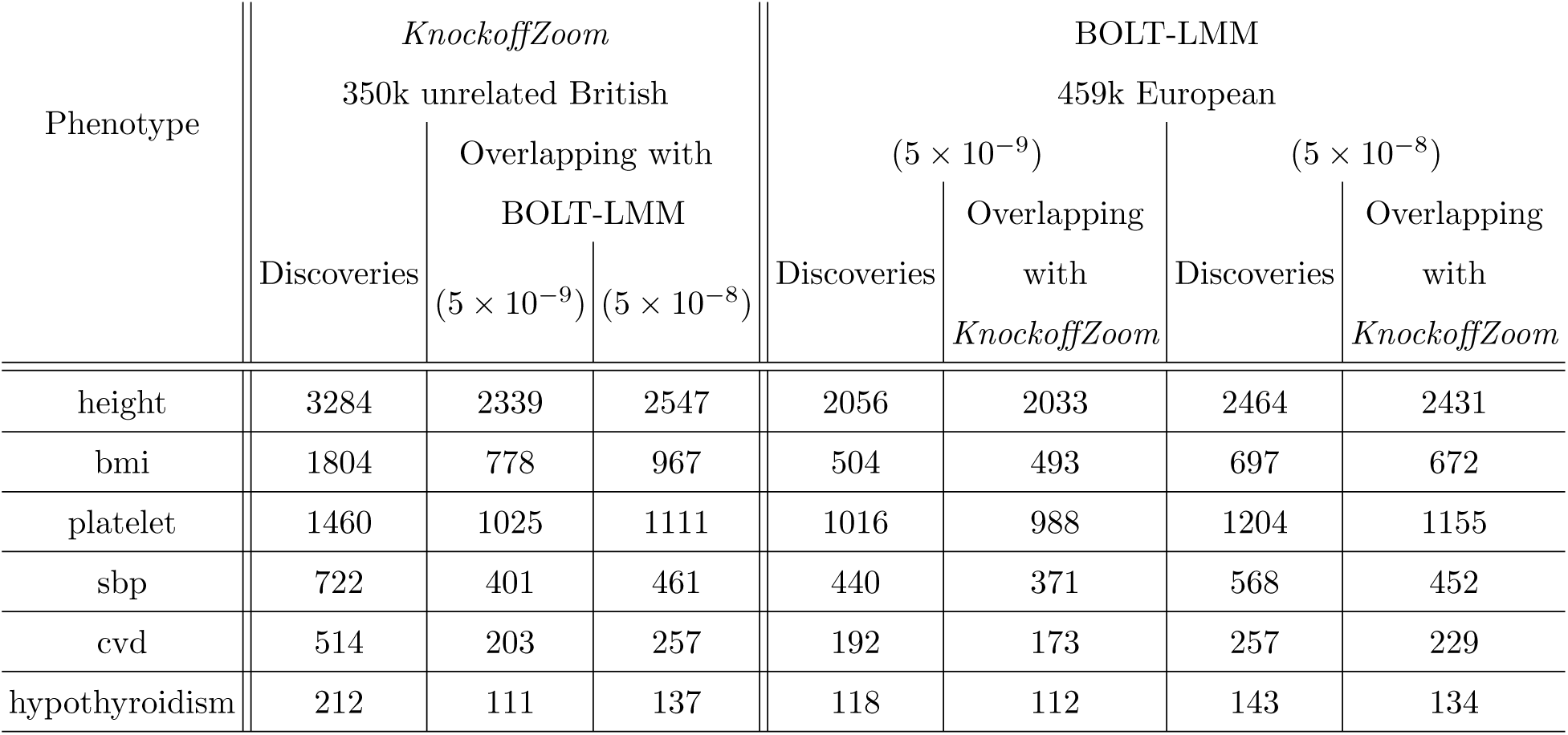
Comparison of the low-resolution (0.226 Mb) discoveries reported by *KnockoffZoom*, as in Table I, and those obtained by BOLT-LMM with a larger dataset. For example, BOLT-LMM reports 2056 discoveries for *height* at the significance level 5 × 10^−9^, 2033 of which overlap with at least one of our 3284 findings, while 2339 of our findings overlap with at least one discovery made by BOLT-LMM.

#### D. Coordinating discoveries across resolutions

Table S9 summarizes the results obtained by coordinating the discoveries at different resolutions.

**TABLE S9.**
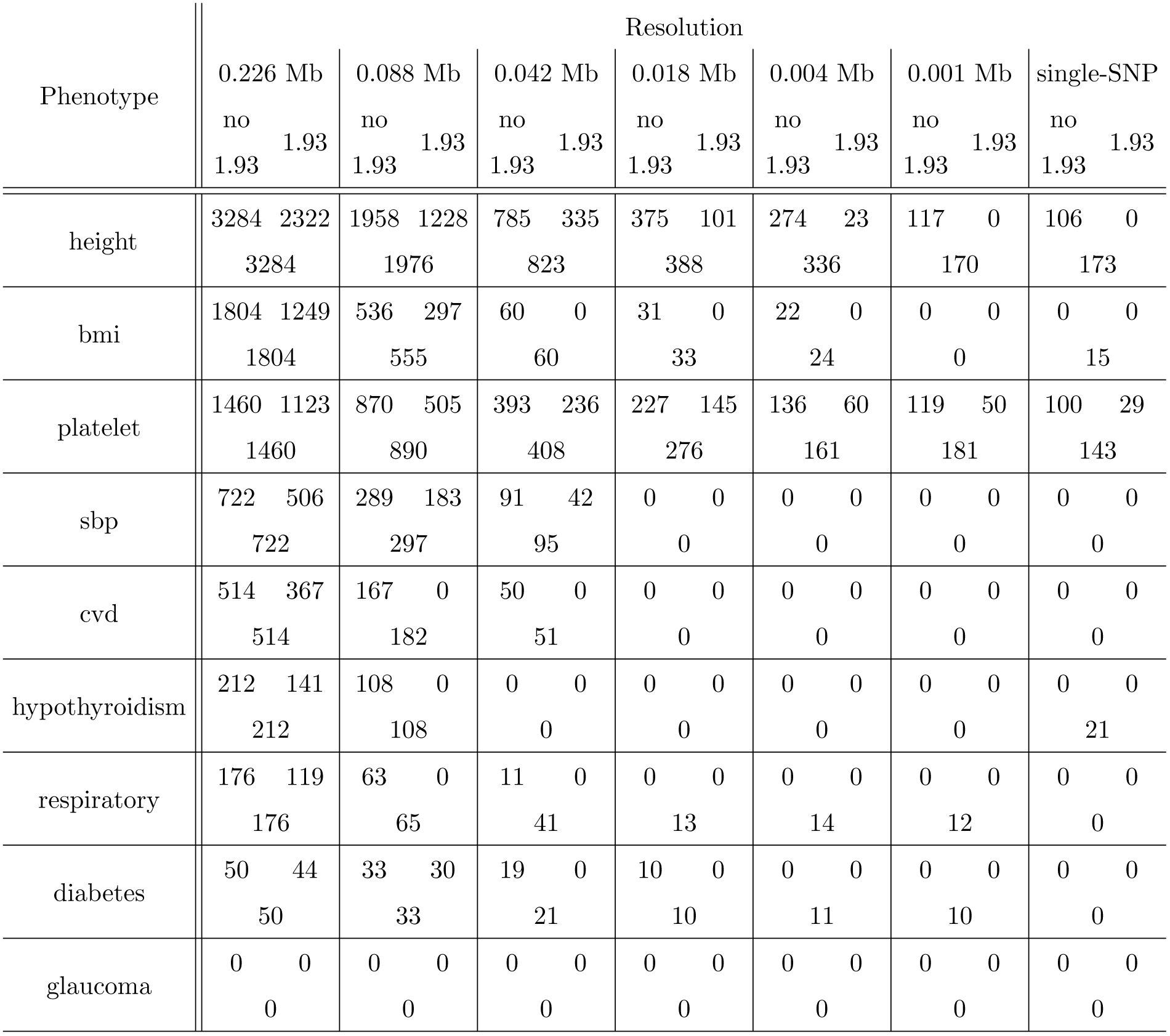
Distinct findings made by *KnockoffZoom* using the consistent-layers knockoff filter, without or with the 1.93 correction factor. The third number in each cell (on the new line) indicates the number of discoveries obtained separately at each resolution, without applying the consistent-layers knockoff filter.

Note that the consistent-layers knockoff filter is equivalent to the usual knockoff filter at the lowest resolution if the 1.93 correction factor is omitted. Therefore, the values in the first column of Table S9 are identical to the values in the first column of Table S6. At higher resolutions, the consistent-layers knockoff filter leads to slightly fewer findings, as also observed earlier in the simulations in Section S4 J. Including the theoretical 1.93 correction factor makes the consistent-layers knockoff filter significantly more conservative, as in the simulations in Section S4 J.

#### E. Reproducibility (low resolution)

**TABLE S10.** Overlap of the discoveries made with *KnockoffZoom* (low-resolution) and BOLT-LMM (clumped without consolidation). Other details as in Table II. For example, we make 121 discoveries for *height*, 56 of which are within 0.1 Mb of at least one of the 54 clumps found by the LMM, while 65 are distinct. In this case, only 5 out of 54 discoveries made by BOLT-LMM are not detected by *KnockoffZoom*.

| Phenotype | Method | # Discoveries |  |  |
| --- | --- | --- | --- | --- |
|  |  | Total | Overlapping | Distinct |
| height | *KnockoffZoom* | 121 | 56 | 65 |
|  | BOLT-LMM | 54 | 49 | 5 |
| platelet | *KnockoffZoom* | 81 | 44 | 37 |
|  | BOLT-LMM | 47 | 42 | 5 |

**TABLE S11.** Estimated low-resolution power of *KnockoffZoom* and BOLT-LMM. Other details as in Table S10. We say that a finding reported by BOLT-LMM on the larger dataset is anticipated by those in the smaller dataset if it is within 0.1 Mb of at least one of them. For example, among the 2056 unconsolidated discoveries reported by BOLT-LMM for *height* on the large dataset, 398 are anticipated by the findings of *KnockoffZoom* on the smaller dataset, while only 280 are anticipated by BOLT-LMM using the same data.

| Phenotype | Method | 30k unrelated British |  |  | 459k European (BOLT-LMM) |  |  |  |
| --- | --- | --- | --- | --- | --- | --- | --- | --- |
|  |  | Discoveries |  |  | Discoveries |  | Discoveries (consolidated) |  |
|  |  | # | Not replicated | Size (Mb) | Anticipated | Size (Mb) | Anticipated | Size (Mb) |
| height | *KnockoffZoom* | 121 | 8 (6.6%) | 0.308 | 398/2056 (19.4%) | 0.789 | 92/807 (11.4%) | 1.162 |
|  | LMM | 54 | 0 (0.0%) | 0.965 | 280/2056 (13.6%) | 0.789 | 48/807 (5.9%) | 1.162 |
|  | LMM (BH) | 714 | 203 (28.4%) | 0.379 | 1047/2056 (50.9%) | 0.789 | 325/807 (40.3%) | 1.162 |
| platelet | *KnockoffZoom* | 81 | 5 (6.2%) | 0.319 | 235/1016 (23.1%) | 0.805 | 65/525 (12.4%) | 0.953 |
|  | LMM | 47 | 0 (0.0%) | 0.674 | 219/1016 (21.6%) | 0.805 | 42/525 (8.0%) | 0.953 |
|  | LMM (BH) | 272 | 92 (33.8%) | 0.433 | 422/1016 (41.5%) | 0.805 | 136/525 (25.9%) | 0.953 |

#### F. Reproducibility (high resolution)

**TABLE S12.** Gene ontology enrichment analysis using GREAT^29^ for the *KnockoffZoom* results on *platelet*. The uncorrected p-values for 5 relevant biological processes are shown at each resolution.

| Resolution | Platelet activation | Hemostasis | Blood coagulation | Wound healing | Response to wounding |
| --- | --- | --- | --- | --- | --- |
| 0.226 | $1.8 \times 10^{-08}$ | $2.8 \times 10^{-12}$ | $3.9 \times 10^{-12}$ | $2.0 \times 10^{-11}$ | $5.6 \times 10^{-13}$ |
| 0.088 | $2.8 \times 10^{-10}$ | $3.1 \times 10^{-20}$ | $6.1 \times 10^{-20}$ | $1.2 \times 10^{-19}$ | $4.8 \times 10^{-21}$ |
| 0.042 | $1.0 \times 10^{-08}$ | $1.1 \times 10^{-20}$ | $3.6 \times 10^{-20}$ | $3.6 \times 10^{-18}$ | $6.8 \times 10^{-17}$ |
| 0.018 | $5.4 \times 10^{-08}$ | $3.0 \times 10^{-21}$ | $1.3 \times 10^{-20}$ | $4.0 \times 10^{-18}$ | $3.9 \times 10^{-17}$ |
| 0.004 | $2.8 \times 10^{-09}$ | $5.4 \times 10^{-23}$ | $3.4 \times 10^{-22}$ | $1.8 \times 10^{-19}$ | $4.7 \times 10^{-17}$ |
| 0.001 | $3.5 \times 10^{-09}$ | $3.0 \times 10^{-23}$ | $1.8 \times 10^{-22}$ | $3.1 \times 10^{-20}$ | $3.5 \times 10^{-19}$ |
| single-SNP | $2.6 \times 10^{-09}$ | $1.8 \times 10^{-24}$ | $1.4 \times 10^{-23}$ | $5.4 \times 10^{-21}$ | $1.8 \times 10^{-17}$ |

**TABLE S13.** Most serious previously known consequence (dbSNP^30^) of each of the 143 variants discovered by *KnockoffZoom* at the highest resolution for the phenotype *platelet* in the UK Biobank.

| Consequence | Discoveries |
| --- | --- |
| nonsense | 1 |
| missense | 26 |
| synonymous | 2 |
| 3-prime UTR | 7 |
| splice donor | 1 |
| intronic | 74 |
| 500B downstream | 2 |
| 2KB upstream | 4 |
| intergenic | 26 |

**TABLE S14.** Most relevant previously reported associations for some of the 143 variants discovered by *KnockoffZoom* at the highest resolution for the phenotype *platelet* in the UK Biobank.

| Previous association | Discoveries |
| --- | --- |
| platelet | 61 |
| hemoglobin | 4 |
| red cells | 2 |
| squamous cell carcinoma | 2 |
| white cells | 2 |
| breast cancer | 1 |
| coronary artery disease | 1 |
| fatty liver disease | 1 |
| macular degeneration | 1 |
| menarche (age of onset) | 1 |
| obesity | 1 |
| none | 66 |

**TABLE S15.** Missense variants localized by *KnockoffZoom* at the highest resolution, for the phenotype *platelet* in the UK Biobank, that have not been reported before.

| Chromosome | SNP | Position | Gene | Consequence |
| --- | --- | --- | --- | --- |
| 19 | Affx-15656246 | 19765499 | ATP13A1 | missense |
| 19 | rs12983010 | 39229089 | CAPN12 | missense |
| 1 | rs140584594 | 110232983 | GSTM1 | missense |
| 17 | rs79007502 | 33880305 | SLFN14 | missense |
| 2 | rs76774368 | 160604514 | MARCH7 | missense |
| 3 | rs34095724 | 184099050 | CHRD | missense |

#### G. Distribution of minor allele frequencies

The distribution of minor allele frequencies for the genotyped variants in the UK Biobank is summarized in Table S16. This table also shows the distribution of minor allele frequencies for the lead variants in the discoveries reported by *KnockoffZoom*. In this setting, the lead variant in each selected group is defined as that having the largest estimated regression coefficient in absolute value. For simplicity, we focus on the discoveries obtained. These results show that lower-frequency variants are less likely to be selected by *KnockoffZoom*. This observation is consistent with the fact that effects on rarer variants are intrinsically harder to detect.

**TABLE S16.** Minor allele frequency distribution of variants selected by *KnockoffZoom* at low resolution (0.226 Mb) for different traits, compared to the distribution of all genotyped variants in the UK Biobank.

|  | Min. | 1st Qu. | Median | Mean | 3rd Qu. | Max. |
| --- | --- | --- | --- | --- | --- | --- |
| All variants | 0.001 | 0.025 | 0.066 | 0.133 | 0.213 | 0.500 |
| Discoveries |  |  |  |  |  |  |
| height | 0.001 | 0.091 | 0.224 | 0.230 | 0.360 | 0.500 |
| bmi | 0.001 | 0.112 | 0.244 | 0.241 | 0.364 | 0.500 |
| platelet | 0.001 | 0.091 | 0.225 | 0.230 | 0.355 | 0.500 |
| sbp | 0.001 | 0.098 | 0.238 | 0.239 | 0.371 | 0.500 |
| cvd | 0.002 | 0.114 | 0.248 | 0.246 | 0.368 | 0.496 |
| hypothyroidism | 0.001 | 0.121 | 0.246 | 0.247 | 0.373 | 0.497 |
| respiratory | 0.001 | 0.082 | 0.234 | 0.228 | 0.354 | 0.496 |
| diabetes | 0.007 | 0.128 | 0.279 | 0.259 | 0.382 | 0.494 |

#### H. Chicago plots

Figure 6 complements the example in Figures 1 and 5 by including an estimate of the local FDP computed as in Figure S16, for each level of resolution. Moreover, we also include the results obtained with BOLT-LMM for comparison.

**FIG. S15.**
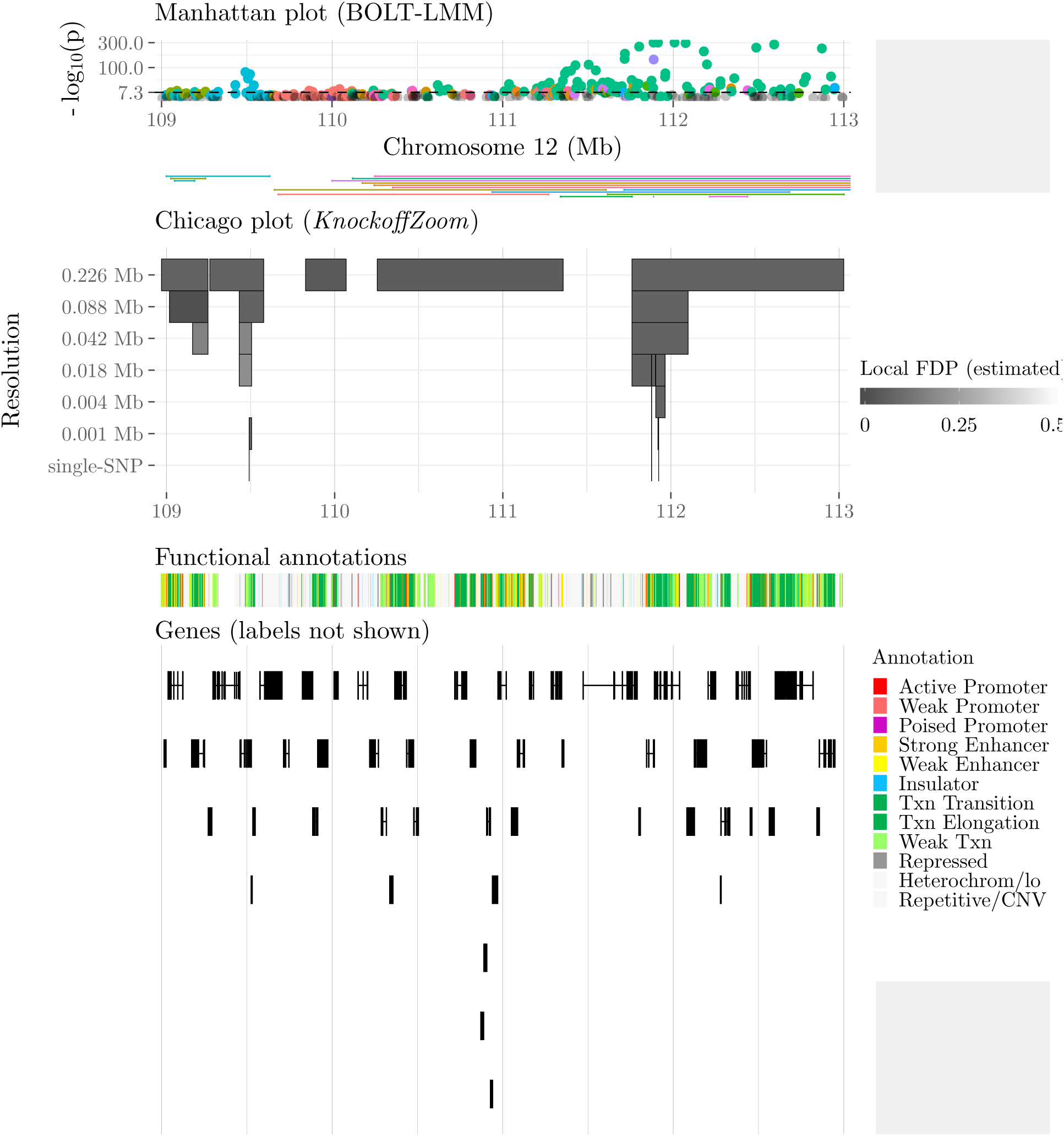
Visualization of some discoveries made with *KnockoffZoom* and BOLT-LMM for *platelet* in the UK Biobank. Other details as in Figures 1 and 5. This Chicago plot is upside-down.

#### I. Assessing the individual significance of each discovery

We assess the individual significance of our discoveries on real data as discussed in Section S4 K. Figure S16 shows an example of the estimated local and global FDP as a function of the number of selected variants, based on the ordering defined by the *KnockoffZoom* test statistics. This information is included in the visualization of our discoveries in Section S5 H.

**FIG. S16.**
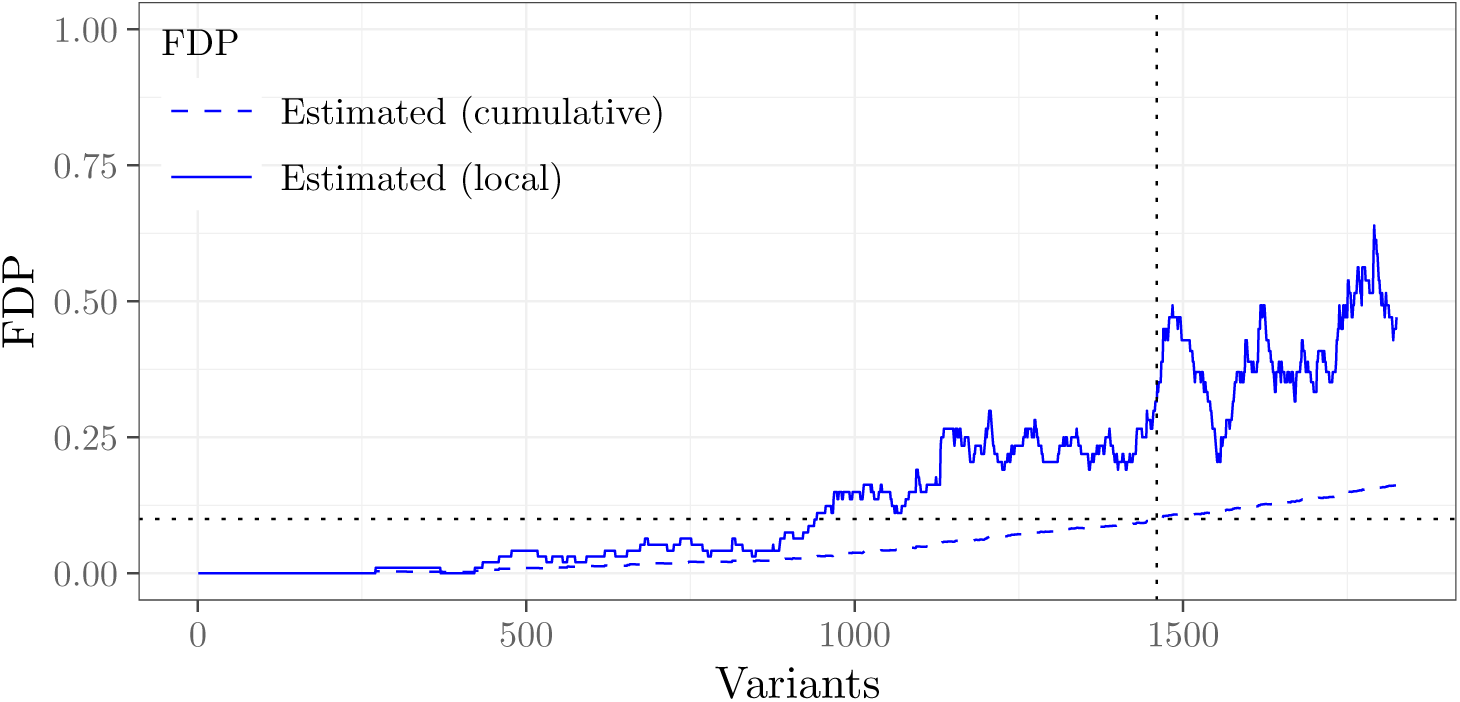
Estimated proportion of false discoveries obtained with *KnockoffZoom* at low-resolution (0.226 Mb) for the phenotype *platelet* in the UK Biobank. Other details as in Figure S14.

#### J. Variability upon knockoff resampling

We repeat the analysis of the UK Biobank phenotypes starting from the generation of knockoffs, setting a different seed for the random number generator. The numbers of discoveries obtained with these two independent sets of knockoffs are compared in Table S17, along with their overlap. Only the analyses for which *KnockoffZoom* reports at least 100 discoveries with the first random seed are shown here—more details on this choice are discussed below.

Table S17 shows that the *KnockoffZoom* results are quite stable (approximately 80% of the discoveries obtained with two different random seeds match), especially when the number of discoveries is large. In fact, most of the variability affects signals that are close to the FDR significance level and hence most difficult to detect (perhaps because they are tested at very high resolution, or because they have weak signals, or because they are false positives). We can see this explicitly in Tables S18–S20, where the discoveries are stratified by their individual significance measured in terms of the estimated local FDP; see also Sections S4 K and S5 I. These results indicate that discoveries with lower local FDP (those about which we are more confident) are much more stable. The same trend can also be seen more explicitly in Table S21, where we stratify the discoveries based on different levels of estimated local FDP.

In this section, we focus on phenotypes and resolutions with at least 100 discoveries because we cannot otherwise reliably estimate the local FDP. For the same reason, the relative variability of the *KnockoffZoom* discoveries may otherwise be higher. In any case, the guarantee that the FDR is controlled for any realization of the knockoffs holds regardless of the number of discoveries.

Finally, since we have already observed that a single wide locus reported by BOLT-LMM may contain multiple distinct *KnockoffZoom* discoveries, even when the latter is applied at low resolution, it is interesting to count how many of the loci reported by BOLT-LMM are identified by at least one *KnockoffZoom* discovery consistently upon knockoff resampling. The results summarized in Table S22 show that the vast majority (between 80% and 99%) of the significant loci identified by BOLT-LMM (in a larger sample) are also consistently reported by *KnockoffZoom*. Instead, the variability in *KnockoffZoom* typically involves either completely new loci (near the FDR threshold) or finer-grained details (multiple distinct discoveries within the same locus) that cannot be detected by BOLT-LMM.

**TABLE S17.** Numbers of *KnockoffZoom* discoveries obtained with different random seeds for the generation of knockoffs. The first and second values in each column refer to the results obtained without and with explicit coordination of results at different resolutions using the consistent-layers knockoff filter (these are equivalent at the lowest resolution). The stability percentage is defined as the average proportion of *KnockoffZoom* discoveries that are consistently reported with both random seeds.

| Resolution | Phenotype | Seed 1 |  | Seed 2 |  | Both seeds |  | Stable (%) |  |
| --- | --- | --- | --- | --- | --- | --- | --- | --- | --- |
|  |  | no CL | CL | no CL | CL | no CL | CL | no CL | CL |
| 0.226 Mb | height | 3284 |  | 3155 |  | 2792 |  | 86.8 |  |
|  | bmi | 1804 |  | 1841 |  | 1508 |  | 82.8 |  |
|  | platelet | 1460 |  | 1424 |  | 1196 |  | 83.0 |  |
|  | sbp | 722 |  | 802 |  | 616 |  | 81.1 |  |
|  | cvd | 514 |  | 472 |  | 379 |  | 77.0 |  |
|  | hypothyroidism | 212 |  | 186 |  | 167 |  | 84.3 |  |
|  | respiratory | 176 |  | 158 |  | 131 |  | 78.7 |  |
| 0.088 Mb | height | 1976 | 1958 | 1803 | 1791 | 1577 | 1562 | 83.6 | 83.5 |
|  | bmi | 555 | 536 | 670 | 646 | 447 | 426 | 73.6 | 72.7 |
|  | platelet | 891 | 870 | 975 | 914 | 755 | 729 | 81.1 | 81.8 |
|  | sbp | 297 | 289 | 360 | 323 | 234 | 223 | 71.9 | 73.1 |
|  | cvd | 182 | 167 | 227 | 215 | 147 | 131 | 72.8 | 69.7 |
|  | hypothyroidism | 108 | 108 | 77 | 76 | 67 | 67 | 74.5 | 75.1 |
| 0.042 Mb | height | 823 | 785 | 841 | 810 | 655 | 625 | 78.7 | 78.4 |
|  | platelet | 408 | 393 | 466 | 442 | 326 | 310 | 74.9 | 74.5 |
| 0.018 Mb | height | 388 | 375 | 331 | 304 | 265 | 244 | 74.2 | 72.7 |
|  | platelet | 276 | 227 | 231 | 213 | 185 | 164 | 73.6 | 74.6 |
| 0.004 Mb | height | 336 | 274 | 202 | 145 | 175 | 127 | 69.4 | 67.0 |
|  | platelet | 161 | 136 | 226 | 180 | 139 | 112 | 73.9 | 72.3 |
| 0.001 Mb | height | 170 | 117 | 121 | 103 | 100 | 78 | 70.7 | 71.2 |
|  | platelet | 181 | 119 | 202 | 154 | 148 | 97 | 77.5 | 72.2 |
| single-SNP | height | 173 | 106 | 134 | 92 | 115 | 72 | 76.1 | 73.1 |
|  | platelet | 143 | 100 | 124 | 100 | 100 | 72 | 75.3 | 72.0 |

**TABLE S18.** Numbers of *KnockoffZoom* discoveries at low resolution (0.226 Mb) using different random seeds. These discoveries are stratified by their estimated local FDP.

| Phenotype | Estimated local FDP $\leq 0.1$ | | | | Estimated local FDP $> 0.1$ | | | |
| --- | --- | --- | --- | --- | --- | --- | --- | --- |
|  | Seed 1 | Seed 2 | Both | Stable (%) | Seed 1 | Seed 2 | Both | Stable (%) |
| height | 1886 | 1870 | 1752 | 93.3 | 1398 | 1285 | 1040 | 77.7 |
| bmi | 1080 | 1091 | 984 | 90.7 | 724 | 750 | 524 | 71.1 |
| platelet | 932 | 927 | 836 | 89.9 | 528 | 497 | 360 | 70.3 |
| sbp | 456 | 490 | 421 | 89.1 | 266 | 312 | 195 | 67.9 |
| cvd | 356 | 332 | 289 | 84.1 | 158 | 140 | 90 | 60.6 |
| hypothyroidism | 162 | 158 | 143 | 89.4 | 50 | 28 | 24 | 66.9 |
| respiratory | 160 | 154 | 129 | 82.2 | 16 | 4 | 2 | 31.2 |

**TABLE S19.** Numbers of *KnockoffZoom* discoveries at intermediate resolution (0.088 Mb) using different random seeds. Other details as in Table S18.

| Phenotype | Estimated local FDP $\leq 0.1$ | | | | Estimated local FDP $> 0.1$ | | | |
| --- | --- | --- | --- | --- | --- | --- | --- | --- |
|  | Seed 1 | Seed 2 | Both | Stable (%) | Seed 1 | Seed 2 | Both | Stable (%) |
| height | 1033 | 988 | 907 | 89.8 | 943 | 815 | 670 | 76.6 |
| bmi | 335 | 432 | 289 | 76.6 | 220 | 238 | 158 | 69.1 |
| platelet | 519 | 537 | 469 | 88.9 | 372 | 438 | 286 | 71.1 |
| sbp | 220 | 240 | 184 | 80.2 | 77 | 120 | 50 | 53.3 |
| cvd | 174 | 200 | 140 | 75.2 | 8 | 27 | 7 | 56.7 |

**TABLE S20.**
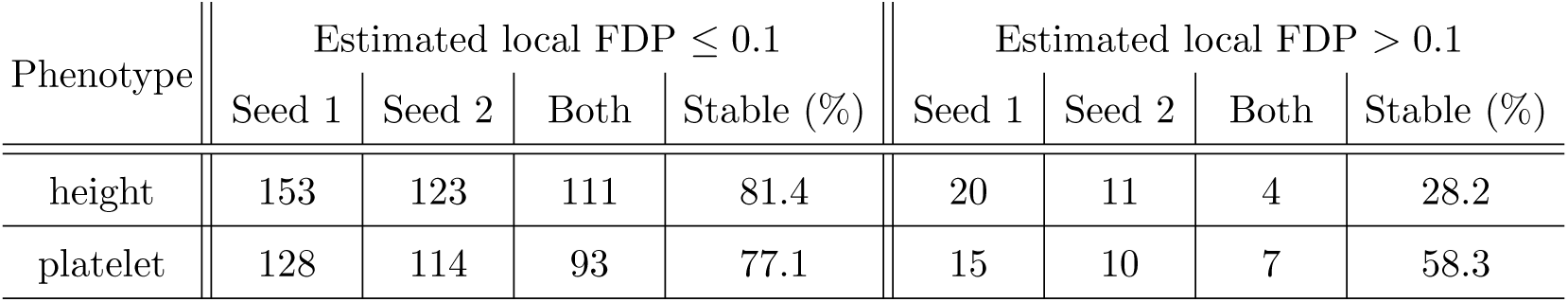
Numbers of *KnockoffZoom* discoveries at high resolution (single SNP) using different random seeds. Other details as in Table S18.

| Phenotype | Estimated local FDP $\leq 0.1$ | | | | Estimated local FDP $> 0.1$ | | | |
| --- | --- | --- | --- | --- | --- | --- | --- | --- |
|  | Seed 1 | Seed 2 | Both | Stable (%) | Seed 1 | Seed 2 | Both | Stable (%) |
| height | 153 | 123 | 111 | 81.4 | 20 | 11 | 4 | 28.2 |
| platelet | 128 | 114 | 93 | 77.1 | 15 | 10 | 7 | 58.3 |

**TABLE S21.** Variability in the numbers of *KnockoffZoom* discoveries for the phenotype *height* at low resolution (0.226 Mb) using different random seeds, as a function of the estimated local FDP. For example, 96.7% of the discoveries obtained with the first seed whose estimated local FDP is below 0.01 are also found using the second random seed.

| FDP | Seed 1 |  |  | Seed 2 |  |  |
| --- | --- | --- | --- | --- | --- | --- |
|  | Discoveries | Confirmed | Stable (%) | Discoveries | Confirmed | Stable (%) |
| [0,0.01) | 553 | 529 | 95.7 | 679 | 644 | 94.8 |
| [0.01,0.02) | 343 | 323 | 94.2 | 159 | 153 | 96.2 |
| [0.02,0.05) | 554 | 515 | 93.0 | 713 | 667 | 93.5 |
| [0.05,0.1) | 496 | 445 | 89.7 | 359 | 328 | 91.4 |
| [0.1,0.2) | 753 | 622 | 82.6 | 647 | 561 | 86.7 |
| [0.2,0.5) | 585 | 358 | 61.2 | 598 | 439 | 73.4 |

**TABLE S22.** Numbers of discoveries made by BOLT-LMM (459k European individuals) that are reproduced by *KnockoffZoom* (350k unrelated British individuals) at low resolution (0.226 Mb), using different random seeds for the generation of knockoffs. For example, 2431 out of 2464 discoveries reported by BOLT-LMM for *height* are consistently found by *KnockoffZoom* with both random seeds. The *KnockoffZoom* results obtained with the first random seed correspond to those in Table S8.

| Phenotype | BOLT-LMM<br>($5 \times 10^{-8}$) | *KnockoffZoom* | | |
| --- | --- | --- | --- | --- |
|  |  | Seed 1 | Seed 2 | Both seeds |
| height | 2464 | 2431 | 2434 | 2431 (98.7%) |
| bmi | 697 | 672 | 672 | 672 (96.4%) |
| platelet | 1204 | 1155 | 1154 | 1154 (95.8%) |
| sbp | 568 | 452 | 482 | 452 (79.6%) |
| cvd | 257 | 229 | 227 | 227 (88.3%) |
| hypothyroidism | 143 | 134 | 133 | 133 (93.0%) |

### S6. DISCUSSION

#### A. Missing and imputed variants

Section II C indicates that we have not included imputed variants in our analysis because they do not contain any information on the trait of interest, in addition to that carried by the genotyped variants. Since this statement appears to clash with common practice, we discuss it here.

Note that imputed genotypes are not directly observed; instead, their values are set on the basis of the observed genotypes and models for linkage disequilibrium derived from other datasets. The imputation process may not be deterministic—there is some randomness in the reconstruction of the imputed genotypes—but this extra stochasticity is independent of the phenotype and hence uninformative. Even if the imputation is very accurate, the following fundamental reality does not change: the imputed genotypes are not observed, they are reconstructed entirely on the basis of the observed ones. Therefore, imputed genotypes cannot provide meaningful additional information to that already carried by the genotyped variants. Why then has the field found them useful?

Imputed genotypes may help can increase the power to detect associated loci, as a result of the mismatch between the models used to analyze the relation between genotypes and phenotypes (typically linear) and the true biological mechanism (unknown). To illustrate, consider an example in the context of the standard analysis of GWAS data, where a univariate test probes for *marginal* association between a phenotype and each variant. Consider the case where an untyped variant *B* can be imputed with high accuracy, using the genotypes of two neighboring variants *A* and *C* (in the interest of concreteness, let’s say that the minor allele of *B* is usually observed only in conjunction with a specific haplotype of *A* and *C*). It is quite possible that the association test between the imputed variant *B* and the phenotype rightly results in a substantially smaller p-value than the association tests between the phenotype and either of the two variants *A* and *C* whose genotypes are used for imputation. This could happen because *B* is causal, and each of the genotyped variants only provides an imperfect proxy. Or it could happen because the phenotype depends on the two observed variants *A* and *C* in a nonlinear fashion: one needs the two alleles that define the haplotype corresponding to the minor allele of *B* to observe a change in the phenotype mean. The two situations are indistinguishable on the basis of data on *A* and *C* alone. However, this is not a problem if we simply want to identify the presence of an association. As long as we interpret a significant p-value as indicating the presence of an associated locus, without describing causal mechanisms, the increase in power is all that matters.

The situation changes substantially if we move the target from identifying a locus that contains causal variants to pinpointing the causal variants. In the first situation described above, *B* is causal, and changing the value of its allele would result in a change of the expected value of the phenotype. In the second situation, changing the major allele of *B* to its minor form, without touching the alleles at the neighboring variants *A* and *C*, will result in no change on the expected value of the phenotype. Distinguishing between these two situations on the basis of imputed data alone is not possible: any algorithm that would report {*B*} or {*A, C*} separately (as opposed to {*A, B, C*}) as containing the causal variants would do so either by relying on extra assumptions (e.g. causal sets contain the smallest possible number of variables or the true model is linear), or arbitrarily. Only genotyping *B*, and observing the few cases in which its minor allele occurs without the specific haplotype of {*A, C*} used for imputation, could allow one to choose between the two causal models. To put these issues in a broader perspective, identifying causal mechanisms without relying on experiments is notoriously difficult (recall the refrain “correlation is not causation”). Getting a handle on causal effects from large-scale observational data is one of the contemporary challenges in statistics and data science. From this perspective, the attempt to attribute causal effects to variables that are not even observed, let alone modified in an experiment, seems quite audacious. This underscores the danger of attributing causal effects to imputed variants.

The inferential framework of *KnockoffZoom* naturally steers the researchers away from making misleading causal statements on imputed variants. By construction, the imputed genotypes are independent of the phenotype after conditioning on the genotyped variants, and all the conditional hypotheses involving imputed variants are null. The situation is different for other fine-mapping tools (e.g., CAVIAR or SUSIE), even if these also investigate “conditional” hypotheses that probe the role of a variant given the other SNPs. These methods analyze several variants in the same locus using a Bayesian multivariate regression model in which the genetic design matrix *X* is considered fixed. Therefore, it is not meaningful to ask about conditional independence between the trait *Y* and *X*, since the latter is not random. Instead, in this context, “conditional” testing corresponds to asking which coefficients are nonzero in the assumed multivariate linear model for *Y*|*X*. Once *X* is considered fixed, there is no technical difference between a measured variant and an imputed one: the machinery will take them all as input and treat them equally.

Nevertheless, the same conceptual difficulties in establishing causal models on the basis of imputed variants discussed above still apply. On the one hand, if the imputation quality is low, one ends up analyzing variants that do not exist in reality, so the possible discoveries would not be scientifically relevant. On the other hand, high-quality imputation is only possible if the genotyped variants explain almost all of the variation in the missing variants. If this dependence is linear, the genotyped and imputed variants may be highly collinear and testing in the multivariate linear model powerless. This lack of power may be mitigated if the imputation process is highly nonlinear, since linear models are only sensitive to linear dependencies among variables. However, trusting the scientific validity of these discoveries would require one to place a lot of faith in the correctness of the linear model, as explained above.

While the conditional testing framework of *KnockoffZoom* intrinsically acknowledges the impossibility of differentiating between the effect of a missing variant from that of the genotyped ones used to impute it, two observations that have motivated practitioners to look at imputed variants continue to remain valid: (1) not all genetic variation is genotyped and therefore we need to consider the possibility that causal variants are untyped; (2) imputed variants can be utilized to construct more powerful tests. We now discuss how we can leverage these in our framework.

First, we have been careful in defining and interpreting our hypotheses so as to be meaningful in a context where not all variants are typed (1). This is why we choose to test spatially contiguous blocks of genotyped variants for association and report genomic intervals spanned by these blocks. This way, if *KnockoffZoom* returns a significant group of genotyped variants, we can be reasonably sure that either one of these variants is causal or there is at least one untyped causal variant in the corresponding genomic interval. While the *KnockoffZoom* methodology is agnostic to the choice of variable grouping and therefore can be applied to non-contiguous groups, such groups would be more difficult to interpret as long as there are untyped variants. For these reasons, we also find it most meaningful to interpret the “size” of a knockoff discovery as the width of the genomic interval it spans, rather than the number of genotyped variants it contains.

Second, while the resolution of the discoveries is fundamentally limited by the genotyped data, we remark that it is possible to leverage imputed variants with *KnockoffZoom* to increase power (2). Even if we are testing hypotheses defined only in terms of genotyped variants, we can use test statistics that capitalize on imputed ones, so as to most effectively capture the signal associated with a group of genotyped variants. We leave the implementation of such an approach for future work.

### S7. MATHEMATICAL PROOFS

#### Proof of Proposition 2.

Our algorithm implements the SCIP recipe,^15^ which provably generates valid knockoffs, for the vector-valued random variables *Z*^1^, …, *Z*^*L*^. Therefore, it suffices to show that (S3) gives a correct expression for 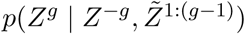. To this end, we proceed by induction. Assuming that (S3) holds ∀*g* ∈ {1, …, *g*′ − 1}, for some *g*′ > 1, it follows that:

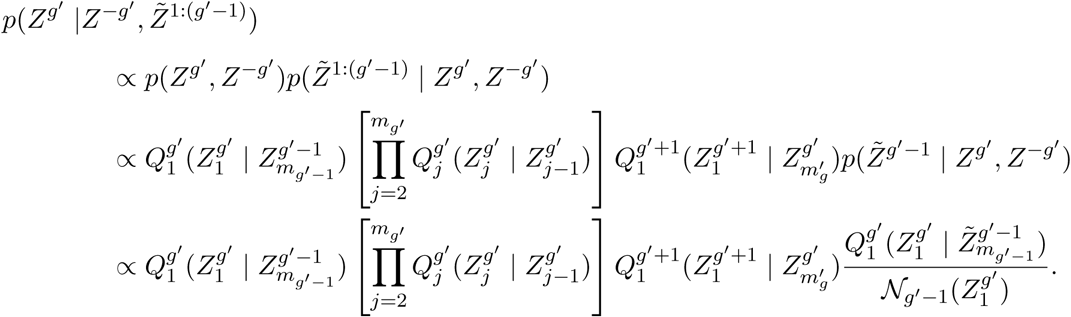

Above, the proportionality holds across values of the vector *Z*^*g*′^. In view of the normalization in (S4), we conclude that (S3) gives a correct expression for 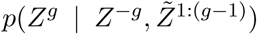 for all *g* ∈ {1, …, *g*′}. Since the base case with *g* = 1 clearly holds, the proof is complete. □

#### Proof of Lemma 1.

This follows immediately by replacing (S7) into (S8) and simplifying. □

#### Proof of Proposition 3.

It suffices to prove (S6), since marginalizing over 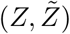 implies that 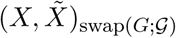 has the same distribution as 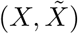.^2^ Conditioning on the latent variables yields:

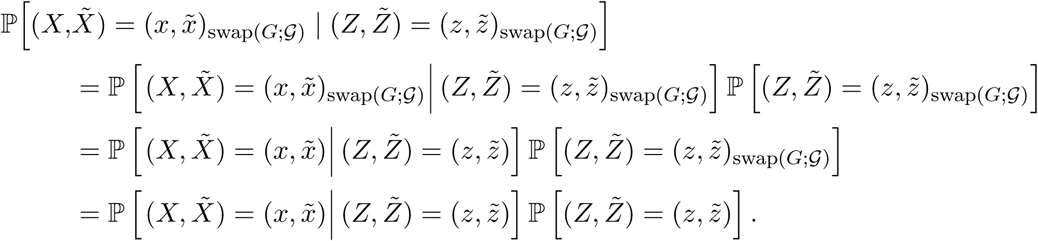

Above, the first equality follows from the first line of Algorithm 3, the second from the conditional independence of the emission distributions in an HMM, and the third from Proposition 2). □

#### Proof of Proposition 4.

The 𝒩 function for the *g*-th group can be written as:

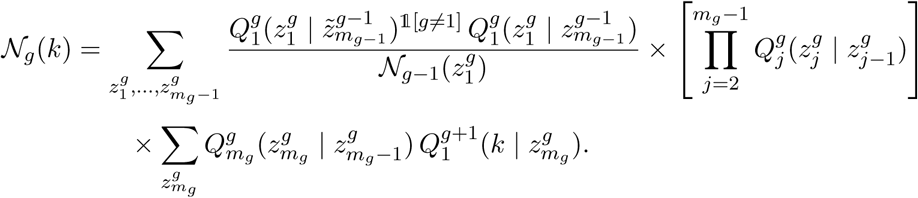

To simplify the notation, we define 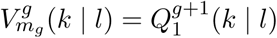, and, recursively for *j* ∈ {1, …, *m*_*g*_ −1},

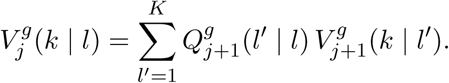

Then, we see that the 𝒩 function can be reduced to:

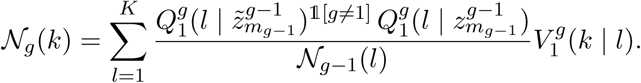

It remains to be shown how to compute 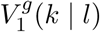. We guess that 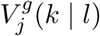 can be written as:

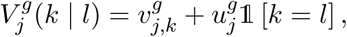

since 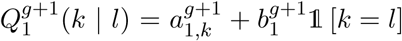. This ansatz is clearly correct if *j* = *m*_*g*_, by definition of 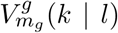, in which case: 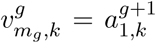 and 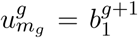. When *j* ∈ {1, …, *m*_*g*_ − 1}, backward induction shows that

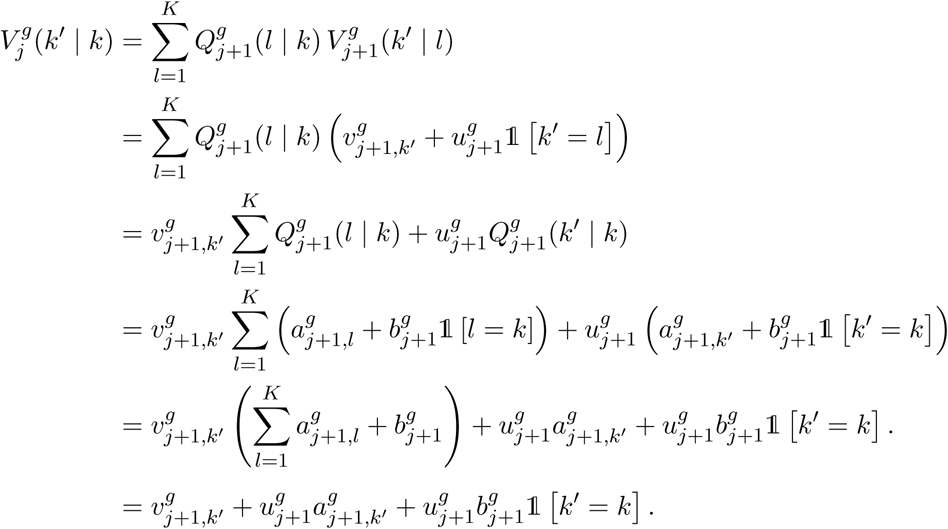

Therefore, our guess is correct for *j*, as long as it is for *j* + 1, since

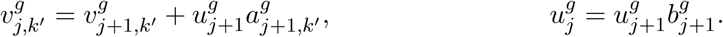

□

#### Proof of Proposition 5.

We start from

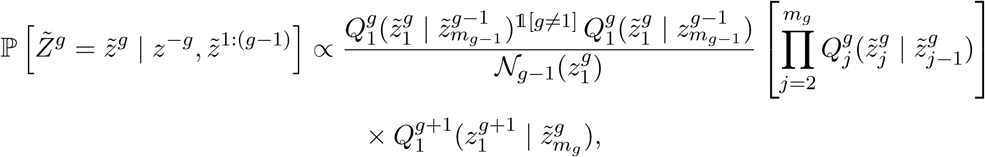

and marginalize over the remaining *m*_*g*_ − 1 components, finding:

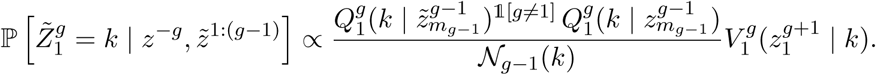

Above, 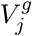 is as defined in the proof of Proposition 4. Given 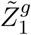, we can sample each 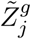 sequentially, from *j* = 2 to *j* = *m*_*g*_. It is easy to verify that, at the *j*-th step, we need to sample 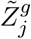 from:

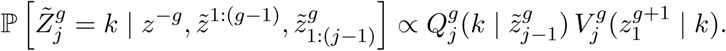

□

#### Proof of Proposition 6.

Starting from

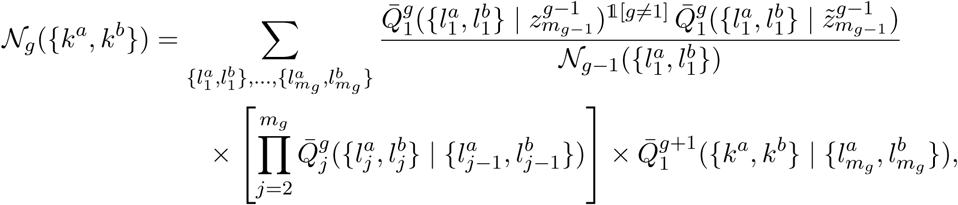

we proceed as in the proof of Proposition 4, defining:

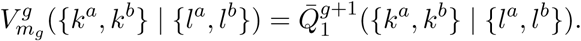

Similarly, for *j* = 1, …, *m*_*g*_ − 1, we recursively define:

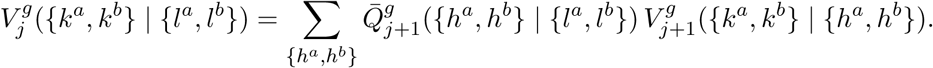

Using Lemma 1, we can write:

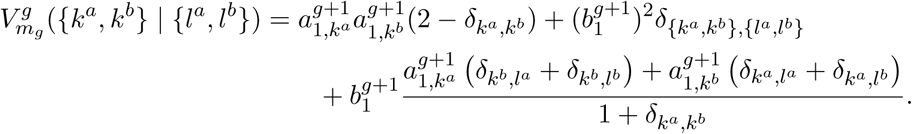

It is now time for an ansatz. For *j* = 1, …, *m*_*g*_ − 1, we guess that:

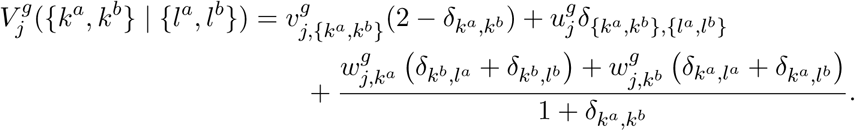

Assuming that our guess is correct for *j* + 1, with some *j* ∈ {1, …, *m*_*g*_ − 1}, we can write:

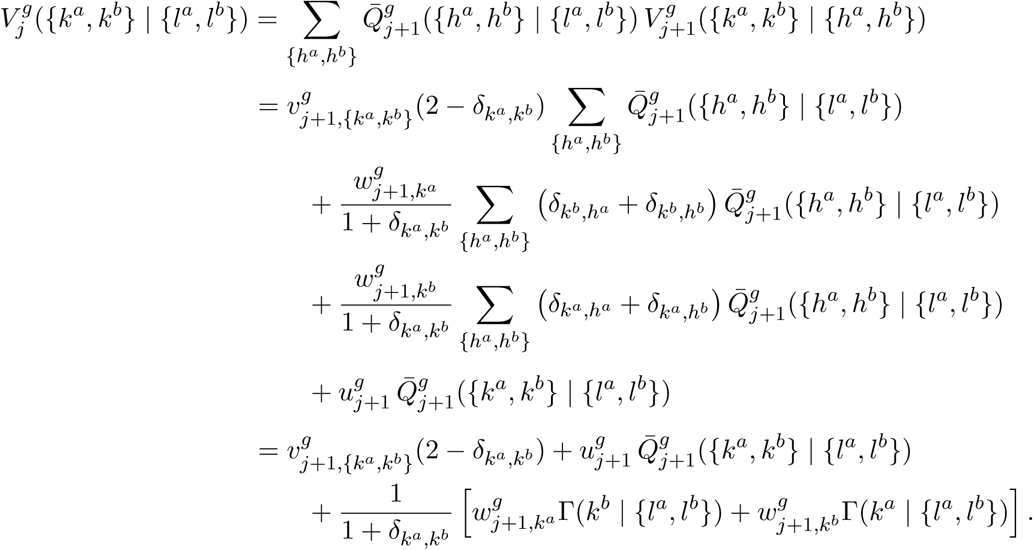

Above, we have defined:

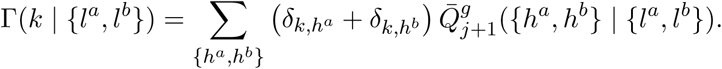

This can be further simplified:

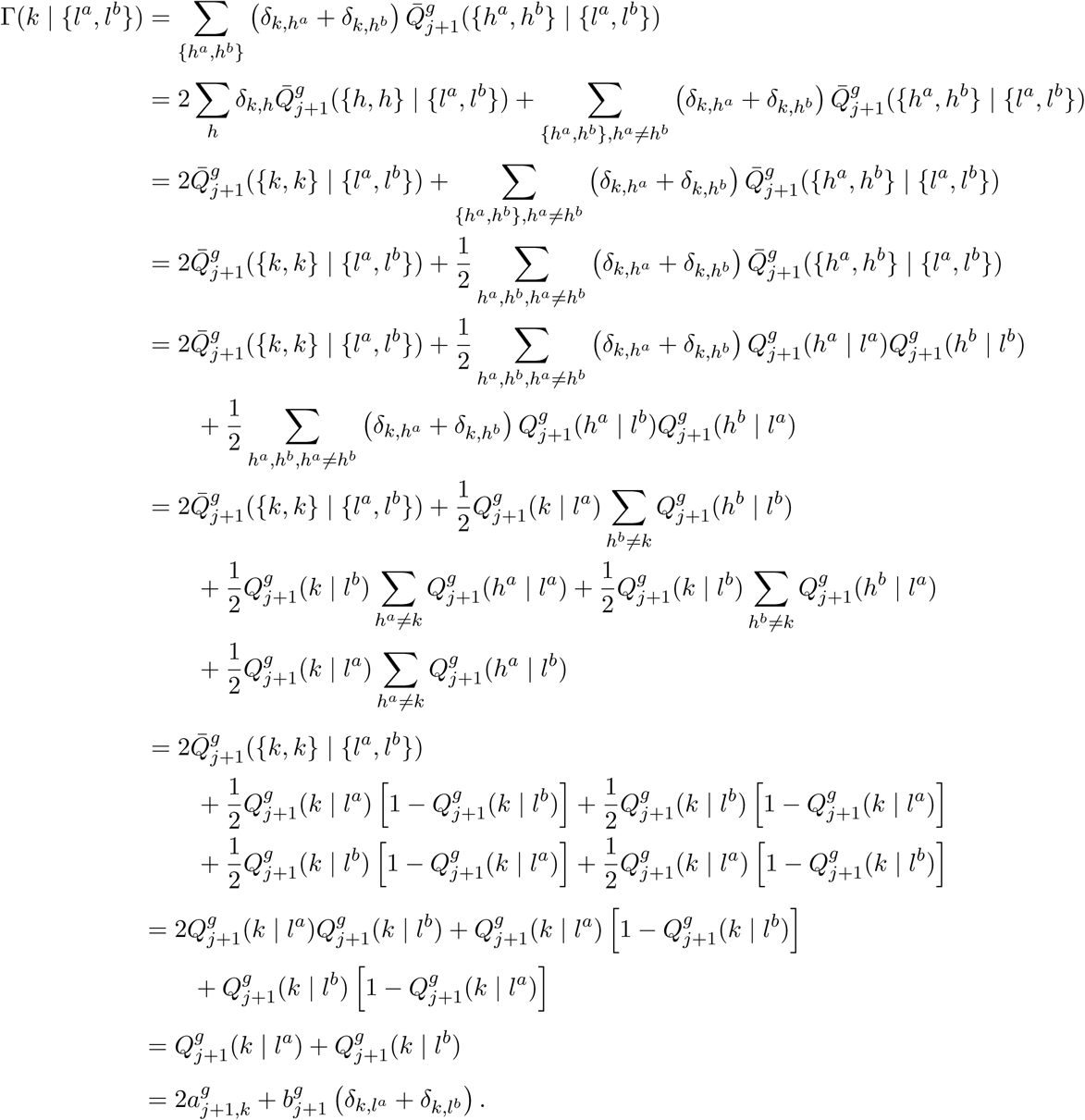

Therefore, we can rewrite 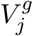, as follows:

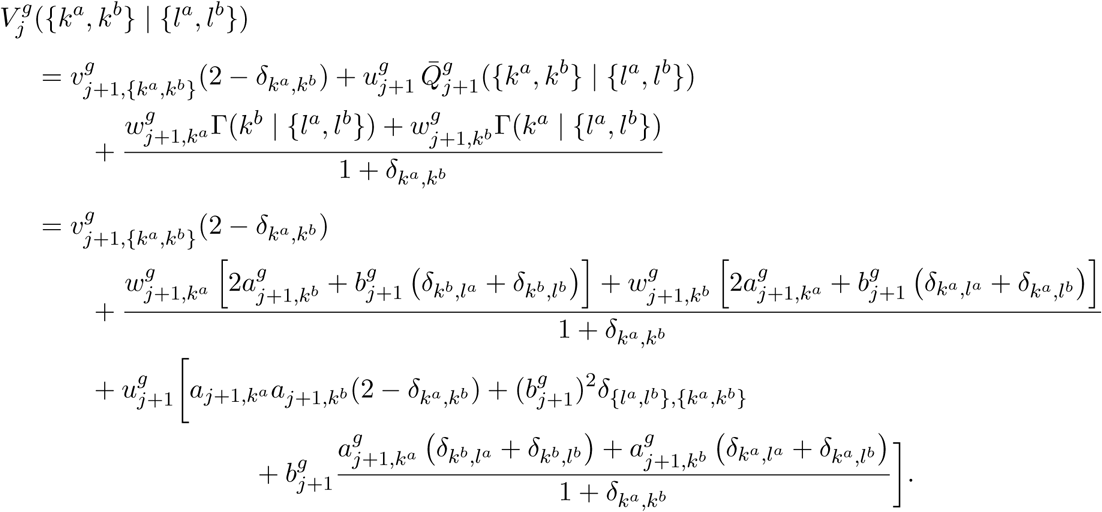

Now, we only need to collect these terms to obtain the recursion rules:

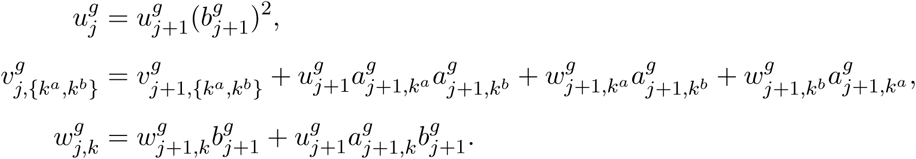

Finally, the 𝒩 function for the *g*-th group is given by:

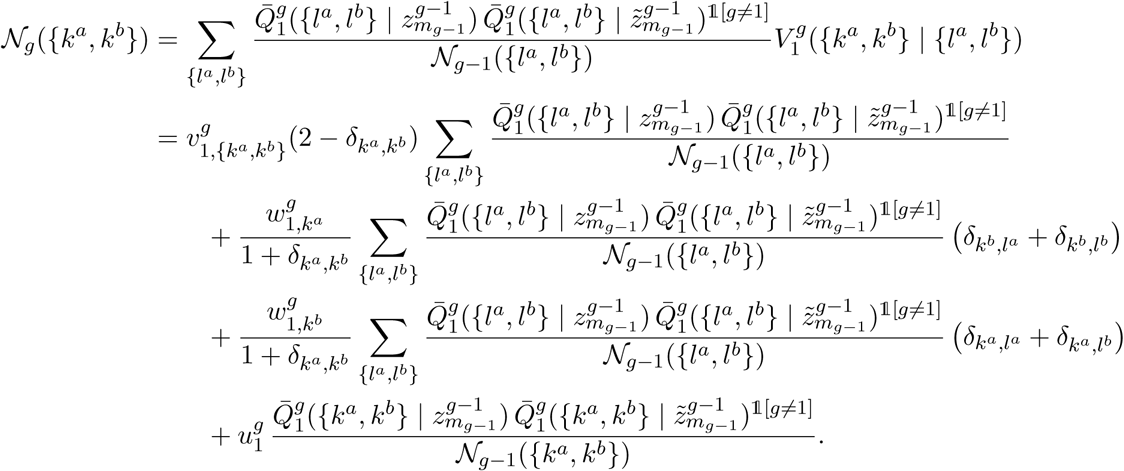

Let us now define

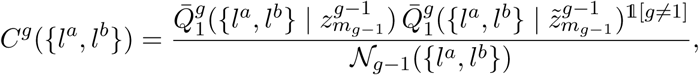

and

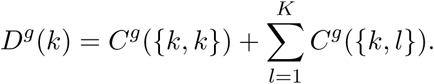

Then, we can write the 𝒩 function more compactly, as follows:

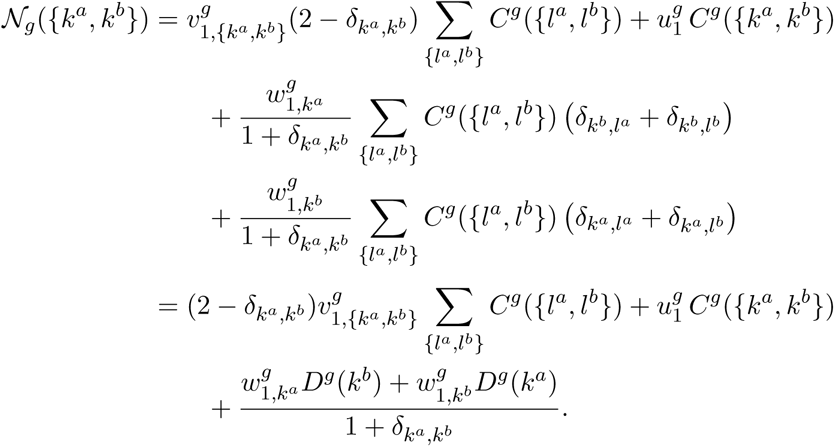

□

#### Proof of Proposition 7.

We proceed as in the proof of Proposition 5, using the notation developed in the proof of Proposition 6. By marginalizing over the remaining *m*_*g*_ − 1 components, it turns out that the marginal distribution of 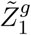 is:

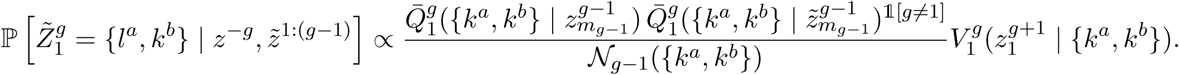

Since we have already obtained 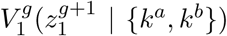, by computing the 𝒩 function, sampling 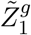 can be performed easily, in 𝒪(*K*^2^) time. Given 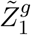, we proceed to sample each 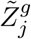 sequentially, from *j* = 2 to *j* = *m*_*g*_. It is easy to verify that, at the *j*-th step, we sample 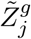 from:

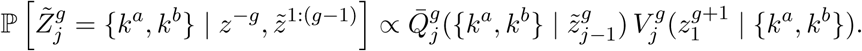

Again, this can be easily performed in 𝒪(*K*^2^) time, for each *j*. □

#### Proof of Proposition 8.

The result for *F*_1_(*k*) is immediate from (S7). For *j* ∈ {2, …, *p*},

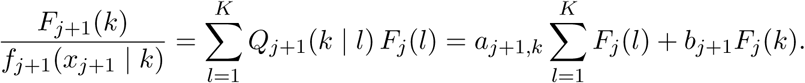

□

#### Proof of Proposition 9.

The case if *F*_1_({*k*^*a*^, *k*^*b*^}) follows directly from (S8). For *j* ∈ {2, …, *p*},

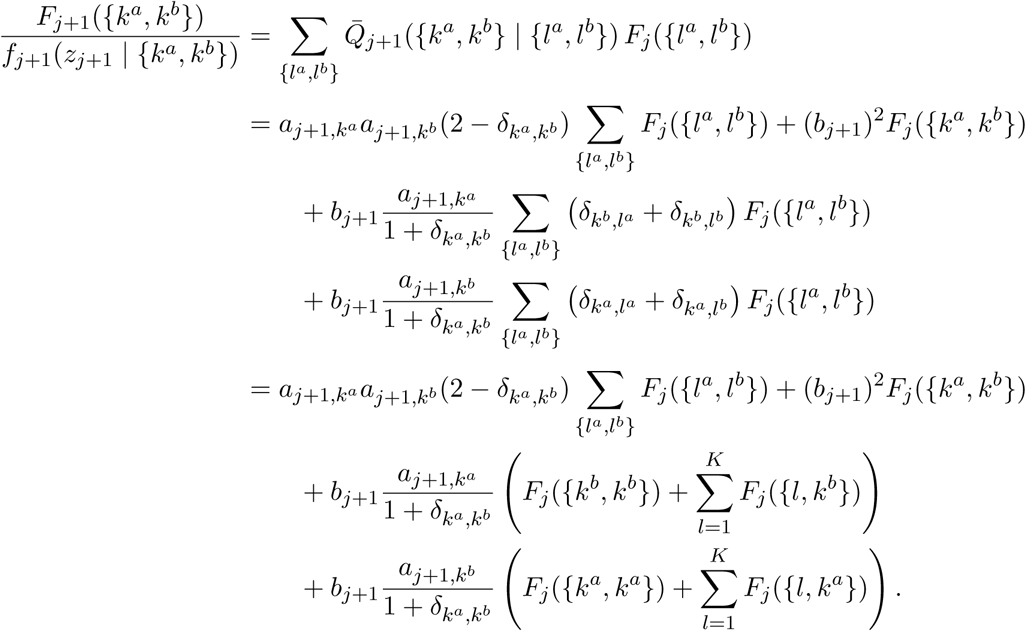

□

## References

1 Visscher, P. M. et al. 10 years of GWAS discovery: biology, function, and translation. Am. J. Hum. Genet. 101, 5–22 (2017).

2 Tam, V. et al. Benefits and limitations of genome-wide association studies. Nature Reviews Genetics (2019). URL https://doi.org/10.1038/s41576-019-0127-1.

3 Yu, J. et al. A unified mixed-model method for association mapping that accounts for multiple levels of relatedness. Nat. Genet. 38, 203–208 (2006).

4 Kang, H. M. et al. Efficient control of population structure in model organism association mapping. Genetics 178, 1709–1723 (2008).

5 Kang, H. M. et al. Variance component model to account for sample structure in genome-wide association studies. Nat. Genet. 42, 348–354 (2010).

6 Zhang, Z. et al. Mixed linear model approach adapted for genome-wide association studies. Nat. Genet. 42, 355–360 (2010).

7 Loh, P.-R. et al. Efficient Bayesian mixed-model analysis increases association power in large cohorts. Nat. Genet. 47, 284 (2015).

8 Loh, P.-R., Kichaev, G., Gazal, S., Schoech, A. P. & Price, A. L. Mixed-model association for biobank-scale datasets. Nat. Genet. 50, 906–908 (2018).

9 Reich, D. E. et al. Linkage disequilibrium in the human genome. Nature 411, 199–204 (2001).

10 Pritchard, J. K. & Przeworski, M. Linkage disequilibrium in humans: models and data. Am. J. Hum. Genet. 69, 1–14 (2001).

11 Boyle, E. A., Li, Y. I. & Pritchard, J. K. An expanded view of complex traits: from polygenic to omnigenic. Cell 169, 1177–1186 (2017).

12 Purcell, S. et al. PLINK: a tool set for whole-genome association and population-based linkage analyses. Am. J. Hum. Genet. 81, 559–575 (2007).

13 Schaid, D. J., Chen, W. & Larson, N. B. From genome-wide associations to candidate causal variants by statistical fine-mapping. Nat. Rev. Genet. 19, 491–504 (2018).

14 Hormozdiari, F., Kostem, E., Kang, E. Y., Pasaniuc, B. & Eskin, E. Identifying causal variants at loci with multiple signals of association. Genetics 198, 497–508 (2014).

15 Kichaev, G. et al. Integrating functional data to prioritize causal variants in statistical fine-mapping studies. PLoS Genet. 10, e1004722 (2014).

16 Benner, C. et al. FINEMAP: efficient variable selection using summary data from genome-wide association studies. Bioinformatics 32, 1493–1501 (2016).

17 Wang, G., Sarkar, A. K., Carbonetto, P. & Stephens, M. A simple new approach to variable selection in regression, with application to genetic fine-mapping. bioRxiv (2018).

18 Benjamini, Y. & Hochberg, Y. Controlling the false discovery rate: a practical and powerful approach to multiple testing. J. R. Stat. Soc. B. 57, 289–300 (1995).

19 Li, N. & Stephens, M. Modeling linkage disequilibrium and identifying recombination hotspots using single-nucleotide polymorphism data. Genetics 165, 2213–2233 (2003).

20 Scheet, P. & Stephens, M. A fast and flexible statistical model for large-scale population genotype data: applications to inferring missing genotypes and haplotypic phase. Am. J. Hum. Genet. 78, 629–644 (2006).

21 Marchini, J. & Howie, B. Genotype imputation for genome-wide association studies. Nat. Rev. Genet. 11, 499–511 (2010).

22 O’Connell, J. et al. Haplotype estimation for biobank scale datasets. Nat. Genet. 48, 817–820 (2016).

23 Candès, E. J., Fan, Y., Janson, L. & Lv, J. Panning for gold: Model-x knockoffs for high-dimensional controlled variable selection. J. R. Stat. Soc. B. 80, 551–577 (2018).

24 Sesia, M., Sabatti, C. & Candès, E. J. Gene hunting with hidden Markov model knockoffs. Biometrika 106, 1–18 (2019).

25 Bottolo, L. & Richardson, S. Discussion of gene hunting with hidden Markov model knockoffs. Biometrika 106, 19–22 (2019).

26 Jewell, S. W. & Witten, D. M. Discussion of gene hunting with hidden Markov model knockoffs. Biometrika 106, 23–26 (2019).

27 Rosenblatt, J. D., Ritov, Y. & Goeman, J. J. Discussion of gene hunting with hidden Markov model knockoffs. Biometrika 106, 29–33 (2019).

28 Marchini, J. L. Discussion of gene hunting with hidden Markov model knockoffs. Biometrika 106, 27–28 (2019).

29 Sesia, M., Sabatti, C. & Candès, E. J. Rejoinder: Gene hunting with hidden Markov model knockoffs. Biometrika 106, 35–45 (2019).

30 Bycroft, C. et al. The UK biobank resource with deep phenotyping and genomic data. Nature 562, 203–209 (2018).

31 Sabatti, C. Multivariate Linear Models for GWAS, 188–207 (Cambridge University Press, 2013).

32 Davidson, I. Agglomerative hierarchical clustering with constraints: Theoretical and empirical results. European Conference on Principles of Data Mining and Knowledge Discovery 59–70 (2005).

33 Weller, J. I., Song, J. Z., Heyen, D. W., Lewin, H. A. & Ron, M. A new approach to the problem of multiple comparisons in the genetic dissection of complex traits. Genetics 150, 1699–1706 (1998).

34 Sabatti, C., Service, S. & Freimer, N. False discovery rate in linkage and association genome screens for complex disorders. Genetics 164, 829–833 (2003).

35 Brzyski, D. et al. Controlling the rate of GWAS false discoveries. Genetics 205, 61–75 (2017).

36 Barber, R. F. & Candès, E. J. Controlling the false discovery rate via knockoffs. Ann. Stat. 43, 2055–2085 (2015).

37 Dai, R. & Barber, R. F. The knockoff filter for FDR control in group-sparse and multitask regression. J. Mach. Learn. Res. 48, 1851–1859 (2016).

38 Privé, F., Aschard, H., Ziyatdinov, A. & Blum, M. G. B. Efficient analysis of large-scale genome-wide data with two R, packages: bigstatsr and bigsnpr. Bioinformatics 34, 2781–2787 (2018).

39 Katsevich, E. & Sabatti, C. Multilayer knockoff filter: controlled variable selection at multiple resolutions. Ann. Appl. Stat. 13, 1–33 (2019).

40 Efron, B. Large-Scale Inference: Empirical Bayes Methods for Estimation, Testing, and Prediction (Cambridge University Press, 2010).

41 Price, A. L. et al. Principal components analysis corrects for stratification in genome-wide association studies. Nat. Genet. 38, 904–909 (2006).

42 Klasen, J. R. et al. A multi-marker association method for genome-wide association studies without the need for population structure correction. Nat. Commun. 7 (2016).

43 Katsevich, E., Sabatti, C. & Bogomolov, M. Controlling FDR while highlighting distinct discoveries. arXiv preprint 1809.01792 (2018).

44 Ernst, J. & Kellis, M. Discovery and characterization of chromatin states for systematic annotation of the human genome. Nat. Biotechnol. 28, 817–825 (2010).

45 Pruim, R. J. et al. LocusZoom: regional visualization of genome-wide association scan results. Bioinfor-matics 26, 2336–2337 (2010).

46 McLean, C. Y. et al. GREAT improves functional interpretation of cis-regulatory regions. Nature biotechnology 28, 495 (2010).

47 Hoggart, C. J., Whittaker, J. C., De Iorio, M. & Balding, D. J. Simultaneous analysis of all SNPs in genome-wide and re-sequencing association studies. PLoS genetics 4, e1000130 (2008).

48 Guan, Y. & Stephens, M. Bayesian variable selection regression for genome-wide association studies and other large-scale problems. The Annals of Applied Statistics 5, 1780–1815 (2011).

49 Buzdugan, L. et al. Assessing statistical significance in multivariable genome wide association analysis. Bioinformatics 32, 1990–2000 (2016).

50 Renaux, C., Buzdugan, L., Kalisch, M. & Bühlmann, P. Hierarchical inference for genome-wide association studies: a view on methodology with software. arXiv preprint 1805.02988 (2018).

51 Zhou, W. et al. Efficiently controlling for case-control imbalance and sample relatedness in large-scale genetic association studies. Nat. Genet. 50, 1335–1341 (2018).

52 Wu, T. T., Chen, Y. F., Hastie, T., Sobel, E. & Lange, K. Genome-wide association analysis by lasso penalized logistic regression. Bioinformatics 25, 714–721 (2009).

53 Wu, J., Devlin, B., Ringquist, S., Trucco, M. & Roeder, K. Screen and clean: a tool for identifying interactions in genome-wide association studies. Genet Epidemiol. 34, 275–285 (2010).

## References

1 Katsevich, E. & Sabatti, C. Multilayer knockoff filter: controlled variable selection at multiple resolutions. Ann. Appl. Stat. 13, 1–33 (2019).

2 Sesia, M., Sabatti, C. & Candès, E. J. Gene hunting with hidden Markov model knockoffs. Biometrika 106, 1–18 (2019).

3 Scheet, P. & Stephens, M. A fast and flexible statistical model for large-scale population genotype data: applications to inferring missing genotypes and haplotypic phase. Am. J. Hum. Genet. 78, 629–644 (2006).

4 Marchini, J. & Howie, B. Genotype imputation for genome-wide association studies. Nat. Rev. Genet. 11, 499–511 (2010).

5 O’Connell, J. et al. Haplotype estimation for biobank scale datasets. Nat. Genet. 48, 817–820 (2016).

6 Fearnhead, P. & Donnelly, P. Estimating recombination rates from population genetic data. Genetics 159, 1299–1318 (2001).

7 Privé, F., Aschard, H., Ziyatdinov, A. & Blum, M. G. B. Efficient analysis of large-scale genome-wide data with two R, packages: bigstatsr and bigsnpr. Bioinformatics 34, 2781–2787 (2018).

8 Abraham, G., Kowalczyk, A., Zobel, J. & Inouye, M. Performance and robustness of penalized and unpenalized methods for genetic prediction of complex human disease. Genet. Epidemiol. 37, 184–195 (2013).

9 Vilhjá, B. J. lmsson et al. Modeling linkage disequilibrium increases accuracy of polygenic risk scores. Am. J. Hum. Genet. 97, 576–592 (2015).

10 Wei, Z. et al. Large sample size, wide variant spectrum, and advanced machine-learning technique boost risk prediction for inflammatory bowel disease. Am. J. Hum. Genet. 92, 1008–1012 (2013).

11 Botta, V., Louppe, G., Geurts, P. & Wehenkel, L. Exploiting SNP correlations within random forest for genome-wide association studies. PLoS One 9 (2014).

12 Bellot, P. & Pérez-Enciso, M. Can deep learning improve genomic prediction of complex human traits? Genetics 210, 809–819 (2018).

13 Klasen, J. R. et al. A multi-marker association method for genome-wide association studies without the need for population structure correction. Nat. Commun. 7 (2016).

14 Zeng, Y. & Breheny, P. The biglasso package: A memory-and computation-efficient solver for lasso model fitting with big data in R. arXiv (2017).

15 Candès, E. J., Fan, Y., Janson, L. & Lv, J. Panning for gold: Model-x knockoffs for high-dimensional controlled variable selection. J. R. Stat. Soc. B. 80, 551–577 (2018).

16 Benjamini, Y. & Hochberg, Y. Controlling the false discovery rate: a practical and powerful approach to multiple testing. J. R. Stat. Soc. B. 57, 289–300 (1995).

17 Simes, R. J. An improved Bonferroni procedure for multiple tests of significance. Biometrika 73, 751–754 (1986).

18 Katsevich, E., Sabatti, C. & Bogomolov, M. Controlling FDR while highlighting distinct discoveries. arXiv preprint 1809.01792 (2018).

19 Siegmund, D. O., Zhang, N. R. & Yakir, B. False discovery rate for scanning statistics. Biometrika 98, 979–985 (2011).

20 Brzyski, D. et al. Controlling the rate of GWAS false discoveries. Genetics 205, 61–75 (2017).

21 Benner, C. et al. FINEMAP: efficient variable selection using summary data from genome-wide association studies. Bioinformatics 32, 1493–1501 (2016).

22 Kichaev, G. et al. Integrating functional data to prioritize causal variants in statistical fine-mapping studies. PLoS Genet. 10, e1004722 (2014).

23 Schaid, D. J., Chen, W. & Larson, N. B. From genome-wide associations to candidate causal variants by statistical fine-mapping. Nature Reviews Genetics 19, 491–504 (2018).

24 Wang, G., Sarkar, A. K., Carbonetto, P. & Stephens, M. A simple new approach to variable selection in regression, with application to genetic fine-mapping. bioRxiv (2018).

25 Hormozdiari, F., Kostem, E., Kang, E. Y., Pasaniuc, B. & Eskin, E. Identifying causal variants at loci with multiple signals of association. Genetics 198, 497–508 (2014).

26 Barber, R. F. & Candès, E. J. Controlling the false discovery rate via knockoffs. Ann. Stat. 43, 2055–2085 (2015).

27 Efron, B. Large-Scale Inference: Empirical Bayes Methods for Estimation, Testing, and Prediction (Cambridge University Press, 2010).

28 Loh, P.-R., Kichaev, G., Gazal, S., Schoech, A. P. & Price, A. L. Mixed-model association for biobank-scale datasets. Nat. Genet. 50, 906–908 (2018).

29 McLean, C. Y. et al. GREAT improves functional interpretation of cis-regulatory regions. Nature biotechnology 28, 495 (2010).

30 Smigielski, E. M., Sirotkin, K., Ward, M. & Sherry, S. T. dbSNP: a database of single nucleotide polymorphisms. Nucleic acids research 28, 352–355 (2000).

